# Extreme Hydrophobicity of Cytotoxic Drugs Enables Design of Next Generation Antibody-Drug Conjugates Nanotherapeutics

**DOI:** 10.64898/2026.04.29.721383

**Authors:** Ashwini R. Khyade, Abhishek Sharma, Britto S. Sandanaraj

## Abstract

Antibody and protein-drug conjugates (XDCs) have emerged as promising cancer therapeutics, yet their clinical utility remains constrained by dose-limiting toxicities and narrow therapeutic windows. These safety challenges stem primarily from two factors: premature payload release during systemic circulation, and poor physicochemical properties inherent to the hydrophobic cytotoxic drugs they carry. Prior strategies attempted to address these limitations by appending water-soluble tags to reduce overall conjugate hydrophobicity, but achieved only modest improvements. As a result, the hydrophobic nature of cytotoxic payloads has remained a persistent obstacle in XDC development. Here, we report a fundamentally different chemical strategy that reframes this liability as a design opportunity. Rather than masking drug hydrophobicity, we exploit it as the driving force for self-assembly of facially amphiphilic protein-drug conjugates with programmable drug moieties (PDCs). In this architecture, the hydrophobic cytotoxic drug and the hydrophilic protein serve as the core and shell, respectively, spontaneously assembling into monodisperse, well-defined spherical protein nanotherapeutics of controlled size. This design principle transforms a longstanding physicochemical challenge into a functional engineering tool, enabling precise nanostructure formation without sacrificing potency. In vitro studies confirm that the resulting nanotherapeutics effectively kill cancer cells, establishing a strong foundation for further therapeutic development.

## Introduction

Antibody/protein–drug conjugates (XDCs) have emerged as a preferred therapeutic modality for the treatment of various diseases, including cancer.^1–5^ To date, the Food and Drug Administration (FDA) has approved 15 ADCs, and over 1000 are currently under evaluation at various stages of clinical and preclinical development.^6–12^ ADCs comprise three core structural components: (i) an antibody, (ii) a cytotoxic drug (payload), and (iii) a linker that covalently joins the antibody to the drug.^13–17^ADCs with high drug-to-antibody ratios (DAR ≥ 8) have demonstrated superior efficacy; however, achieving such high DARs is technically challenging for several reasons.^17–19^ First, most cytotoxic payloads are highly hydrophobic, requiring the use of organic co-solvents, such as DMSO, during bioconjugation reaction. Second, hydrophobic ADCs are prone to non-specific aggregation in aqueous media driven by hydrophobic interactions.^20–22^ Third, and most critically, high-DAR ADCs often exhibit poor in vivo pharmacokinetic (PK), safety, and efficacy profiles.^15,17,22^

As a consequence, the hydrophobicity of cytotoxic payloads has been widely regarded as a significant liability in ADC design.^20^ Several strategies have been explored to mitigate this issue, including the use of inherently hydrophilic drugs.^21^ However, such compounds often exhibit poor membrane permeability and lack bystander activity, resulting in diminished potency. ^23^ The most widely adopted approach has been to incorporate hydrophilic polymers or solubility-enhancing tags into the linker.^24–26^ While these modifications can improve certain properties of the ADC, they do not comprehensively resolve all hydrophobicity-related challenges.^27^ To our knowledge, all prior strategies have focused on reducing the overall hydrophobicity of ADCs by covalently attaching exogenous hydrophilic polymers or tags.^23,25–28^

In parallel, our laboratory has pioneered a suite of chemical technologies for the synthesis of facially amphiphilic proteins (FAPs).^29–36^ The FAPs has three core structural elements: (i) a globular protein scaffold, (ii) a hydrophilic linker, and (iii) a hydrophobic tail - a structural architecture that closely resembles that of ADCs. In prior studies, we have demonstrated that FAPs self-assemble into precise, spherical protein nanoparticles (SPNPs) of defined size, molecular weight, and oligomeric state, with self-assembly driven exclusively by hydrophobic interactions mediated by lipid-like moieties.^29–36^

These observations prompted a fundamental question: could a hydrophobic cytotoxic drug replace the hydrophobic tail of FAPs, and could the inherent hydrophobicity of the drug be harnessed to drive self-assembly of FAPs into engineered SPNPs? Such a design would not only mask the payload’s hydrophobicity but also protect the drug by sequestering it within the nanoparticle core.^29,31,32^ Herein, we report the design and synthesis of facially amphiphilic protein–drug conjugates and demonstrate that these conjugates self-assemble into SPNPs of defined size and capable of killing cancer cells.

## Results

### Selection of Protein, Drug, and Linker

Human serum albumin (HSA) was selected as the model protein scaffold for the following reasons: (i) it is a well-characterized globular protein with a molecular weight of 66 kDa; (ii) bioconjugation can be performed site-specifically via its single free cysteine residue (Cys34); ^37,38^ and (iii) HSA is biocompatible, biodegradable, and non-immunogenic. Importantly, HSA has been incorporated as an active ingredient in several clinically approved nanomedicines.^39–49^ Camptothecin (CT) was chosen as the cytotoxic payload because: (i) it is a widely used drug in clinically approved ADCs; (ii) it possesses a more favorable safety profile relative to other cytotoxic agents; and (iii) it contains a functional handle amenable to chemical derivatization.^50–55^ Oligoethylene glycol (OEG) linkers of defined chain length were employed owing to their conformational flexibility and biocompatibility. OEG-based linkers are also widely used in clinically approved ADCs and nanomedicine formulations.^56^

### Synthesis of Azide Functionalized Drug Conjugates and HSA-COT conjugate

To establish a modular conjugation strategy, we designed a hydrophilic bi-functional linker containing a maleimide (MI) group for site-selective cysteine conjugation and a cyclooctyne (COT) moiety for copper-free strain-promoted azide–alkyne cycloaddition (SPAAC).^31,57^An octaethylene glycol (OEG) spacer was incorporated to enhance aqueous solubility and minimize aggregation during bioconjugation.^29,30,58^ The MI–OEG–COT linker was synthesized in five steps, yielding 15% overall **(scheme 1a)**. Camptothecin (CPT) was selected as a model cytotoxic payload. To enhance its hydrophobic character and promote post-conjugation self-assembly, we designed a modified CPT incorporating (i) a decyl spacer to increase hydrophobicity, (ii) an acid-labile carbonate linkage for pH-triggered intracellular release ^59–61^, and (iii) a terminal azide for orthogonal click conjugation. The CPT intermediate (3d) was synthesized in four steps from CPT (36% overall yield) and confirmed by NMR and MALDI-TOF-MS **(scheme 1b)**. To control drug valency, mono-, di-, and tri-alkyne OEG modules (1T/2T/3T) were prepared in four steps (overall yields: 45%, 14%, and 16%, respectively). The corresponding CPT azide tails were obtained *via* CuAAC coupling with the hydrophobic CPT intermediate, followed by terminal azidation to regenerate SPAAC reactivity. The final 1T/2T/3T constructs required 10 total steps from CPT, with overall yields of 5%, 2%, and 3%, respectively **(scheme 1c-d)**. Despite modest cumulative yields, this modular approach enables precise control over drug valency, hydrophobicity, and release functionality. All final compounds were purified by normal-phase chromatography and fully characterized by ^1^H, ^13^C NMR and MALDI-TOF MS **(see Supporting Information)**. The bioconjugation of HSA with the compound MI-OEG-COT (1e), featuring a maleimide–cyclooctyne functionality, led to effective thiol–maleimide conjugation at Cys34, resulting in the formation of the HSA-MI-OEG-COT conjugate with complete conversion **(scheme 2a**), as validated by MALDI-TOF-MS revealing a single peak at 67193 Da **(figure 1a)**. Because of the near-complete conversion, the crude reaction mixture was used directly in the subsequent SPAAC reactions without further purification. The comprehensive bioconjugation procedure is detailed in the methods section.

**Figure 1:**
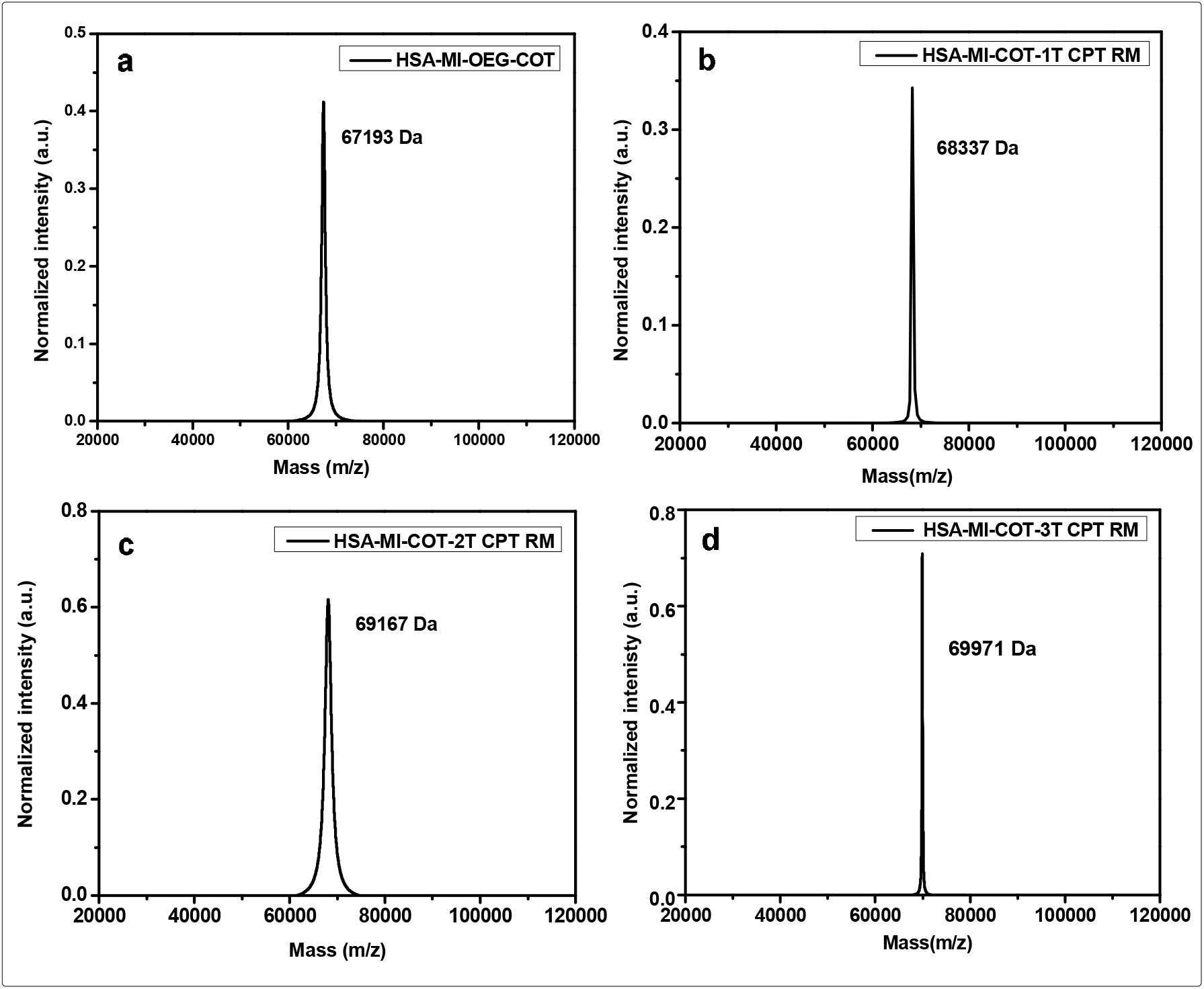
MALDI-TOF MS analysis of HSA conjugation and post-click modification. a. MALDI-TOF MS spectrum of RM after 16 h shows a single peak at 67,193 Da, confirming successful HSA conjugation. b. MALDI-TOF MS of the post-click reaction displays an increased molecular weight, indicating quantitative conjugation of the CPT-azide tail to HSA–MI–OEG–COT.

### SPAAC Conjugation via Supramolecular Assisted Protein Labeling Technology (SAPLabTech)

For the strain-promoted azide–alkyne cycloaddition (SPAAC) involving HSA–MI–OEG–COT and the 1T/2T/3T CPT azide tails, a significant challenge was the limited solubility of the CPT azide derivatives in water due to their high hydrophobic nature, especially with the more complex drug-valency constructs. To overcome this issue, we utilized SAPLabTech (Supramolecular Assisted Protein Labeling Technology), developed in our lab, to facilitate efficient bioconjugation in entirely aqueous conditions. At first, we assessed the solubility of the 1T CPT azide in the presence of γ-cyclodextrin. Fortunately, γ-cyclodextrin significantly improved the solubility of the hydrophobic CPT derivative in water, through host–guest inclusion complexes. The solubilized 1T CPT azide was then combined with the crude reaction mixture (RM) containing HSA–MI–OEG–COT and incubated at room temperature while gently mixing with a rotaspin **(scheme 2a)**. After 24 hours, complete conversion to the intended conjugate was achieved. MALDI-TOF mass spectrometry analysis of the reaction mixture showed a single peak at 68,337 Da, which corresponds to HSA–MI–OEG–COT–1T CPT, with no identifiable residual starting material **(Figure 1b)**. Encouraged by these results, the same supramolecular approach was utilized for the more hydrophobic 2T and 3T CPT azide tails. Interestingly, γ-cyclodextrin effectively solubilized both multivalent constructs without any precipitation. MALDI-TOF MS analysis of the crude reaction mixtures revealed single peaks at 69,167 Da and 69,971 Da, which correspond to HSA–MI–OEG–COT–2T CPT and HSA–MI– OEG–COT–3T CPT, respectively **(Figure 1c-d)**. These results indicate that SAPLabTech successfully addresses the solubility challenges posed by highly hydrophobic multivalent drug payloads, enabling quantitative SPAAC conjugation under mild aqueous conditions. Detailed experimental methods are included in the Methods section.

### Purification and Characterization of Protein-Drug Conjugates

The purification of the HSA–MI–OEG–COT–CPT conjugates was carried out in two stages. Initially, ion-exchange chromatography (IEC) was used to remove excess unreacted CPT azide and γ-cyclodextrin that did not bind to the column under the chosen conditions **(Figure 2a–c)**. The fractions of conjugated HSA were then combined and desalted by chromatography to remove residual salts from the IEC process **(Figure 2d–f)**. Comprehensive procedures are outlined in the Methods section. The desalted conjugate fractions were collected in Milli-Q water and lyophilized to yield purified HSA–MI–OEG–COT–CPT conjugates in solid form, which were stored at -20°C for subsequent physicochemical and biological evaluations. The lyophilized conjugates were dissolved in sodium phosphate buffer (pH 7.4) and examined using MALDI-TOF MS as outlined in the Methods section. The isolated HSA–MI–OEG–COT–1T CPT, HSA– MI–OEG–COT–2T CPT, and HSA–MI–OEG–COT–3T CPT conjugates displayed distinct, single peaks at 68,142 Da, 69,084 Da, and 69,428 Da, respectively **(Figure 1a-c)**. The conjugates’ structure is illustrated in **Figure 4**. Refer to **Table 1** for the summarized MALDI-TOF-MS results.

**Table 1:** Summary of MW obtained for protein conjugate in MALDI-TOF MS analysis.

| Sr.No. | Molecules | Expected MW<br>for protein<br>conjugate (Da) | MW obtained<br>for RM (Da) | MW obtained for<br>purified protein<br>conjugate |
| --- | --- | --- | --- | --- |
| 1. | HSA-MI-COT-1T CPT | 68454 | 68337 | 68142 |
| 2. | HSA-MI-COT-2T CPT | 69216 | 69167 | 69084 |
| 3. | HSA-MI-COT-3T CPT | 69978 | 69971 | 69428 |

**Figure 2:**
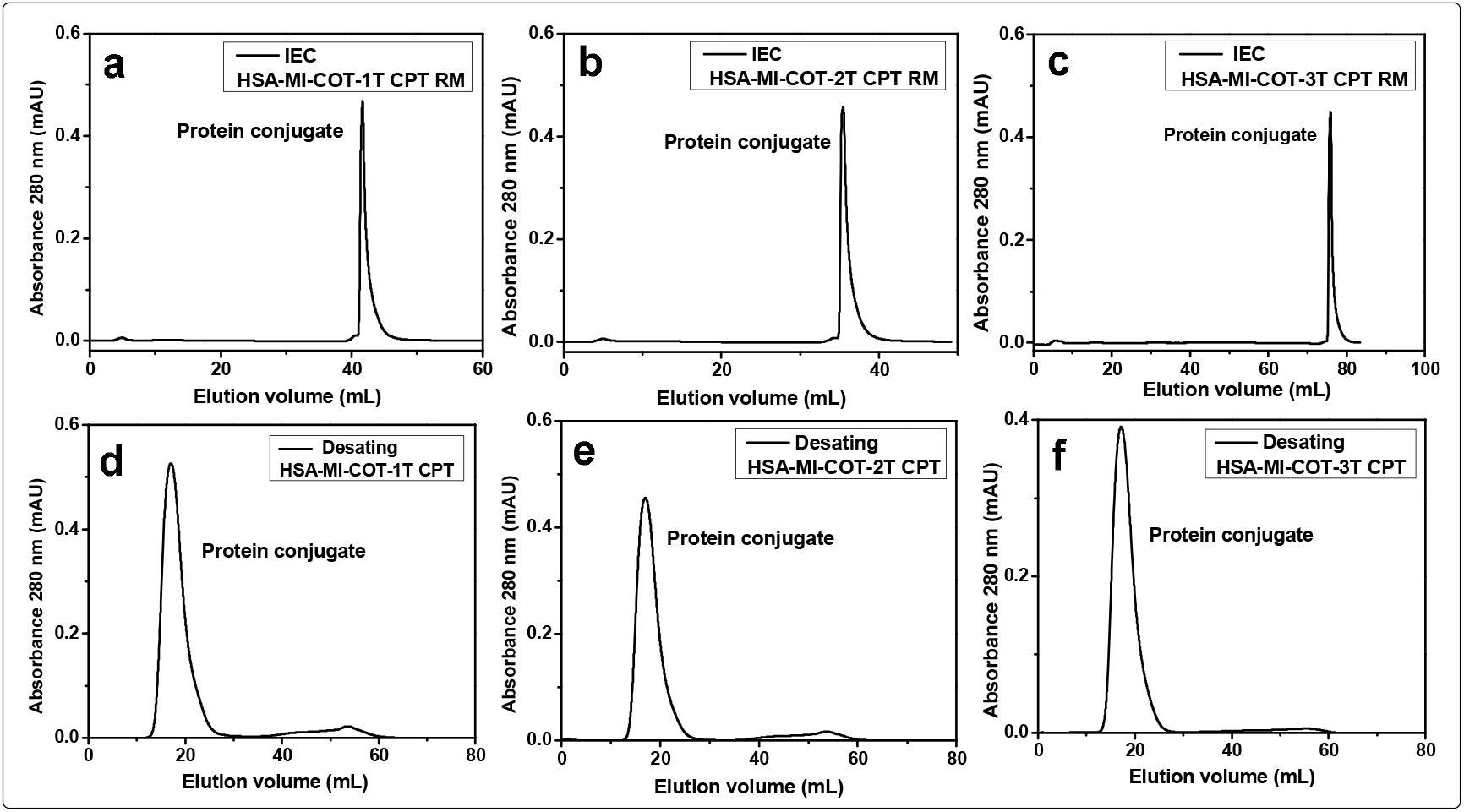
The purification process consists of ion exchange chromatography (IEC) followed by desalting. The ion exchange chromatograms illustrate the separation of the protein conjugate from the reaction mixture, which includes Gymma CD and unreacted probes (labelled a, b, c). This is followed by a desalting chromatogram that demonstrates the removal of salts from the purified conjugates (labelled d, e, f).

**Figure 3:**
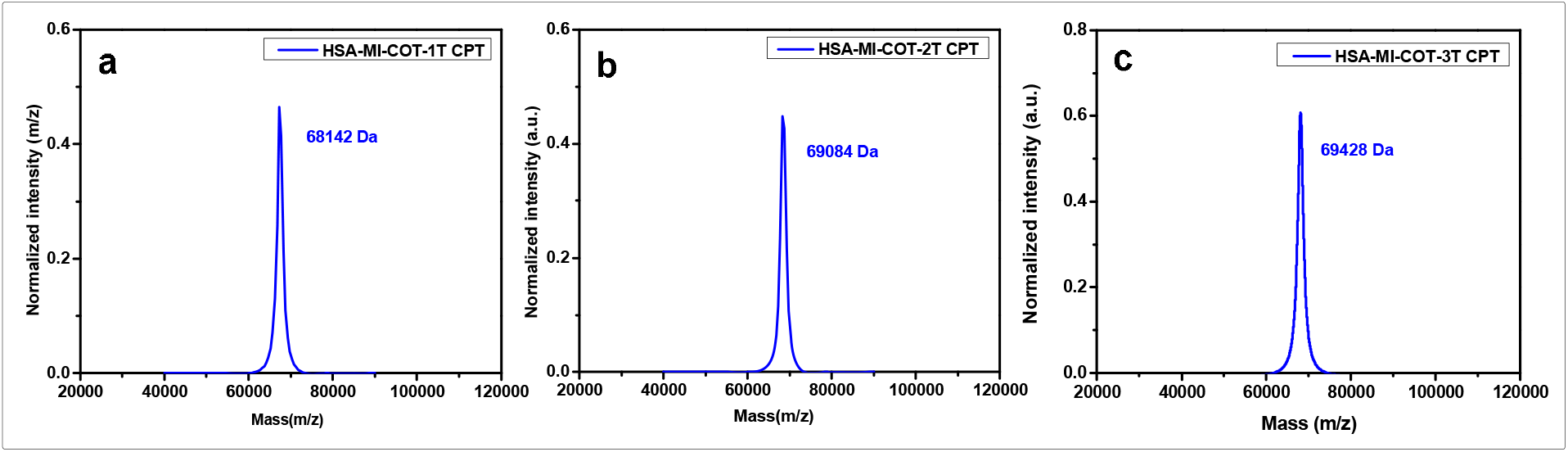
The MALDI-TOF MS analysis of the purified protein conjugate indicates high purity.

**Figure 4:**
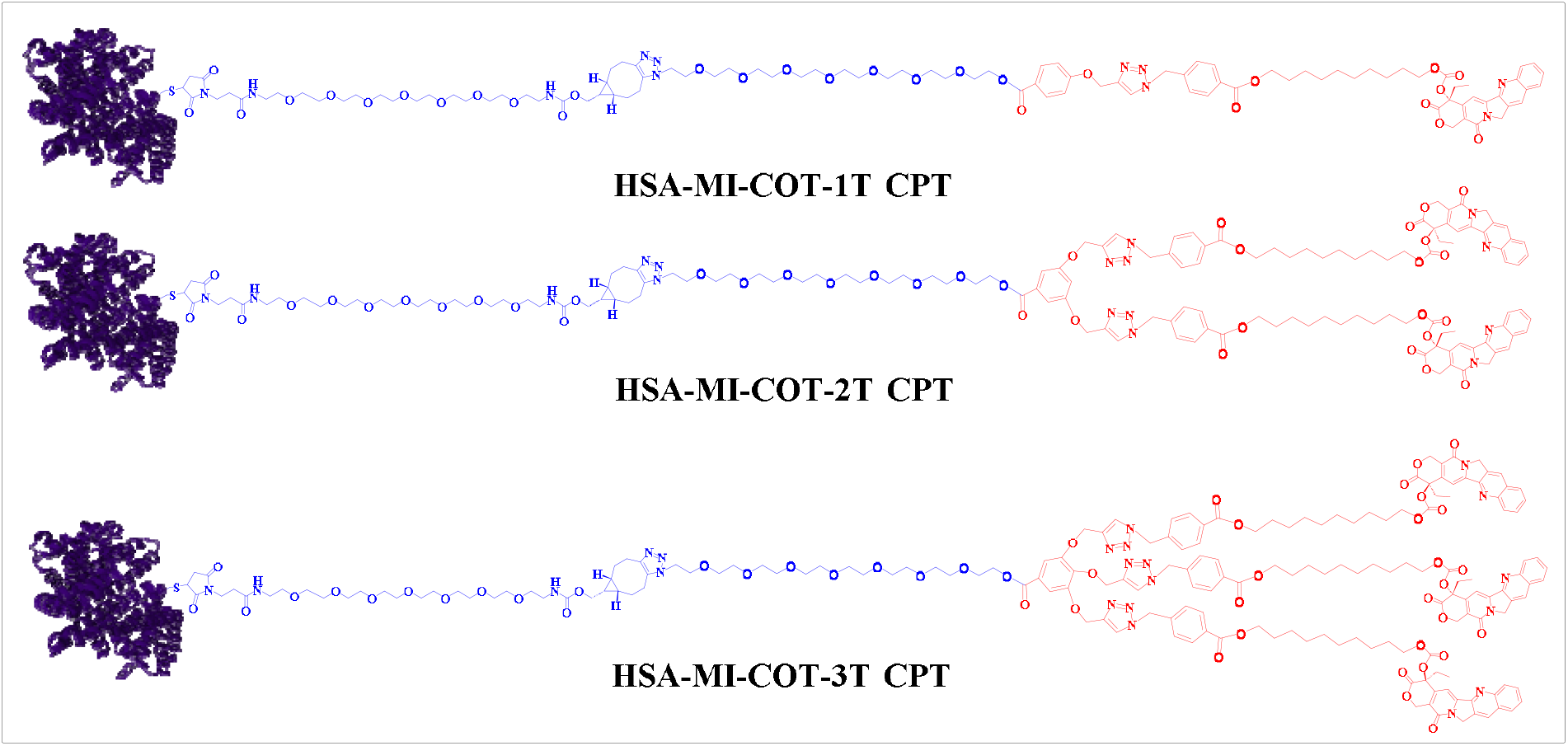
The composition of the protein conjugate suggested a varying number of CPT molecules associated with each protein.

### Self-assembly Studies of SPDCs

Dynamic light scattering (DLS) was used to assess the hydrodynamic size and self-assembly behavior of purified protein-drug conjugates in aqueous solution. The DLS profiles indicated the formation of nanoscale assemblies for all three conjugates, demonstrating that the amphiphilic nature of the constructs drove nanoparticle formation. The average hydrodynamic diameters (Z-average) of the nanoparticles formed by three bioconjugates were measured to be around 11 nm **(Table 2)**. The narrow size distribution observed for all conjugates suggests the formation of relatively uniform nanoassemblies in solution. Zeta potential measurements of the HSA–CPT conjugates in Milli-Q water showed moderately negative surface charges, with values ranging from −12 mV to −14 mV. **(Table 2)**.

**Table 2.** Summary of self-assembly characteristics of HSA–CPT conjugates, including hydrodynamic size (DLS, Z-average diameter) and surface charge (zeta potential). The data confirm the formation of nanoscale particles with moderately negative surface charge, indicative of stable nanoparticle assemblies in aqueous media.

| Sr. No. | Proteins | DLS<br>Size (d / nm) | PDI | Zeta potential<br>(mv) |
| --- | --- | --- | --- | --- |
| 1. | HSA | 7 | 0.102 | -4 |
| 2. | HSA-MI-COT-1T CPT | 11 | 0.476 | -12 |
| 3. | HSA-MI-COT-2T CPT | 12 | 0.613 | -14 |
| 4. | HSA-MI-COT-3T CPT | 11 | 0.458 | -12 |

**Table 3:**
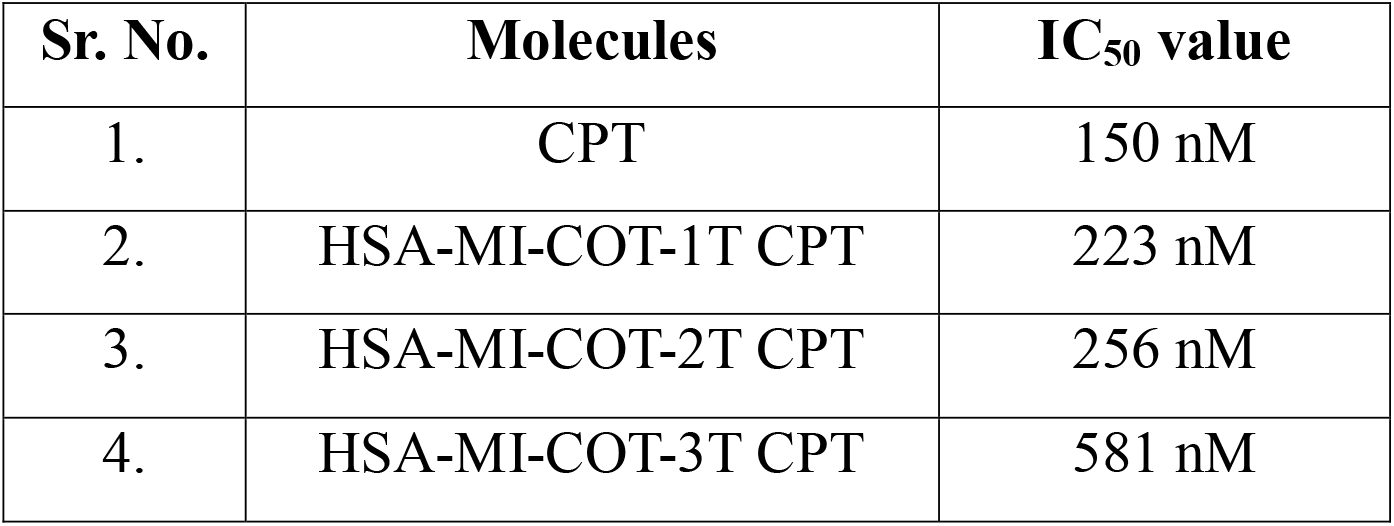
Summary of IC_50_ values obtained during MTT assay in MCF-7 cell line.

### In Vitro Cytotoxic Studies of SPDCs

The cytotoxic effects of the conjugates were assessed in MCF-7 cells, a breast cancer cell line. Dose-response experiments were conducted for each conjugate as well as for free camptothecin (CPT) after a 72-hour incubation at 37 °C. The free CPT showed an IC_50_ of 150 nM **(Figure 5a)**, consistent with its well-documented potent cytotoxicity. Encouragingly, all HSA conjugates demonstrated noticeable, concentration-dependent dose-response behavior. The IC50 values for bioconjugates HSA–MI–OEG–COT–1T/2T/3T CPTs were found to be 223 nM, 256 nM, and 581 nM, respectively **(Figure 5b-c)**. Despite conjugates exhibiting somewhat lower potency than free CPT, they still showed significant cytotoxicity. This finding suggests effective cellular uptake and release of the active drug, thereby confirming the functional integrity of the acid-sensitive linker system.

**Figure 5:**
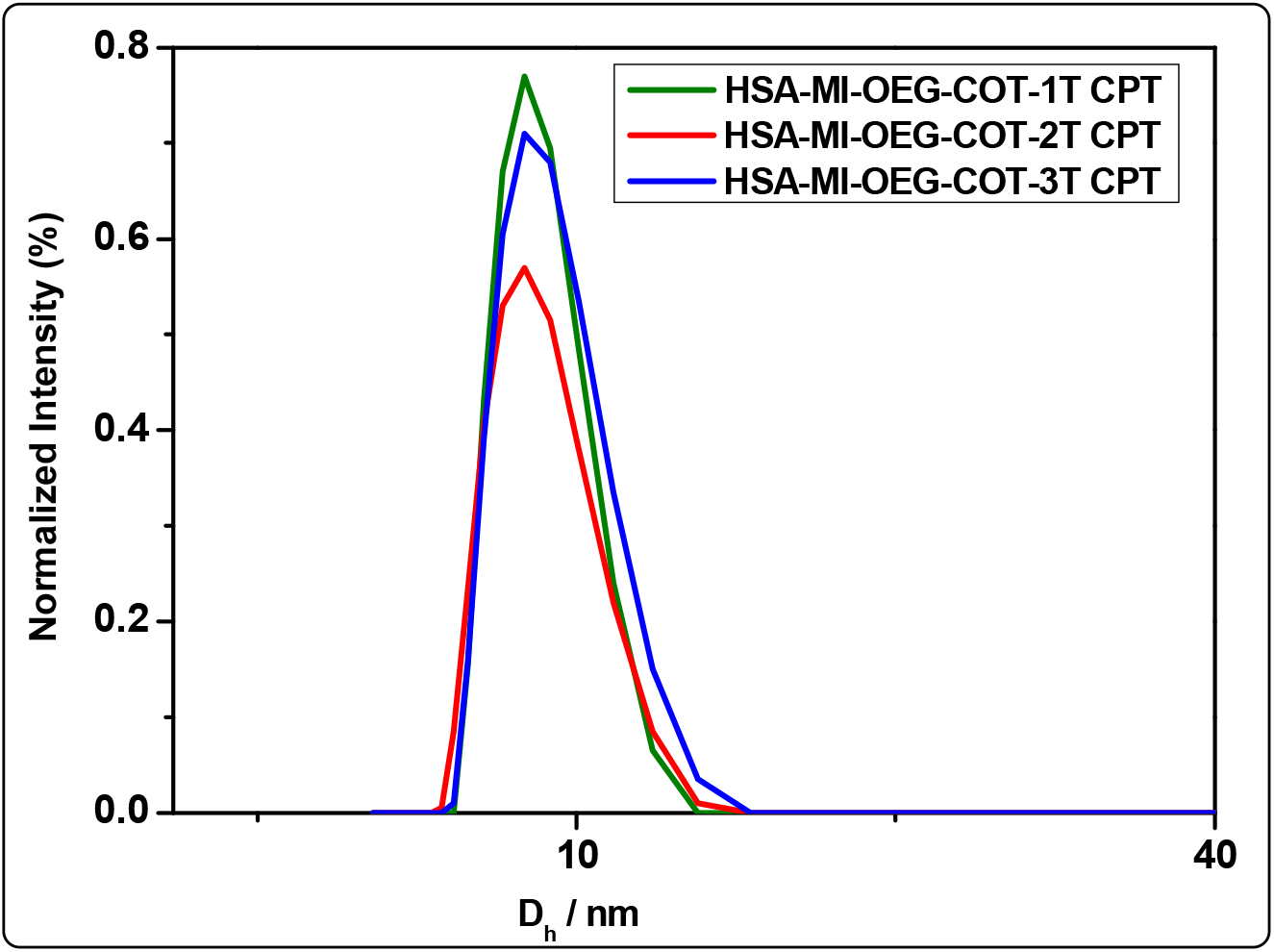
Dynamic light scattering (DLS) intensity-based size distribution plots of HSA–MI– OEG–COT–1T CPT, HSA–MI–OEG–COT–2T CPT, and HSA–MI–OEG–COT–3T CPT measured at 25 °C in aqueous solution. All conjugates exhibit a single dominant nanoscale population with Z-average hydrodynamic diameters of 11,12, and 11 nm, respectively.

**Figure 6:**
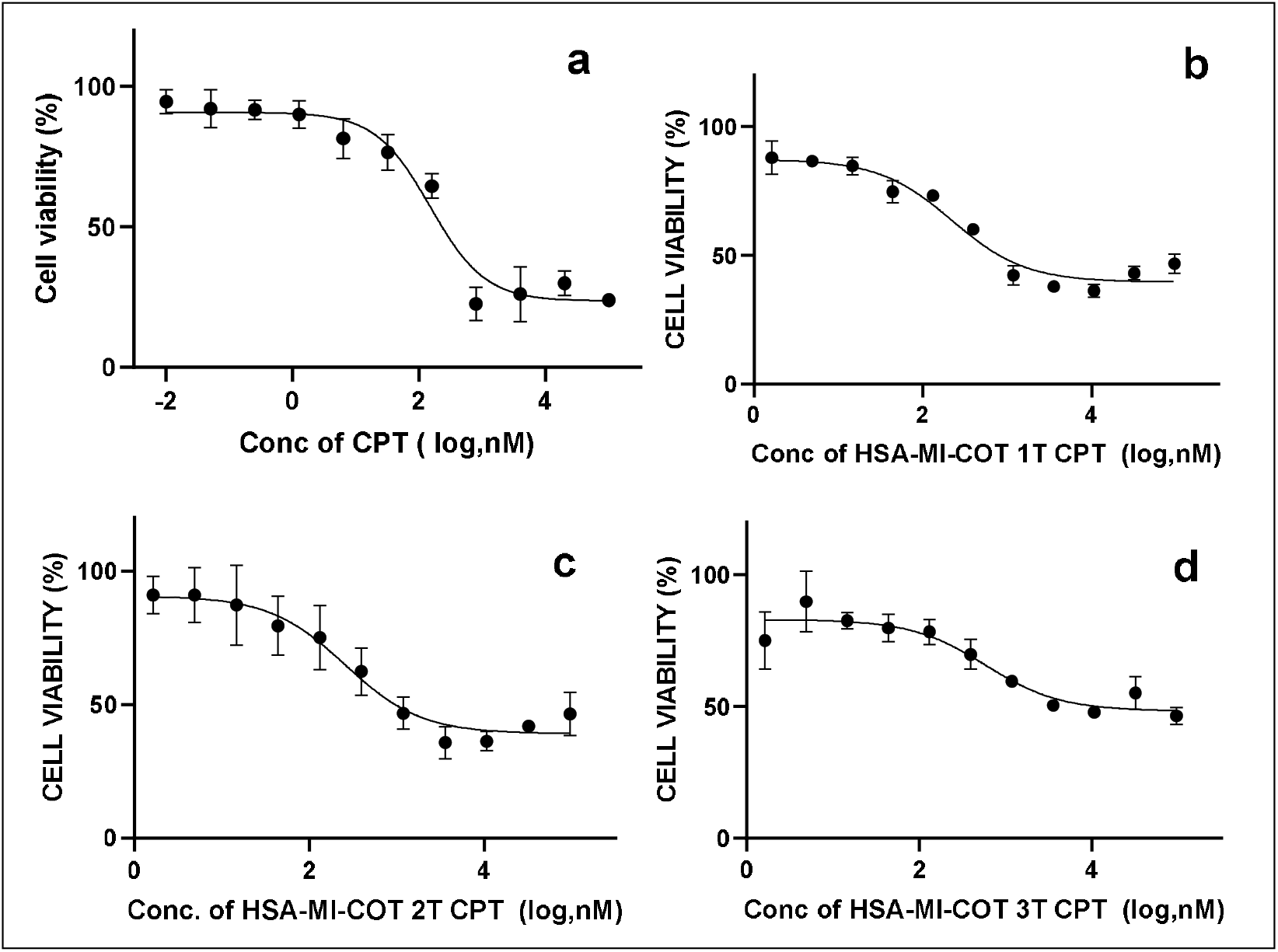
Cytotoxicity assessments of HSA-MI-COT 1T/2T/3T CPT in the MCF7 cell line were carried out after a 72-hour incubation at 37°C. The free CPT, used as a reference, along with HSA, served as control molecules during the experiments.

## Discussion

The hydrophobicity of cytotoxic payloads remains one of the most significant challenges in ADC development, and numerous strategies have been explored to address this limitation.^18–20,22,28^ the most commonly adopted approach—introducing solubility-enhancing tags within the linker region—has partially alleviated some issues associated with payload hydrophobicity but has not resolved them entirely.^13,21,24–26^ Furthermore, appending additional solubility tags increases the synthetic complexity of ADC production. PEG-based solubility tags, in particular, are known to elicit anti-PEG immune responses in humans.^62,63^ Critically, none of these approaches shields the cytotoxic drug from the biological milieu, leaving the payload vulnerable to premature release or non-specific interactions with off-target biological components.^9,11,17^

In the present study, we asked whether the hydrophobic character of cytotoxic drugs could be redirected as a driving force for the design of next-generation XDCs, rather than treated solely as a liability. To this end, we deliberately increased the hydrophobicity of CT by appending a decyl alkyl chain, and designed self-assembling facially amphiphilic protein–drug conjugates inspired by our earlier work on self-assembling FAPs.^29–36^ A central challenge in this work was the site-specific bioconjugation of hydrophobic drugs to HSA. This was accomplished using our in-house developed supramolecule-assisted protein labeling technology (SAPLabTech).

As anticipated, all newly synthesized bioconjugates exhibited excellent aqueous solubility, despite bearing highly lipophilic drug molecules. This enhanced solubility is attributed to the combined hydrophilic contributions of the HSA scaffold and the OEG linker. Bioconjugates 1–3 self-assembled into ultra-small protein nanoparticles with hydrodynamic diameters in the range of 11 nm, in which the drug molecules are localized within the hydrophobic core, and HSA are displayed on the outer surface. DLS and neutron scattering analyses confirmed the spherical geometry of these assemblies. The oligomeric state and molecular weight of the protein nanoparticles increased systematically with increasing drug-to-protein ratio (DPR), consistent with a hydrophobically driven self-assembly mechanism. Dose–response studies conducted in MCF-7 breast cancer cells demonstrated that native CPT and all three bioconjugates retained cytotoxic activity. However, modest reductions in potency were observed for bioconjugates 1–3 relative to free CT.

## Conclusion

In summary, we have demonstrated that the hydrophobicity of cytotoxic payloads can be strategically exploited to design next-generation protein–drug conjugates. The resulting facially amphiphilic protein–drug conjugates self-assemble into molecularly defined, ultra-small protein nanoparticles. In vitro studies confirm that these nanoparticles retain potent activity against breast cancer cells. Nevertheless, several important questions remain unresolved: What cellular uptake mechanism(s) do these nanoparticles employ? At what intracellular site and under what conditions is the drug released from the protein conjugate? What accounts for the modest decrease in IC_50_values observed at higher drug loadings? These questions are currently under active investigation in our laboratory, and the findings will be reported in due course.

## Experimental Methods

### Materials

All reagents and chemicals were sourced from reliable commercial vendors and used without further purification.

### Nuclear Magnetic Resonance (NMR) Spectroscopy

The ^1^H and ^13^C NMR spectra were recorded using a Bruker 400 MHz spectrometer. The proton chemical shifts (δ) are reported in parts per million (ppm) relative to the residual solvent peaks in CDCl_3_ (δ 7.26), while the carbon chemical shifts are referenced to CDCl_3_ (δ 77.16). High-purity CDCl_3_ solvent was purchased from Sigma-Aldrich.

### Protein

The human serum albumin (HSA, 66.6 kDa) was chosen as a representative cysteine-containing protein. These were obtained from Sigma-Aldrich, and their molecular weights were confirmed before modification.

### Bioconjugation scheme for the synthesis of HSA-MI-OEG-COT

The 1.2 eq MI-OEG-COT probe was dissolved in 500 μL of sodium phosphate buffer at pH 7.2, containing 0.1% DMSO. The solution was vortexed for 15 minutes until it became clear. Following this, 500 μL of a protein solution, also prepared in sodium phosphate buffer at pH 7.2, was added. The final protein concentration was kept at 100 μM. The reaction mixture was rotated for 16 hours at 20 rps on a rotaspin at room temperature. After the procedure was completed, the mixture was analyzed using MALDI-TOF mass spectrometry. To evaluate the extent of protein modification, samples were directly extracted from the reaction mixture with a pipette and analyzed.

### SPAAC Conjugation scheme via Supramolecular Assisted Protein Labeling Technology (SAPLabTech)

A total of 10 µL of CPT-azide tail (at a concentration of 50 mM in DMSO) was mixed with 190 µL of gamma cyclodextrin (200 mM in PBS at pH 7.2) at a molar ratio of 1:76 for 1T /2T CPT azide tail (for 3T CPT azide tail, the molar ratio was 1:160). The resulting mixture was sonicated for 10 minutes and subsequently diluted with PBS containing 0.02% Triton X-100 (0.2 µL Triton X-100 per mL) to a final volume of 500 µL, allowing the solution to equilibrate at room temperature for 30 minutes on a rotaspin set to 20 RPM. After this step, 500 µL of HSA-functionalized maleimide COT RM (HSA-MI-OEG-COT RM) was added to the mixture and incubated for 24 hours at room temperature on a rotaspin at 20 RPM. The resulting protein concentration was 50 µM. Upon completion of the reaction, the RM was characterized using MALDI-TOF-MS.

### Matrix Preparation and Molecular Weight Determination

The molecular weights of both native and modified proteins were accurately measured using MALDI-TOF MS (Shimadzu MALDI-8020). A solution was prepared by dissolving 15 mg of sinapinic acid in 1 mL of a 70:30 mixture of water and acetonitrile containing 0.1% TFA. Protein samples were combined with the matrix in a 1:5 ratio, placed onto MALDI plates (2 µL), and allowed to air-dry for 30 minutes. Mass scanning was conducted over a range of 40,000–80,000 Da, with a protein concentration set at 50 µM.

### Sample Preparation for MALDI-TOF MS

3 µL of the reaction mixture was combined with 15 µL of the matrix, vortexed, and then placed onto the MALDI plate. The MALDI-TOF MS assessment enabled precise tracking of protein modifications and verification of conjugate formation.

### Purification of modified HSA -CPT conjugates

Purification of modified proteins was performed using an AKTA Pure chromatography system (Cytiva, USA). The purification procedure included ion-exchange chromatography (IEC) followed by desalting or buffer exchange. The first step in the purification procedure involves ion-exchange chromatography (IEC), during which excess unreacted probes and gymma CD are removed, as they do not bind to the column. The IEC process was carried out using a HiScreen Q FF anion-exchange column at pH 10. During this stage, the column was pre-equilibrated with a buffer composed of 50 mM Tris base at pH 10 before the sample was added. After injecting the sample, the column was operated for at least 5 column volumes (CVs) without utilizing any elution buffer, ensuring that all unreacted probe and gymma CD were eliminated. The protein conjugate was then eluted with a buffer containing 50 mM Tris base and 1 M NaCl at pH 10. After ion-exchange chromatography, the salt was removed using a HiPrep 26/10 Desalting column. Milli-Q water was used as both the elution and equilibration buffer. Once the sample was injected, the protein conjugate fraction was eluted first, followed by the salt. This desalted fraction was then lyophilized to yield a pure protein conjugate powder.

### MALDI-TOF MS for purified protein conjugate

The lyophilized protein conjugate was examined using MALDI-TOF mass spectrometry.

Protein solution A - A lyophilized protein conjugate weighing 1 mg was reconstituted in sodium phosphate buffer adjusted to a pH of 7.4.

Triton X-100 solution - 2 µL of Triton X-100 was added to 1 mL of sodium phosphate buffer at pH 7.4.

Sample for MALDI-TOF MS - 4 µL of the Triton X-100 solution was combined with 96 µL of protein solution A. The analysis was conducted using the same matrix as previously mentioned.

### Dynamic Light Scattering (DLS) studies

The hydrodynamic diameter of the protein complexes was measured using a Zetasizer Nano 2590 device (Malvern Instruments, UK). Protein samples were prepared at 200 µM in 50 mM sodium phosphate buffer, pH 7.4. A 1 mL filtered sample was placed in disposable polystyrene cuvettes, and the average particle size was measured at a scattering angle of 90° under standard conditions.

### Zeta potential measurements

The measurements were performed using a Zetasizer Nano ZS (Malvern Instruments Ltd., UK). Protein samples were prepared at a concentration of 5 mg/mL in Milli-Q water.

### In vitro cytotoxic studies

The comprehensive guidelines for culturing the MCF-7 cell line in mammalian cells, including cell seeding and splitting procedures, are provided in the supplementary information.

#### Control experiment

The purpose of the experiment was to measure the IC_50_ values of the unbound drug CPT as a reference. The drug stock solution was prepared at 20 mM (100 µM). Dilutions were prepared at a 1:5 ratio, with two control lanes containing only media and 0.5% DMSO. Each concentration was evaluated in triplicate, and the 96-well plate was incubated at 37 °C for 72 hours. The MTT assay was utilized to determine cell viability.

#### Sample studies

For HSA-MI-COT-1T CPT, HSA-MI-COT-2T CPT, and HSA-MI-COT-3T CPT, the stock solution was prepared in 1X PBS, pH 7.4, at a stock concentration of 250 µM (96 µM). Further dilutions were prepared at a 1:3 ratio with media, while control lanes contained a mixture of media and PBS. Each concentration was tested in triplicate, and the 96-well plate was maintained at 37 °C for 72 hours. The MTT assay was employed to assess cell viability.

### MTT Assay

After 72 hours of incubation, the drug solution was removed from the wells, and 100 µL of MTT solution (0.5 mg/ml) was added. The plate was incubated for 4 hours at 37 °C. Once the incubation was complete, the MTT solution was removed, and the Formazan crystals were dissolved in 100 µL of DMSO. The absorbance of each well was measured at 570 nm, and the average absorbance for each concentration was calculated. Cell viability was determined by dividing the absorbance at a specific concentration by the average absorbance of the DMSO control and multiplying by 100 to express cell viability as a percentage. The IC_50_ curve was generated from the data to derive the IC_50_ values for the drugs and probes.

## ASSOCIATED CONTENT

### Supplementary Information

#### Author Information

**Notes**: We have filed Indian and US patents for the technology disclosed in this paper.

## ACKNOWLEDGEMENTS

We want to thank IISER-Pune and Molbio Diagnostic Pvt. Ltd. for their funding.

**Scheme 1:**
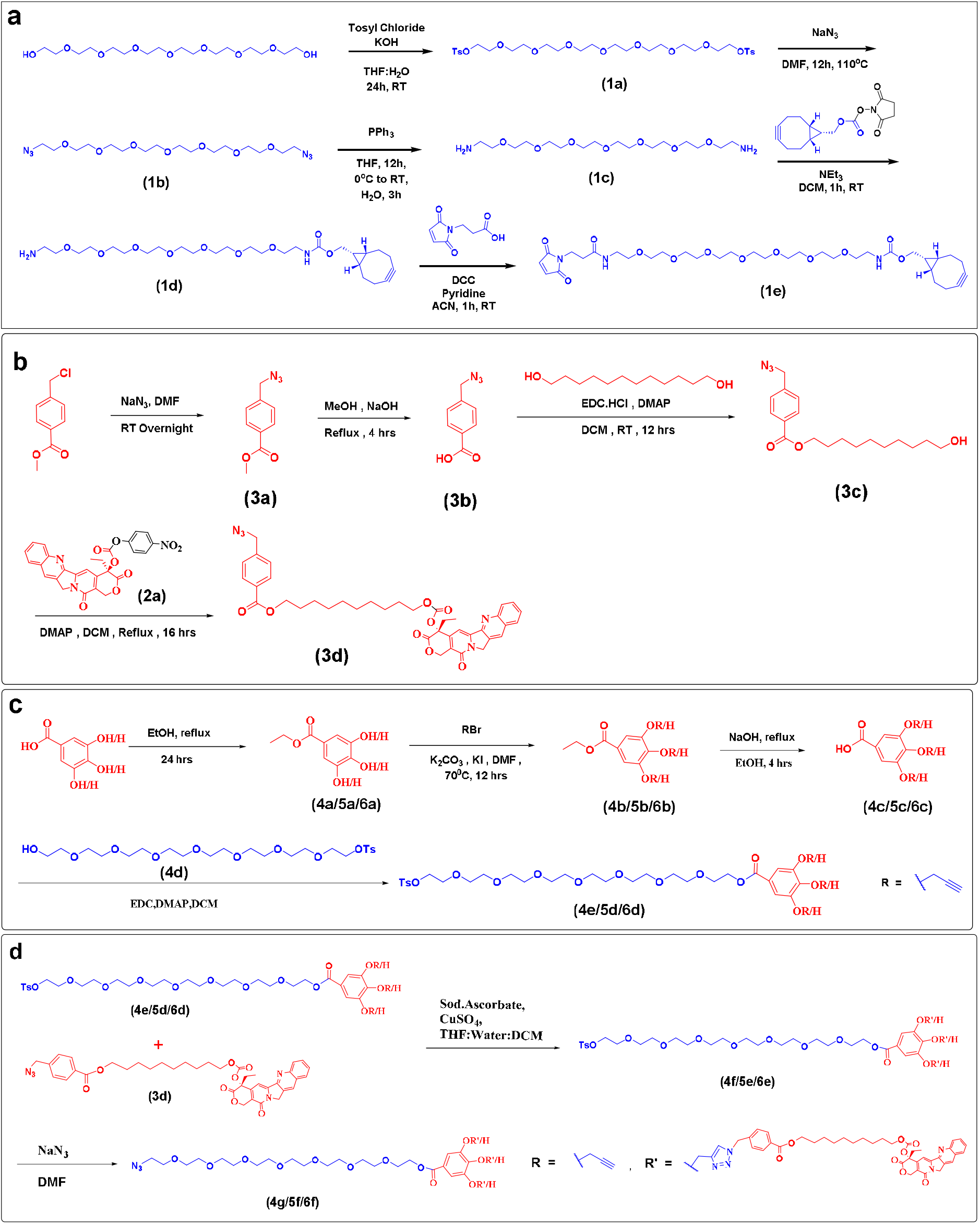
Synthetic scheme for CPT-functionalized probes and modular tail constructs. **a**. Bifunctional probe bearing a maleimide for cysteine conjugation and a cyclooctyne for SPAAC reactions. **b**. Synthesis of CPT–intermediate, in which CPT is linked to a C10 hydrophobic moiety via a pH-sensitive carbonate linker. **c**. Preparation of octaethylene glycol– based hydrophilic spacers functionalized with 1T/2T/3T alkyne groups. **d**. Synthesis of 1T/2T/3T CPT-azide tails via azide–alkyne click chemistry.

**Scheme 2:**
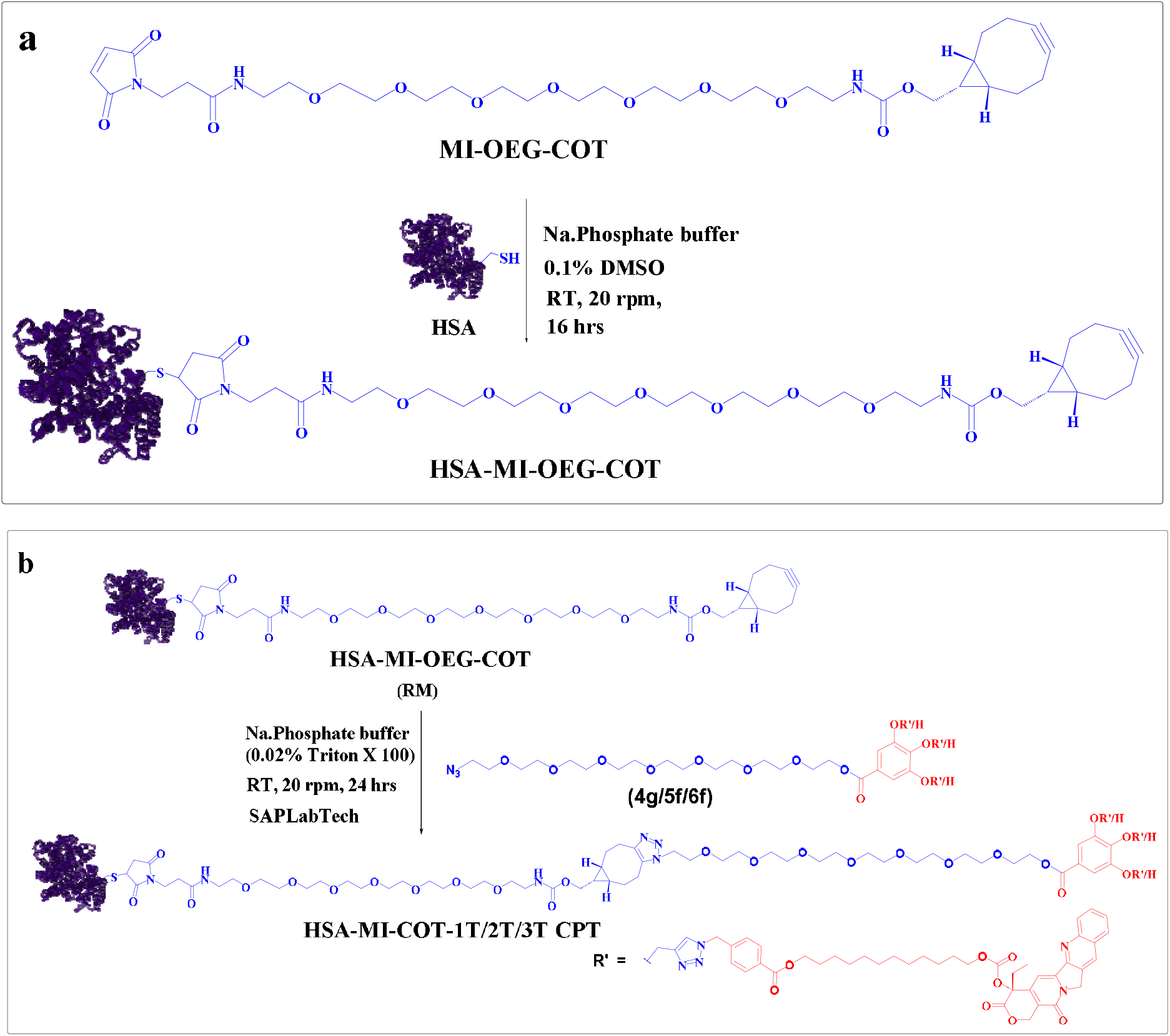
Scheme for the bioconjugation and post-click functionalization of HSA. **a**. Bioconjugation of human serum albumin (HSA) with MI–OEG–COT via cysteine-selective maleimide chemistry under mild conditions. **b**. post-click conjugation of 1T/2T/3T CPT–azide tails to HSA–MI–OEG–COT following γ-cyclodextrin inclusion complex formation, yielding CPT-functionalized HSA conjugates.

## Notes

### Competing Interest Statement

We have filed Patents for the technology disclosed in this paper

