## Supplementary material for "Extreme Hydrophobicity of Cytotoxic Drugs Enables Design of Next Generation Antibody-Drug Conjugates Nanotherapeutics": Supp Info

**Table of Contents:**

1. **Synthesis of chemical probes and their intermediates**
   1.1. Synthetic scheme for the synthesis of MI-OEG-COT Spacer
   1.2. Synthetic scheme for activation of camptothecin
   1.3. Synthetic Scheme for the synthesis of the CPT-azide
   1.4. The general procedure for the synthesis of the hydrophilic spacer and its intermediates
   1.5. The general procedure for the synthesis of the CPT-azide tail and its intermediates
2. **Cellular studies**
   2.1. Protocol for Mammalian Cell Culture: MCF7 cells
   2.2. Procedure for experiment set-up for cell viability studies.
3. **NMR data**

**Experimental results**

1. **Synthesis of chemical probes and their intermediates**

**1.1. Synthetic scheme for the synthesis of MI-OEG-COT spacer**

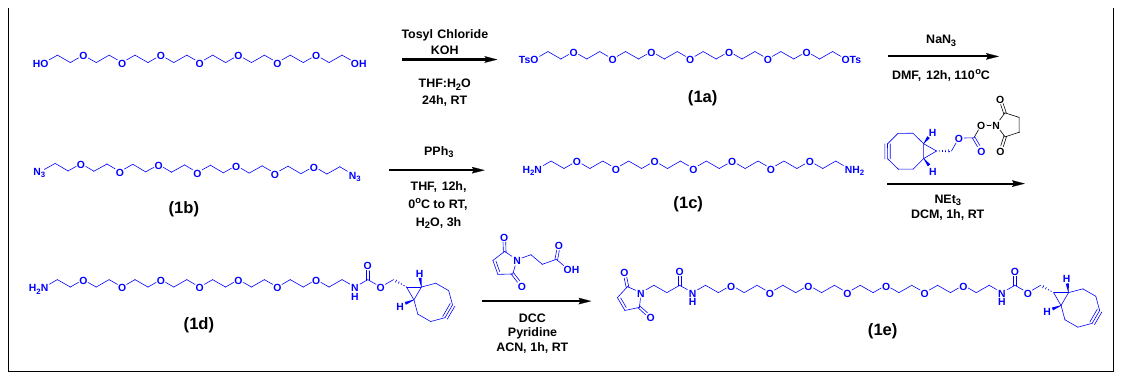

**Scheme 1:** The synthetic pathway for synthesizing a chemical probe features a maleimide as the reactive head group, which targets cysteine. Simultaneously, it includes a cyclooctyne group on the other side for click reactions.

In a round-bottom flask (RBF) that has been dried in an oven, octaethylene glycol (1.0 eq.) and tosyl chloride (TsCl) (3 eq.) were mixed and dissolved in THF while at 0 °C. After one hour, a KOH (7 eq.) solution, prepared from a 1:1 blend of THF and water, was gradually added to the reaction mixture, which was stirred at room temperature for 24 hours. Once the reaction was finished, it was quenched using an aqueous sodium bicarbonate solution and extracted three times with ethyl acetate. The organic phase was dried with sodium sulphate (Na_2_SO_4_) and concentrated to yield the crude product. This material was then purified through normal-phase chromatography (NPC) using methanol (MeOH) and dichloromethane (DCM) as the eluting solvents.

The ditosylate (1a) (1 eq.) was then dissolved in dimethylformamide (DMF) and heated to 110 °C. At this point, sodium azide (NaN_3_) (7 eq.) was added to the reaction mixture, which was allowed to react for 12 hours. Following this, the reaction mixture was concentrated and purified by NPC, without any prior workup, utilizing MeOH and DCM as the eluent to isolate product (1b).

The azide obtained (1 eq.) was dissolved in THF, and a solution of PPh_3_ (3 eq.) in THF was gradually added while stirring at 0 °C. The reaction mixture was kept at room temperature for 12 hours. Once the reaction was complete, water was added to the mixture and stirred for 3 hours. This was followed by extraction using DCM, and the aqueous layer was evaporated under vacuum to obtain the crude product, which was then purified through normal-phase chromatography with MeOH/DCM as the eluent, yielding compound (1c).

In an oven-dried round-bottom flask (RBF), the previously synthesized amine (2 eq.) and BCN-N-hydroxysuccinimidyl carbonate (1 eq.) were dissolved in dichloromethane (DCM) while stirring. Then, triethylamine (Et_3_N) (1.1 eq.) was added slowly at room temperature and allowed to react for 20 minutes. After the completion of the reaction, excess water was added, and the mixture was extracted with DCM. The organic layer was then concentrated and purified via NPC with MeOH/DCM as the eluent to obtain compound (1d).

Next, maleimide propionic acid (1.2 eq.) and compound (1d) (1 eq.) were dissolved in acetonitrile (ACN). Pyridine (0.2 eq.) and dicyclohexylcarbodiimide (DCC) (1.2 eq.) were added to the mixture. The reaction mixture was stirred for 1 hour at room temperature. Afterwards, the reaction mixture was quenched with saturated sodium bicarbonate (NaHCO_3_) and extracted with DCM. The combined organic layers were dried over Na_2_SO_4_, concentrated, and subsequently purified using normal phase chromatography (NPC) with MeOH/DCM, resulting in the final compound (1e).

1.1.1. **Synthesis of MI-OEG-COT spacer and its intermediates**
       1.1.1.1. **Compound (1a)**

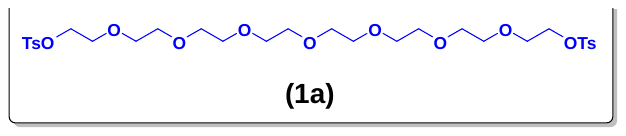

Mol. formula: C_30_H_46_O_13_S_2_

Mol. Weight: 678.80

Physical appearance: Yellowish liquid

Yield: 88.01%

The **compound (1a)** is synthesized following **synthetic scheme 1.1**, beginning with octaethylene glycol (1.00 g, 2.67 mmol), tosyl chloride (1.53 g, 8.01 mmol), and a KOH solution (1.05 g, 18.7 mmol) prepared in 4 mL of a THF: water mixture (1:1). The resulting crude product was purified by normal-phase chromatography using methanol/dichloromethane (MeOH/DCM) as the eluent, yielding a pure product (1.6 g, 2.35 mmol) with an R_f_ value of 0.4 in 5% MeOH/DCM.

**^1^H NMR (400 MHz, CDCl_3_) δ:** 7.86 – 7.74 (m, 4H), 7.34 (d, *J* = 8.1 Hz, 4H), 4.22 – 4.08 (m, 5H), 3.70 – 3.66 (m, 4H), 3.65 – 3.60 (m, 16H), 3.58 (s, 8H), 2.44 (s, 6H).

**^13^C NMR (100 MHz, CDCl_3_) δ:** 144.93, 133.09, 129.95, 128.09, 70.82, 70.68, 70.63, 70.59, 69.37, 68.78, 21.75.

**MALDI-TOF-MS (M+Na):** 701.32

1.1.1.2. **Compound (1b)**

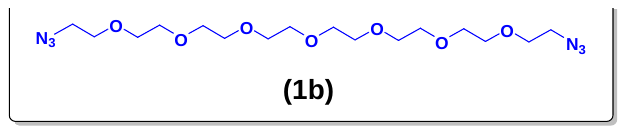

Mol. formula: C_16_H_32_N_6_O_7_

Mol. Weight: 420.46

Physical appearance: Pale yellow

Yield: 77%

The **compound (1b)** is synthesized using **synthetic scheme 1.1**, starting with compound (1a) (1.0 g, 2.78 mmol), NaN_3_ (1.27 g, 19.59 mmol), and DMF (30 mL). The obtained crude product was purified by normal-phase chromatography (NPC) using a methanol/dichloromethane (MeOH/DCM) mixture as the eluent, resulting in a pure product (0.900 g, 2.14 mmol) with an R_f_ value of 0.4 in 5% MeOH/DCM.

**^1^H NMR (400 MHz, CDCl_3_) δ:** 3.72 – 3.60 (m, 29H), 3.38 (t, *J* = 5.1 Hz, 4H).

**^13^C NMR (100 MHz, CDCl_3_) δ:** 70.76, 70.73, 70.70, 70.64, 70.10, 50.76.

**MALDI-TOF-MS (M+Na):** 459.23

1.1.1.3. **Compound (1c)**

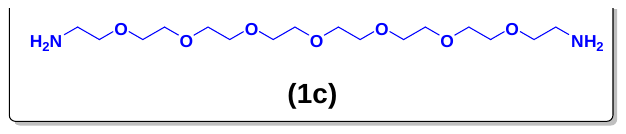

Mol. formula: C_16_H_36_N_2_O_7_

Mol. Weight: 368.47

Physical appearance: Pale yellow oil

Yield: 45.2%

The **compound (1c)** is synthesized using **synthetic scheme 1.1**, starting with compound (1b) (1.06 g, 2.52 mmol), PPh_3_ (1.98 g, 7.56 mmol), THF (50 mL), and water (50 mL). The crude product obtained was purified by normal-phase chromatography (NPC) using a methanol/dichloromethane (MeOH/DCM) mixture as the eluent, resulting in the isolation of the pure product (0.420 g, 1.14 mmol) with an R_f_ value of 0.1 in 5% MeOH/DCM.

**^1^H NMR (400 MHz, CDCl_3_) δ:** 3.67 – 3.59 (m, 25H), 3.49 (t, *J* = 5.2 Hz, 4H), 2.85 (t, *J* = 5.2 Hz, 4H).

**^13^C NMR (100 MHz, CDCl_3_) δ:** 73.34, 70.61, 70.59, 70.30, 41.74.

**MALDI-TOF-MS (M+Na):** 391.30

1.1.1.4. **Compound (1d)**

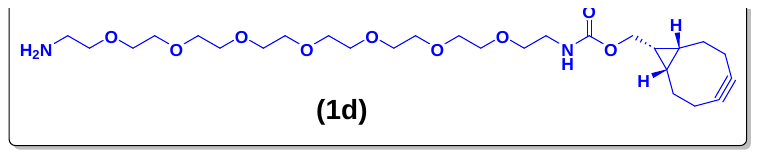

Mol. formula: C_27_H_48_N_2_O_9_

Mol. Weight: 544.68

Physical appearance: Yellowish liquid

Yield: 68.5%

The **compound (1d)** is synthesized according to **synthetic scheme 1.1**, starting with compound (1c) (0.252 g, 0.686 mmol), BCN-N-hydroxysuccinimidyl carbonate (0.100 g, 0.343 mmol), NEt_3_ (0.038 g, 0.378 mmol), and DCM (2 mL). The obtained crude product was purified by normal phase chromatography (NPC) using a mixture of MeOH and DCM as the eluent. This resulted in a pure product (0.128 g, 0.235 mmol) with an R_f_ value of 0.3 in 5% MeOH/DCM.

**^1^H NMR (400 MHz, CDCl_3_) δ:** 7.95 (s, 2H), 4.12 (d, *J* = 8.2 Hz, 2H), 3.94 – 3.84 (m, 2H), 3.76 – 3.60 (m, 23H), 3.55 (t, *J* = 5.2 Hz, 2H), 3.35 (q, *J* = 5.4 Hz, 2H), 3.18 (q, *J* = 5.1 Hz, 2H), 2.34 – 2.14 (m, 6H), 1.65 – 1.51 (m, 2H), 1.35 (t, *J* = 14.7 Hz, 4H).

**^13^C NMR (100 MHz, CDCl_3_) δ:** 157.06, 139.38, 114.16, 98.94, 70.56, 70.40, 70.37, 70.31, 70.20, 70.18, 70.10, 70.08, 70.02, 69.95, 69.89, 68.06, 66.92, 62.72, 40.84, 40.55, 34.53, 33.91, 32.02, 31.75, 31.60, 31.53, 30.28, 30.23, 29.78, 29.75, 29.60, 29.45, 29.25, 29.16, 29.04, 25.70, 22.78, 21.53, 20.20, 17.92, 14.21, 8.73.

**MALDI-TOF-MS (M+H):** 545.49

1.1.1.5. **Compound (1e)**

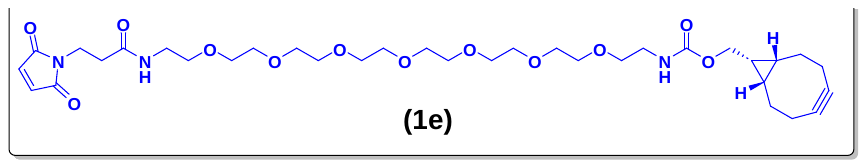

Mol. formula: C_34_H_53_N_3_O_12_

Mol. Weight: 695.80

Physical appearance: Yellowish viscous liquid

Yield: 72%

The **compound (1e)** is synthesized according to **synthetic scheme 1.1**, starting with compound (1d) (0.543 g, 0.997 mmol), maleimide, propionic acid (0.202 g, 1.196 mmol), pyridine (0.015 g, 0.199 mmol), and DCC (0.246 g, 1.196 mmol) in 20 mL of acetonitrile (ACN). The resulting crude mixture was purified by normal-phase chromatography (NPC) using a methanol/dichloromethane (MeOH/DCM) eluent to obtain the pure product (0.499 g, 0.717 mmol), which had an R_f_ value of 0.4 in 5% MeOH/DCM.

**^1^H NMR (400 MHz, CDCl_3_) δ:** 6.75 (s, 1H), 6.69 (s, 2H), 5.36 (s, 1H), 4.14 (d, *J* = 8.0 Hz, 2H), 3.83 (t, *J* = 7.2 Hz, 2H), 3.68 – 3.60 (m, 25H), 3.53 (dt, *J* = 9.8, 5.1 Hz, 4H), 3.39 (dt, *J* = 22.0, 5.0 Hz, 4H), 2.51 (t, *J* = 7.2 Hz, 2H), 2.23 (qd, *J* = 13.2, 2.7 Hz, 6H), 1.65 – 1.51 (m, 2H), 1.26 (d, *J* = 14.0 Hz, 3H).

**^13^C NMR (100 MHz, CDCl_3_) δ:** 170.65, 170.00, 156.98, 134.33, 98.97, 70.68, 70.66, 70.64, 70.61, 70.59, 70.39, 70.30, 70.25, 69.94, 62.80, 40.95, 39.35, 34.63, 34.53, 29.82, 29.49, 29.19, 21.57, 20.24, 17.93.

**MALDI-TOF-MS (M+K):** 734.45

1.2. **Activation of camptothecin**

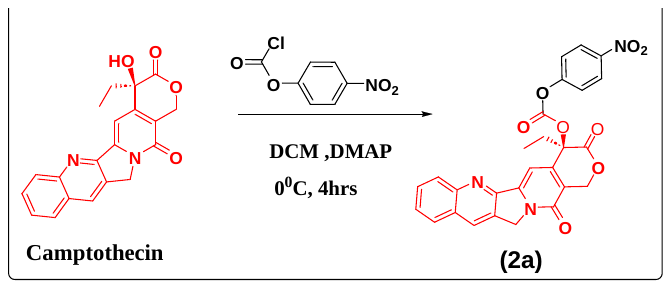

**Scheme 2:** Synthetic scheme for the synthesis of activated camptothecin.

1.2.1. **Compound (2a)**

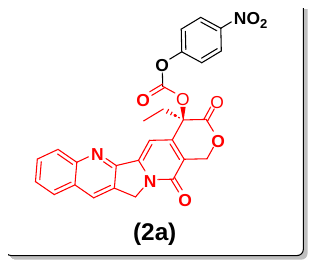

Mol. formula: C_27_H_19_N_3_O_8_

Mol. weight: 513.46

Physical appearance: Yellowish solid

Yield: 62 %

Camptothecin (0.025 g, 0.071 mmol, 1 eq.) and p-nitrophenyl chloroformate (0.050 g, 0.248 mmol, 3.5 eq.) were dissolved in anhydrous DCM at 0 °C, followed by the addition of dimethyl aminopyridine (DMAP) (0.053 g, 0.426 mmol, 6 eq.). After allowing the reaction mixture to proceed for 4 hours, it was washed with 1 N HCl and extracted using DCM. The organic layer was dried over Na_2_SO_4_, concentrated, and purified by normal-phase chromatography with MeOH/DCM as the eluent, yielding a pure product (0.023 g, 0.044 mmol). The R_f_ value was 0.40 in 5% MeOH/DCM.

**^1^H NMR (400 MHz, CDCl_3_) δ:** 8.42 (s, 1H), 8.29 – 8.19 (m, 3H), 7.96 (dd, *J* = 8.2, 1.4 Hz, 1H), 7.86 (ddd, *J* = 8.5, 6.9, 1.5 Hz, 1H), 7.69 (ddd, *J* = 8.1, 6.8, 1.2 Hz, 1H), 7.40 (dd, *J* = 7.2, 2.0 Hz, 3H), 5.71 (d, *J* = 17.3 Hz, 1H), 5.42 (d, *J* = 17.3 Hz, 1H), 5.31 (dd, *J* = 3.9, 1.2 Hz, 2H), 2.31 (ddq, *J* = 52.0, 14.7, 7.4 Hz, 2H), 1.07 (t, *J* = 7.5 Hz, 3H).

**^13^C NMR (100 MHz, CDCl_3_) δ:** 166.98, 157.34, 155.19, 152.28, 151.38, 149.03, 146.96, 145.71, 145.00, 131.50, 131.04, 129.71, 128.59, 128.41, 125.40, 121.85, 120.51, 95.69, 79.45, 67.34, 50.21, 32.03, 29.83, 7.79.

**MALDI-TOF-MS (M+K):** 551.40

1.3. **Synthetic scheme for the synthesis of the CPT-azide**

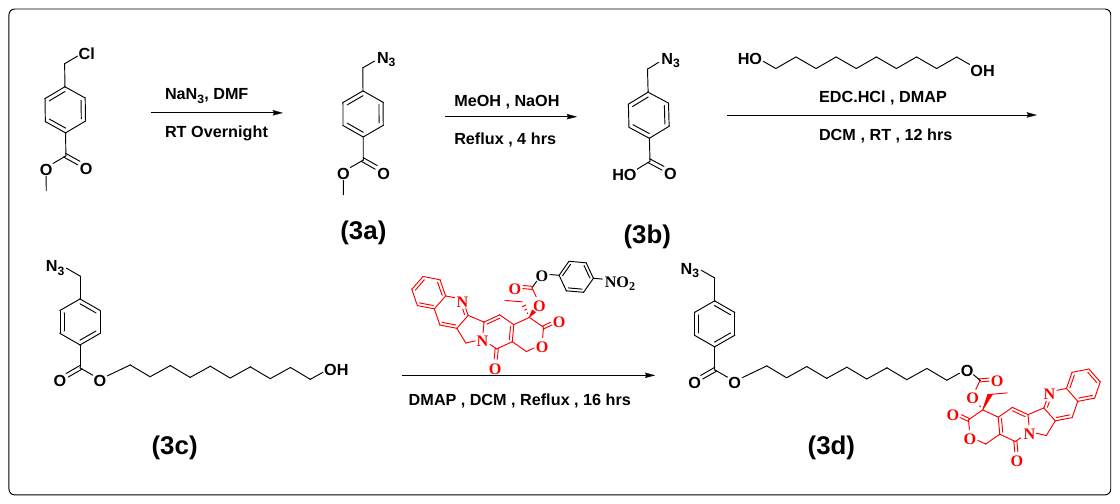

**Scheme 3:** The synthetic scheme for synthesizing the CPT-azide molecule features a hydrophobic part (C10) connected to CPT through a pH-sensitive carbonate bond, allowing for its release in acid-sensitive environments.

The methyl 4-(chloromethyl) benzoate (1 eq.) was dissolved in DMF at ambient temperature. Next, sodium azide (NaN_3_, 5 eq.) was introduced to the reaction mixture, which was permitted to react at room temperature overnight. Once the reaction had concluded, the mixture was concentrated and purified through normal-phase chromatography (NPC) without any further workup, using ethyl acetate and hexane as the eluent to yield compound (3a).

The resulting compound (1 eq.) was hydrolyzed using sodium hydroxide (NaOH, 2 eq.) in methanol (MeOH) under reflux conditions at 80°C for four hours. After the reaction was completed, the methanol was evaporated. Water was added to the residue, and the mixture was acidified with concentrated hydrochloric acid (HCl). Subsequently, extraction was performed using ethyl acetate (EtOAc). The organic layer was collected, dried over sodium sulphate (Na_2_SO_4_), and the crude product was utilized in the subsequent reaction without any additional purification.

In the following step, compound (3b) (1 eq.), decanediol (2 eq.), and DMAP (0.5 eq.) were mixed in dichloromethane (DCM) while stirring. A solution of EDC·HCl (2 eq.) in DCM was gradually added to the above mixture, which was allowed to stir for 12 hours at RT. After the reaction was complete, it was quenched with water, and the resulting mixture was extracted with DCM three times. The combined organic layers were dried over Na_2_SO_4_, concentrated, and purified through NPC using ethyl acetate and hexane to produce compound (3c).

The CPT-azide (3d) was synthesized by dissolving camptothecin-4-nitrophenyl carbonate (2a) (0.5 eq.) and alcohol (3c) (1.2 eq.) in DCM. After the addition of dimethylaminopyridine (DMAP) (0.5 eq.), the reaction mixture was refluxed at 55°C. Following a 12-hour reaction period, the mixture was washed with 1 M sodium bicarbonate (NaHCO_3_) and then extracted with DCM. The organic layer was dried over Na_2_SO_4_, concentrated, and purified by NPC using a mixture of ethyl acetate and hexane, resulting in CPT-azide (3d).

1.3.1. **Synthesis of the CPT-azide and its intermediates**
       1.3.1.1. **Compound (3a)**

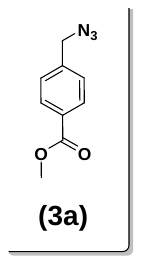

Mol. formula: C_9_H_9_N_3_O_2_

Mol. Weight: 191.19

Physical appearance: white solid

Yield: 69%

The **compound (3a)** is synthesized according to **synthetic scheme 1.3.** The process begins with methyl 4-(chloromethyl) benzoate (7.0 g, 37.91 mmol), sodium azide (NaN_3_) (12.6 g, 189.57 mmol), and dimethylformamide (DMF) (20 mL). The resulting crude material was purified using normal-phase chromatography (NPC) with a solvent mixture of ethyl acetate (EtOAc) and hexane to obtain the pure product (5.0 g, 26.15 mmol), which had an R_f_ value of 0.7 in 15% EtOAc/hexane.

**^1^H NMR (400 MHz, CDCl_3_) δ:** 7.98 – 7.89 (m, 2H), 7.32 – 7.23 (m, 2H), 4.29 (s, 2H), 3.80 (s, 3H).

**^13^C NMR (100 MHz, CDCl_3_) δ:** 166.71, 140.50, 130.20, 130.14, 128.01, 54.35, 52.27.

**MALDI-TOF-MS (M+H):**193.00

1.3.1.2. **Compound (3b)**

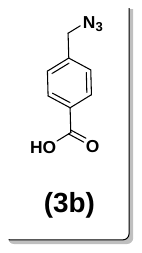

Mol. formula: C_8_H_7_N_3_O_2_

Mol. Weight: 177.16

Physical appearance: yellow solid

Yield: Not calculated

The **compound (3b)** is synthesized according to **synthetic scheme 1.3**, starting with compound (3a) (5.0 g, 26.15 mmol), NaOH (2.09 g, 52.30 mmol), and MeOH (30 mL). The resulting crude material is used in the next step without any purification. The solvent mixture consists of EtOAc/Hexane (4.0 g, 22.57 mmol), with an R_f_ value of 0.3 in 25% EtOAc/Hexane.

1.3.1.3. **Compound (3c)**

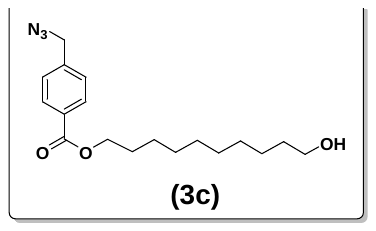

Mol. formula: C_18_H_27_N_3_O_3_

Mol. Weight: 333.43

Physical appearance: Flaky white solid

Yield: 59.8 %

**Compound (3c)** is synthesized according to **synthetic scheme 1.3**, starting with compound (3b) (4.0 g, 22.57 mmol), decandiol (7.86 g, 45.15 mmol), EDC·HCl (8.65 g, 45.15 mmol), DMAP (1.36 g, 11.28 mmol), and dichloromethane (DCM, 30 mL). The resulting crude material was purified using normal-phase chromatography (NPC) with a mixture of ethyl acetate and hexane as the eluent. This process yielded the pure product (4.5 g, 13.49 mmol), which has an Rf value of 0.6 in a 35% ethyl acetate/hexane mixture.

**^1^H NMR (400 MHz, CDCl_3_) δ:** 8.06 – 7.99 (m, 2H), 7.43 – 7.32 (m, 2H), 4.38 (s, 2H), 4.29 (t, *J* = 6.7 Hz, 2H), 3.60 (t, *J* = 6.6 Hz, 2H), 1.79 – 1.70 (m, 2H), 1.53 (p, *J* = 6.7 Hz, 2H), 1.45 – 1.37 (m, 2H), 1.36 – 1.25 (m, 11H).

**^13^C NMR (100 MHz, CDCl_3_) δ:** 166.28, 140.34, 130.42, 130.10, 127.94, 65.31, 62.93, 62.90, 54.29, 32.77, 29.52, 29.45, 29.41, 29.25, 28.70, 26.02, 25.77.

**MALDI-TOF-MS (M+Na):** 356.30

1.3.1.4. **Compound (3d)**

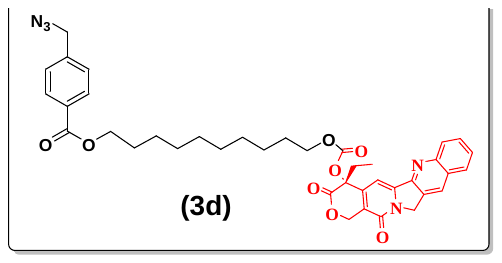

Mol. formula: C_39_H_41_N_5_O_8_

Mol. Weight: 707.78

Physical appearance: Creamy white solid

Yield: 88%

The **compound (3d)** was synthesized following **synthetic scheme 1.3**, starting with compound (3c) (4.5 g, 13.49 mmol), compound (2a) (2.88 g, 5.62 mmol), DMAP (0.687 g, 5.62 mmol), and DCM (30 mL). The resulting crude material was purified using normal-phase chromatography (NPC) with EtOAc/Hexane as the eluent, yielding a pure product (3.5 g, 4.94 mmol) with an R_f_ value of 0.7 in a 70% EtOAc/Hexane mixture.

**^1^H NMR (400 MHz, CDCl_3_) δ:** 8.41 (s, 1H), 8.23 (d, *J* = 8.5 Hz, 1H), 8.06 – 8.02 (m, 2H), 7.94 (dd, *J* = 8.2, 1.4 Hz, 1H), 7.84 (ddd, *J* = 8.5, 6.9, 1.5 Hz, 1H), 7.67 (ddd, *J* = 8.1, 6.8, 1.2 Hz, 1H), 7.38 (d, *J* = 8.5 Hz, 3H), 5.69 (d, *J* = 17.2 Hz, 1H), 5.39 (d, *J* = 17.2 Hz, 1H), 5.35 – 5.23 (m, 2H), 4.41 (s, 2H), 4.28 (t, *J* = 6.7 Hz, 2H), 4.19 – 4.02 (m, 2H), 2.22 (ddt, *J* = 47.1, 14.1, 7.2 Hz, 2H), 1.68 (dt, *J* = 22.4, 7.3 Hz, 4H), 1.41 – 1.31 (m, 4H), 1.24 (d, *J* = 5.3 Hz, 9H), 1.00 (t, *J* = 7.5 Hz, 3H).

**^13^C NMR (100 MHz, CDCl_3_) δ:** 167.61, 166.35, 157.46, 153.98, 152.40, 148.89, 146.40, 146.04, 140.40, 131.46, 130.93, 130.57, 130.21, 129.68, 128.60, 128.36, 128.33, 128.28, 128.06, 126.33, 120.54, 115.81, 96.41, 77.74, 69.39, 67.19, 65.40, 54.43, 50.13, 32.10, 29.83, 29.49, 29.45, 29.30, 29.25, 28.79, 28.64, 26.11, 25.68, 7.78.

**MALDI-TOF-MS (M+Na):** 730.54

1.4. **The general procedure for the synthesis of the alkyated hydrophilic spacer and its intermediates**

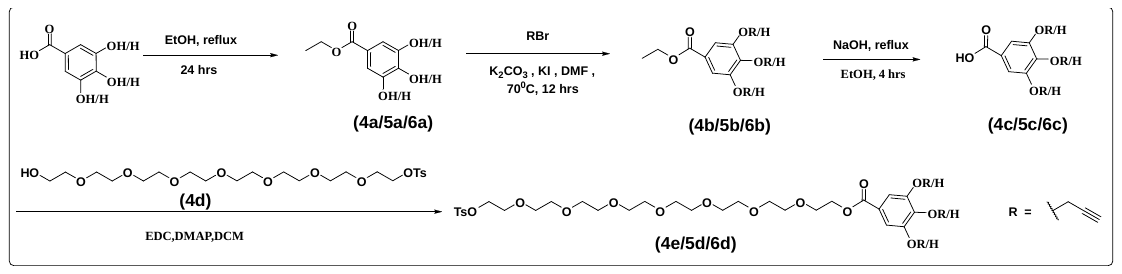

**Scheme 4:** The general synthetic scheme for synthesizing an octaethylene hydrophilic spacer modified with 1T/2T/3T alkyne moieties.

The hydrophilic spacers were synthesized using a hydroxy derivative of benzoic acid. The synthesis begins with esterification using ethanol (EtOH), followed by alkylation with propargyl bromide to produce the alkylated ester. Next, the ester undergoes hydrolysis, and a second esterification is performed with monotosylated octaethylene glycol to obtain the final ester. The detailed methodology for producing the 1T, 2T, and 3T alkyne hydrophilic spacers is outlined below.

**1.4.1.1. Synthesis of ester- procedure A**

The ethyl ester of the hydroxy derivative of benzoic acid was synthesized by refluxing the hydroxy derivative of benzoic acid (1 eq.) in ethanol for 24 hours in the presence of H_2_SO_4_ (2 eq.). Once the reaction was completed, the mixture was neutralized with aqueous sodium bicarbonate solution (NaHCO_3_) and extracted three times with EtOAc to obtain the crude product. This was then dried using Na_2_SO_4_ and purified through silica gel chromatography with EtOAc/hexane as the eluent, resulting in a white solid.

**1.4.1.2. Synthesis of the O-alkylated ester-procedure B**

The obtained ester (1.0 eq.), propargyl bromide (1.5-5 eq.), K_2_CO_3_ (1.5-5 equivalents), and KI (0.05-0.2 eq.) were combined in an oven-dried round-bottom flask (RBF). DMF was added while stirring to dissolve everything, and the mixture was heated at 70 °C for 12 hours. After the reaction was completed, the mixture was neutralized with acidic water and extracted three times with EtOAc. The combined organic phase was dried over Na_2_SO_4_ and evaporated under reduced pressure to yield the crude product, which was then purified by normal-phase chromatography (NPC) using EtOAc/hexane.

**1.4.1.3. Synthesis of the O-alkylated acid-procedure C**

It involves hydrolysing an alkylated ester (1 eq.) using aqueous NaOH (2 eq.) in ethanol under reflux conditions at 90°C for 4 hours. After the reaction was finished, the ethanol was removed, and water was added to the residue, which was then acidified with concentrated hydrochloric acid (HCl). Extraction was performed with ethyl acetate (EtOAc), and the organic layer was collected, dried over sodium sulphate (Na_2_SO_4_), allowing the crude product to be used in the next reaction without further purification.

**1.4.1.4. Synthesis of the O-alkylated** **ester-procedure D**

In the subsequent step, O-alkylated acid (1 eq.), monotosylted OEG (1.2 eq.) (4d), and DMAP (0.5 eq.) were combined in dichloromethane (DCM) while stirring. A solution of EDC·HCl (1.5 eq.) in DCM was gradually added to the mixture, which was stirred for 12 hours at room temperature. Upon completion of the reaction, it was quenched with water, and the resultant mixture was extracted with DCM three times. The combined organic layers were then dried over Na_2_SO_4_, concentrated, and purified via NPC using ethyl acetate and hexane to yield the O-alkylated ester.

**1.4.2. Synthesis of alkyated hydrophilic spacer and its intermediates**

**1.4.2.1. Compound (4a)**

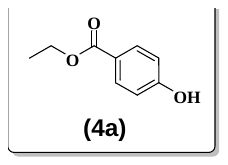

The **compound (4a)** was synthesized using the reported procedure.

**1.4.2.2. Compound (4b)**

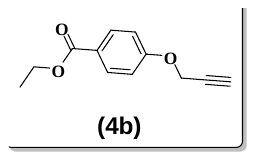

Mol. formula: C_12_H_12_O_3_

Mol. Weight: 204.22

Physical appearance: Yellowish white solid

Yield: 74.1 %

The **compound (4b)** is synthesized using general **procedure B**, starting with compound (4a) (15.09 g, 98.58 mmol), propargyl bromide (17.57 g, 147.88 mmol), K_2_CO_3_ (20.52 g, 147.88 mmol), KI (0.8134 g, 4.9 mmol), and DMF (40 mL). The resulting crude material was purified by normal phase chromatography (NPC) using a mixture of ethyl acetate (EtOAc) and hexane as the eluent. This process yielded a pure product (13.9 g, 73.08 mmol) with an R_f_ value of 0.6 in 50% EtOAc/hexane.

**^1^H NMR (400 MHz, CDCl_3_) δ:** 8.05 – 7.96 (m, 2H), 7.04 – 6.95 (m, 2H), 4.74 (d, *J* = 2.4 Hz, 2H), 3.88 (s, 3H), 2.55 (t, *J* = 2.4 Hz, 1H).

**^13^C NMR (100 MHz, CDCl_3_) δ:** 166.80, 161.26, 131.66, 131.36, 123.56, 114.59, 114.24, 77.93, 76.20, 55.93, 52.03.

**1.4.2.3. Compound (4c)**

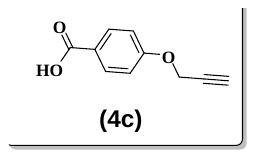

Mol. formula: C_10_H_8_O_3_

Mol. Weight: 176.17

Physical appearance: Pale white solid

Yield: Not calculated

The **compound (4c)** is synthesized using general **procedure C**, starting with compound (4b) (13.9 g, 73.08 mmol), NaOH (5.84 g, 146.16 mmol), and ethanol (30 mL). The resulting crude material was used in the next step without any purification. The crude product obtained (8.0 g, 45.42 mmol) had an R_f_ value of 0.7 in a 70% ethyl acetate/hexane solvent system.

**MALDI-TOF-MS (M+H):** 177.02

**1.4.2.4. Compound (4d)**

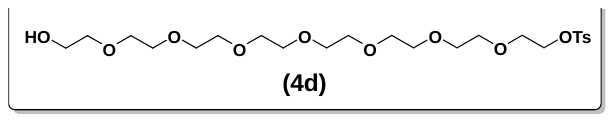

Mol. formula: C_23_H_40_O_11_S

Mol. Weight: 524.62

Physical appearance:

Yield: 42.2%

The **compound (4d)** is synthesized starting with octaethylene glycol (3.0 g, 8.10 mmol, 1 eq.) dissolved in dichloromethane (DCM) while stirring at room temperature. Tosyl chloride (0.772 g, 4.05 mmol, 0.5 eq.) is then added slowly, followed by the addition of DMAP (0.198 g, 1.62 mmol, 0.2 eq.). The reaction mixture is stirred for 10 minutes, after which NEt3 (0.819 g, 8.10 mmol, 1 eq.) is added dropwise. The mixture is stirred at room temperature for 16 hours.

Upon completion of the reaction, water is added to the mixture, and the product is extracted with DCM to obtain the crude product. This crude product is then purified using normal-phase chromatography (NPC) with methanol/DCM as the eluent, yielding a pure product (0.950 g, 1.71 mmol) with an Rf value of 0.4 in 5% methanol/DCM.

**^1^H NMR (400 MHz, CDCl_3_) δ:** 7.78 – 7.70 (m, 2H), 7.29 (d, *J* = 8.0 Hz, 2H), 4.14 – 4.06 (m, 2H), 3.68 – 3.54 (m, 24H), 3.52 (s, 4H), 3.00 (s, 2H), 2.39 (s, 3H).

**^13^C NMR (100 MHz, CDCl_3_) δ:** 144.79, 132.95, 129.82, 127.93, 72.63, 70.67, 70.54, 70.49, 70.46, 70.44, 70.20, 69.27, 68.61, 61.59, 31.87, 29.64, 29.60, 29.30, 22.64, 21.61, 14.10.

**MALDI-TOF-MS (M+K):** 563.29

**1.4.2.5. Compound (4e)**

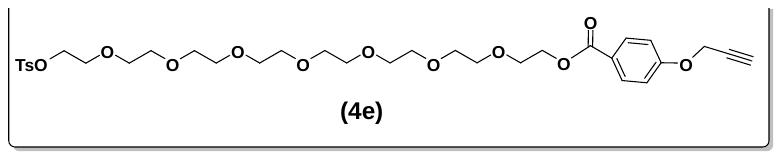

Mol. formula: C_33_H_46_O_13_S

Mol. Weight: 682.77

Physical appearance: pale yellow liquid

Yield: 61.5%

The **compound (4e)** was synthesized using the general **procedure D**, starting with compound (4c) (0.100 g, 0.476 mmol), compound (4d) (0.300 g, 0.572 mmol), EDC·HCl (0.137 g, 0.715 mmol), DMAP (0.029 g, 0.238 mmol), and DCM (30 mL). The resulting crude material was purified by normal phase chromatography (NPC) using a mixture of ethyl acetate (EtOAc) and hexane as the eluent. This purification resulted in the isolation of the pure product (0.200 g, 0.292 mmol) with an R_f_ value of 0.4 in 70% EtOAc/hexane.

**^1^H NMR (400 MHz, CDCl_3_) δ:** 8.02 (d, *J* = 8.9 Hz, 2H), 7.82 – 7.76 (m, 2H), 7.33 (d, *J* = 8.2 Hz, 2H), 7.01 – 6.97 (m, 2H), 4.75 (d, *J* = 2.4 Hz, 2H), 4.44 (dd, *J* = 5.6, 4.1 Hz, 2H), 4.17 – 4.13 (m, 2H), 3.83 – 3.80 (m, 2H), 3.69 – 3.60 (m, 23H), 3.57 (s, 3H), 2.55 (t, *J* = 2.3 Hz, 1H), 2.44 (s, 3H).

**^13^C NMR (100 MHz, CDCl_3_) δ:** 166.27, 161.35, 144.92, 133.15, 131.82, 129.96, 128.12, 123.53, 114.61, 76.26, 70.87, 70.82, 70.77, 70.73, 70.68, 70.64, 69.43, 69.37, 68.81, 64.07, 55.97, 21.78.

**MALDI-TOF-MS (M+Na):** 705.54

**1.4.2.6. Compound (5a)**

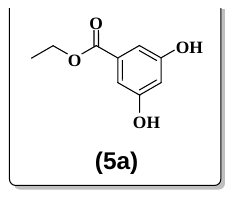

The **compound (5a)** was synthesized using the reported procedure.

**1.4.2.7. Compound (5b)**

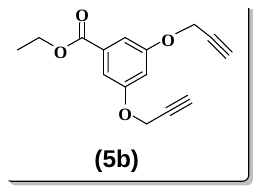

Mol. formula: C_15_H_14_O_4_

Mol. Weight: 258.27

Physical appearance: Fade white solid

Yield: 55.4%

The **compound (5b)** was synthesized using the general **procedure B**, starting with compound (5a) (14.09 g, 76.85 mmol), propargyl bromide (33.09 g, 268.97 mmol), K_2_CO_3_ (37.17 g, 268.97 mmol), KI (0.637 g, 3.84 mmol), and DMF (40 mL). The resulting crude material was then purified by Normal Phase Chromatography (NPC) using EtOAc/Hexane as the eluent. This yielded a pure product (11.0 g, 42.59 mmol), with an R_f_ value of 0.5 in 35% EtOAc/Hexane.

**^1^H NMR (400 MHz, CDCl_3_) δ:** 7.28 (d, *J* = 2.4 Hz, 2H), 6.78 (s, 1H), 4.70 (d, *J* = 2.5 Hz, 4H), 4.35 (q, *J* = 7.1 Hz, 2H), 2.54 (t, *J* = 2.4 Hz, 2H), 1.37 (t, *J* = 7.1 Hz, 3H).

**^13^C NMR (100 MHz, CDCl_3_) δ:** 166.01, 165.47, 158.56, 133.07, 132.62, 108.99, 107.32, 107.07, 106.95, 78.08, 76.59, 76.07, 61.33, 56.98, 56.20, 15.69, 14.36.

**MALDI-TOF-MS (M+H):**259.12

**1.4.2.8. Compound (5c)**

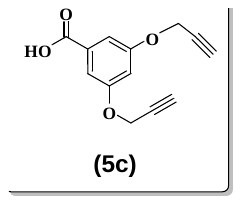

Mol. formula: C_13_H_10_O_4_

Mol. Weight: 230.21

Physical appearance: White solid

Yield: Not calculated

The **compound (5c)** is synthesized using the general **procedure C**, starting with compound (5b) (11 g, 42.59 mmol), NaOH (3.40 g, 85.18 mmol), and ethanol (30 mL). The resulting crude material was used in the next step without any purification. The crude product was obtained (8.0 g, 34.75 mmol) with an R_f_ value of 0.4 in a 70% EtOAc/hexane solution.

**MALDI-TOF-MS (M+Na):** 253.09

**1.4.2.9. Compound (5d)**

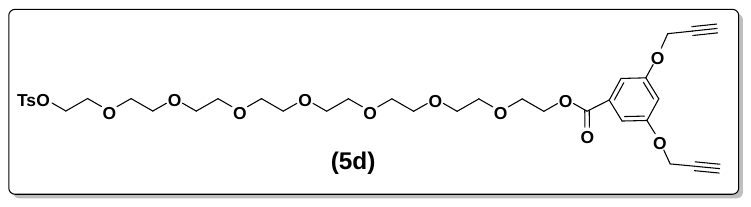

Mol. formula: C_36_H_48_O_14_S

Mol. Weight: 736.82

Physical appearance: Pale yellow liquid

Yield: 26.0%

The **compound (5d)** is synthesized using general **procedure D**, starting with compound (5c) (0.240 g, 1.042 mmol), compound (4d) (0.656 g, 1.25 mmol), EDC·HCl (0.299 g, 1.56 mmol), DMAP (0.063 g, 0.521 mmol), and DCM (30 mL). The resulting crude material was purified by normal-phase chromatography (NPC) using ethyl acetate/hexane as the eluent, yielding the pure product (0.200 g, 0.271 mmol) with an Rf value of 0.3 in a 70% ethyl acetate/hexane mixture.

**^1^H NMR (400 MHz, CDCl_3_) δ:** 7.81 – 7.75 (m, 2H), 7.35 – 7.27 (m, 4H), 6.80 (t, *J* = 2.4 Hz, 1H), 4.71 (d, *J* = 2.5 Hz, 4H), 4.49 – 4.43 (m, 2H), 4.14 (dd, *J* = 5.6, 4.0 Hz, 2H), 3.83 – 3.79 (m, 2H), 3.71 – 3.59 (m, 22H), 3.57 (s, 3H), 2.56 (t, *J* = 2.4 Hz, 2H), 2.43 (s, 3H).

**^13^C NMR (100 MHz, CDCl_3_) δ:** 166.01, 158.61, 144.90, 133.12, 132.26, 129.94, 128.09, 109.20, 107.53, 78.08, 76.20, 70.86, 70.85, 70.74, 70.72, 70.70, 70.67, 70.65, 70.61, 69.36, 69.26, 68.79, 64.61, 56.26, 29.80, 21.76.

**MALDI-TOF-MS (M+Na):** 759.45

**1.4.2.10. Compound (6a)**

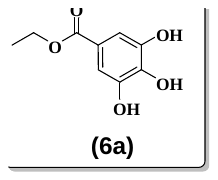

The **compound (6a)** was synthesized using the reported procedure.

**1.4.2.11. Compound (6b)**

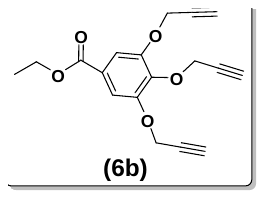

Mol. formula: C_18_H_16_O_5_

Mol. Weight: 312.32

Physical appearance: Yellowish white solid

Yield: 59 %

The **compound (6b)** is synthesized using general **procedure B**, starting with compound (6a) (6.54 g, 33 mmol), propargyl bromide (19.62 g, 165 mmol), potassium carbonate (K_2_CO_3_) (22.80 g, 165 mmol), potassium iodide (KI) (1.095 g, 6.6 mmol), and dimethylformamide (DMF) (40 mL). The resulting crude material was purified by normal phase chromatography (NPC) using a mixture of ethyl acetate (EtOAc) and hexane as the eluent. This process yielded the pure product (6.0 g, 19.50 mmol) with an R_f_ value of 0.5 in 30% EtOAc/hexane.

**^1^H NMR (400 MHz, CDCl_3_) δ:**7.39 (s, 2H), 4.74 (dd, *J* = 3.5, 2.4 Hz, 6H), 4.29 (q, *J* = 7.1 Hz, 2H), 2.58 – 2.41 (m, 3H), 1.31 (t, *J* = 7.1 Hz, 3H).

**^13^C NMR (100 MHz, CDCl_3_) δ:** 165.53, 151.14, 140.88, 125.95, 109.72, 78.63, 77.93, 76.24, 75.66, 61.12, 60.41, 60.13, 56.97, 14.22.

**MALDI-TOF-MS (M+Na):** 335.12

**1.4.2.12. Compound (6c)**

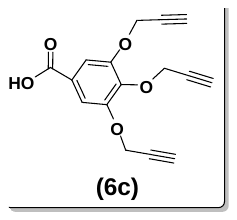

Mol. formula: C_16_H_12_O_5_

Mol. Weight: 284.26

Physical appearance: White solid

Yield: Not calculated

The **compound (6c)** is synthesized using general **procedure C**, starting with compound (6b) (6.0 g, 19.50 mmol), NaOH (1.56 g, 36.0 mmol), and ethanol (30 mL). The resulting crude material was used in the next step without any purification. For the chromatography, a solvent mixture of ethyl acetate (EtOAc) and hexane was used (5.0 g, 17.59 mmol), with an R_f_ value of 0.5 in 70% EtOAc/hexane.

**MALDI-TOF-MS (M+Na):** 307.11

**1.4.2.13. Compound (6d)**

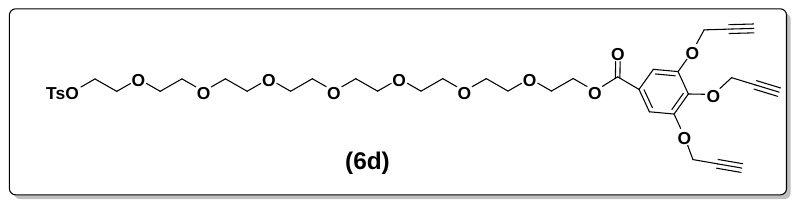

Mol. formula: C_39_H_50_O_15_S

Mol. Weight: 790.87

Physical appearance: Pale yellow liquid

Yield: 26.5 %

The **compound (6d)** is synthesized using general **procedure D**, starting with compound (6c) (0.270 g, 0.950 mmol), compound (4d) (0.597 g, 1.14 mmol), EDC·HCl (0.272 g, 1.42 mmol), DMAP (0.058 g, 0.474 mmol), and DCM (30 mL). The resulting crude material was purified by normal-phase chromatography (NPC) using ethyl acetate (EtOAc) and hexane as the eluents, yielding the pure product (0.200 g, 0.252 mmol) with an R_f_ value of 0.4 in a 70% EtOAc/hexane mixture.

**^1^H NMR (400 MHz, CDCl_3_) δ:** 7.78 (d, *J* = 8.3 Hz, 2H), 7.48 (s, 2H), 7.32 (d, *J* = 8.1 Hz, 2H), 4.80 (dd, *J* = 6.4, 2.4 Hz, 6H), 4.48 – 4.42 (m, 2H), 4.17 – 4.11 (m, 2H), 3.83 – 3.79 (m, 2H), 3.71 – 3.59 (m, 22H), 3.56 (s, 4H), 2.56 (t, *J* = 2.4 Hz, 2H), 2.46 (t, *J* = 2.4 Hz, 1H), 2.43 (s, 3H).

**^13^C NMR (100 MHz, CDCl_3_) δ:** 165.81, 151.38, 144.90, 141.24, 133.05, 129.92, 128.07, 125.81, 110.14, 78.77, 78.07, 76.46, 75.75, 70.82, 70.80, 70.69, 70.65, 70.60, 70.56, 69.35, 69.29, 68.75, 64.62, 60.41, 57.22, 33.91, 32.00, 29.77, 29.74, 29.44, 29.03, 22.77, 21.74, 20.59, 17.51, 14.22.

**MALDI-TOF-MS (M+Na):** 813.47

**1.5. The general procedure for the synthesis of CPT conjugated tail and its intermediates**

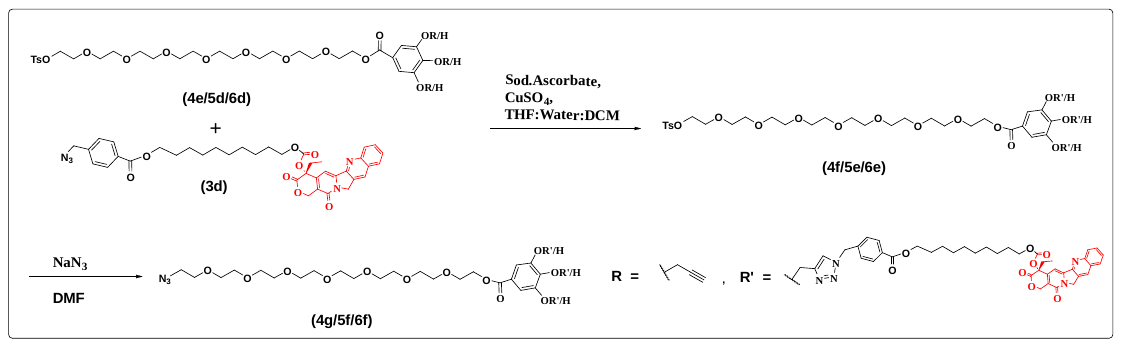

**Scheme 5:** The general synthetic scheme for synthesising the 1T/2T/3T CPT-azide tail utilising 1T/2T/3T alkyne hydrophilic spacers and CPT-azide through click chemistry.

All CPT conjugated tails were synthesised via a [2 + 3] dipolar cycloaddition (click reaction) that involved the use of 1T/2T/3T alkyne hydrophilic spacer and CPT azide. The reaction was facilitated using sodium ascorbate and copper (II) sulphate (CuSO4) in a THF/DCM/H2O mixture at a ratio of 2:1:1. The azidation of tosylated click products was performed using NaN_3_ in DMF. The comprehensive procedure for the synthesis of 1T/2T/3T CPT conjugated tails is provided below.

1.5.1.1. **Synthesis of click product- procedure E**

A hydrophilic alkyne (1 eq.) and CPT azide (2-3.5 eq.) were mixed in a solvent system of tetrahydrofuran (THF) and dichloromethane (DCM). Water was then added, and the mixture was stirred vigorously for an additional 30 minutes, with the ratio of THF, DCM, and water being 2:1:1. Following this, freshly prepared 1 M sodium ascorbate (100 µL) and 1 M copper sulfate (100 µL) were introduced to the reaction mixture, which was allowed to react for 2 hours at room temperature. Upon completion of the reaction, the mixture was extracted with DCM. The organic phase was then dried over sodium sulfate (Na_2_SO_4_) and concentrated under reduced pressure to yield the crude product. The crude product was subsequently purified using normal phase chromatography (NPC) with a methanol (MeOH) and DCM solvent system to obtain the desired clicked product.

1.5.1.2. **Synthesis of CPT-azide tail- procedure F**

The tosylated compound (1 eq.) was dissolved in DMF at room temperature. Sodium azide (NaN_3_, 2-3 eq.) was then added to the reaction mixture, which was allowed to react overnight at room temperature. After the reaction was complete, the mixture was concentrated and purified using normal-phase chromatography (NPC) without any additional processing, employing ethyl acetate and hexane as the eluents to yield the CPT-azide tail.

1.5.2. **Synthesis of CPT conjugated tail and its intermediates**
       1.5.2.1. **Compound (4f)**

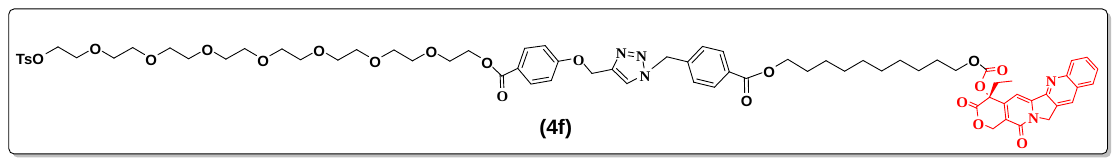

Mol. formula: C_72_H_87_N_5_O_21_S

Mol. Weight: 1390.56

Physical appearance: Yellowish liquid

Yield: 73.7 %

**Compound (4f)** is synthesized using general **procedure E**, starting with compound (4e) (0.200 g, 0.293 mmol) and compound (3d) (0.415 g, 0.586 mmol) and a solution of 1 M sodium ascorbate and 1 M copper (II) sulphate (100 µL of each) is added to a solvent mixture of tetrahydrofuran (THF), dichloromethane (DCM), and water (2:1:1). The crude product obtained is purified using normal-phase chromatography (NPC) with a methanol/dichloromethane (MeOH/DCM) mixture. The final pure product (0.300 g, 0.215 mmol) has an R_f_ value of 0.4 in a 10% MeOH/DCM mixture.

**^1^H NMR (400 MHz, CDCl_3_) δ:** 8.21 (s, 1H), 8.02 (d, *J* = 8.5 Hz, 1H), 7.79 (td, *J* = 15.7, 8.1 Hz, 5H), 7.69 – 7.56 (m, 4H), 7.47 (t, *J* = 7.5 Hz, 1H), 7.21 – 7.11 (m, 5H), 6.81 (d, *J* = 8.8 Hz, 2H), 5.51 (d, *J* = 17.1 Hz, 1H), 5.46 (s, 2H), 5.22 (d, *J* = 17.0 Hz, 1H), 5.11 – 4.92 (m, 4H), 4.24 (dd, *J* = 6.0, 3.7 Hz, 2H), 4.09 (t, *J* = 6.6 Hz, 2H), 4.01 – 3.89 (m, 4H), 3.64 (dd, *J* = 5.8, 3.9 Hz, 2H), 3.57 – 3.34 (m, 26H), 2.05 (ddt, *J* = 37.2, 14.2, 7.3 Hz, 2H), 1.50 (dp, *J* = 21.8, 6.8 Hz, 4H), 1.31 – 1.09 (m, 13H), 0.86 (t, *J* = 7.4 Hz, 3H).

**^13^C NMR (100 MHz, CDCl_3_) δ:** 167.12, 165.57, 165.40, 161.59, 156.75, 153.40, 152.70, 151.81, 148.26, 146.05, 145.43, 144.43, 143.38, 139.28, 138.73, 132.53, 131.26, 130.96, 130.31, 130.21, 129.77, 129.48, 129.06, 128.23, 127.93, 127.77, 127.58, 127.49, 124.33, 123.98, 123.58, 123.24, 122.91, 122.55, 119.63, 115.00, 113.96, 113.78, 95.39, 77.36, 77.30, 70.88, 70.23, 70.19, 70.16, 70.14, 70.11, 70.05, 69.02, 68.81, 68.74, 68.19, 66.57, 64.83, 63.50, 63.34, 61.51, 53.15, 49.60, 34.57, 34.43, 34.11, 34.03, 33.92, 33.76, 33.58, 33.39, 31.49, 31.32, 31.13, 31.05, 29.94, 29.82, 29.77, 29.26, 29.23, 29.19, 29.08, 28.93, 28.89, 28.84, 28.72, 28.67, 28.66, 28.50, 28.19, 28.11, 25.51, 25.12, 24.56, 22.27, 21.22, 13.78, 7.32.

**MALDI-TOF-MS (M+K):** 1428.92

**1.5.2.2. Compound (4g)**

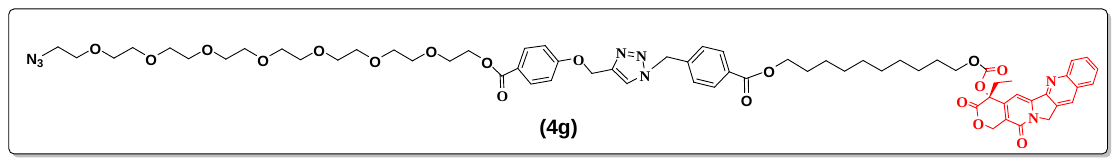

Mol. formula: C_65_H_80_N_8_O_18_

Mol. Weight: 1261.39

Physical appearance: Yellow liquid

Yield: 43.5 %

The **compound (4g)** is synthesized according to general **procedure F**, starting with compound (4f) (0.300 g, 0.273 mmol), sodium azide (NaN_3_) (0.054 g, 0.546 mmol), and dimethylformamide (DMF) (10 mL). The resulting crude product was purified by normal-phase chromatography (NPC) using a mixture of methanol (MeOH) and dichloromethane (DCM) to obtain the pure product (0.150 g, 0.119 mmol), with an R_f_ value of 0.5 in a 10% MeOH/DCM solution.

**^1^H NMR (400 MHz, CDCl_3_) δ:** 8.25 (s, 1H), 8.07 (d, *J* = 8.5 Hz, 1H), 7.84 (td, *J* = 16.8, 15.5, 8.1 Hz, 5H), 7.69 (t, *J* = 7.7 Hz, 1H), 7.64 (d, *J* = 2.6 Hz, 1H), 7.52 (t, *J* = 7.5 Hz, 1H), 7.21 (d, *J* = 7.3 Hz, 3H), 6.88 – 6.83 (m, 2H), 5.55 (d, *J* = 17.1 Hz, 1H), 5.50 (s, 2H), 5.26 (d, *J* = 17.1 Hz, 1H), 5.08 (d, *J* = 13.3 Hz, 4H), 4.33 – 4.25 (m, 2H), 4.13 (t, *J* = 6.6 Hz, 2H), 4.05 – 3.88 (m, 3H), 3.68 (t, *J* = 4.8 Hz, 2H), 3.57 – 3.49 (m, 23H), 3.43 (s, 2H), 3.24 (t, *J* = 5.0 Hz, 1H), 2.09 (ddq, *J* = 47.0, 14.7, 7.3 Hz, 2H), 1.53 (dq, *J* = 22.1, 7.0 Hz, 4H), 1.31 (s, 2H), 1.29 – 1.14 (m, 13H), 0.89 (t, *J* = 7.5 Hz, 3H).

**^13^C NMR (100 MHz, CDCl_3_) δ:** 167.21, 165.71, 165.53, 161.66, 161.31, 156.90, 153.51, 152.74, 151.89, 148.37, 146.10, 145.59, 144.52, 143.57, 141.33, 139.23, 138.85, 135.85, 135.80, 134.82, 132.64, 131.38, 131.07, 130.88, 130.50, 130.37, 129.92, 129.57, 129.17, 128.29, 128.01, 127.88, 127.73, 127.60, 127.58, 124.44, 123.40, 123.17, 123.15, 123.00, 122.69, 119.79, 115.52, 115.10, 114.05, 113.86, 95.64, 77.37, 77.36, 70.99, 70.34, 70.31, 70.30, 70.28, 70.23, 69.69, 69.07, 68.93, 68.86, 68.30, 66.67, 64.97, 64.56, 64.26, 63.60, 61.63, 53.32, 50.33, 49.72, 42.54, 36.17, 35.86, 34.53, 34.01, 33.86, 33.68, 33.49, 32.49, 31.60, 31.46, 31.41, 31.15, 30.71, 30.03, 29.91, 29.37, 29.33, 29.29, 29.18, 29.12, 29.03, 28.99, 28.94, 28.83, 28.77, 28.76, 28.61, 28.29, 28.21, 26.72, 25.61, 25.22, 24.65, 22.37, 21.32, 13.87, 7.40.

**MALDI-TOF-MS (M+Na):** 1284.03

**1.5.2.3. Compound (5e)**

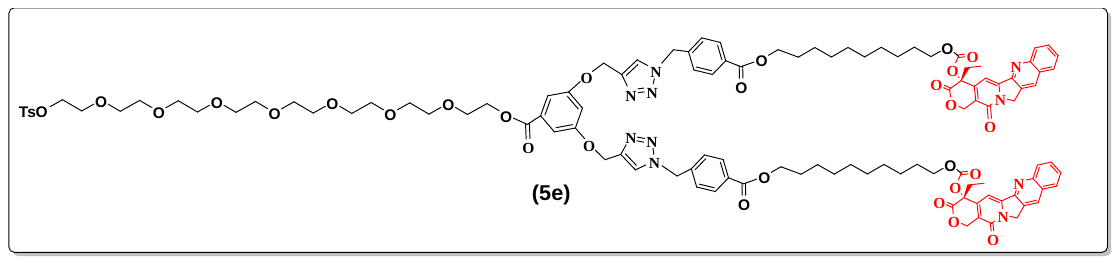

Mol. formula: C_114_H_130_N_10_O_30_S

Mol. Weight: 2152.39

Physical appearance: Yellowish liquid

Yield: 56.8 %

The **compound (5e)** is synthesized using general **procedure E**, starting with compound (5d) (0.150 g, 0.204 mmol) and compound (3d) (0.360 g, 0.509 mmol) and a solution of 1 M sodium ascorbate and 1 M copper (II) sulphate (100 µL of each) is added to a solvent mixture of tetrahydrofuran (THF), dichloromethane (DCM), and water (2:1:1). The resulting crude material is purified by normal phase chromatography (NPC) using a mixture of methanol (MeOH) and DCM to obtain the pure product (0.250 g, 0.116 mmol). The R_f_ value is 0.4 in a 10% MeOH/DCM mixture.

**^1^H NMR (400 MHz, CDCl_3_) δ:** 8.26 (s, 2H), 8.07 (d, *J* = 8.5 Hz, 2H), 7.89 (d, *J* = 7.7 Hz, 4H), 7.80 (d, *J* = 8.2 Hz, 2H), 7.74 – 7.60 (m, 6H), 7.52 (t, *J* = 7.5 Hz, 2H), 7.21 (q, *J* = 6.0, 4.7 Hz, 8H), 7.17 – 7.10 (m, 2H), 6.68 (d, *J* = 9.3 Hz, 1H), 5.71 – 5.46 (m, 6H), 5.27 (d, *J* = 17.0 Hz, 2H), 5.16 – 4.98 (m, 8H), 4.35 – 4.27 (m, 2H), 4.15 (t, *J* = 6.6 Hz, 4H), 4.05 – 3.95 (m, 6H), 3.69 (t, *J* = 5.0 Hz, 3H), 3.57 – 3.48 (m, 23H), 3.44 (s, 4H), 2.31 (s, 3H), 2.22 – 1.98 (m, 5H), 1.56 (dt, *J* = 22.4, 7.3 Hz, 8H), 1.22 (dd, *J* = 17.6, 8.1 Hz, 25H), 0.90 (t, *J* = 7.4 Hz, 6H).

**^13^C NMR (100 MHz, CDCl_3_) δ:** 173.03, 167.26, 165.60, 165.58, 158.96, 158.28, 156.96, 153.58, 152.78, 151.97, 151.89, 148.46, 146.19, 145.64, 144.57, 143.70, 141.42, 139.36, 138.90, 135.82, 134.89, 132.70, 131.85, 131.08, 130.94, 130.71, 130.51, 130.38, 129.97, 129.61, 129.24, 128.57, 128.34, 128.06, 127.93, 127.76, 127.66, 127.61, 124.50, 124.26, 123.73, 123.47, 123.26, 123.07, 119.85, 115.56, 113.90, 108.69, 108.56, 106.73, 106.44, 95.67, 77.84, 77.36, 76.07, 70.40, 70.33, 70.31, 70.25, 70.21, 69.13, 68.92, 68.81, 68.36, 66.74, 65.26, 65.01, 64.62, 64.33, 64.19, 64.12, 61.87, 55.84, 53.34, 49.78, 36.23, 34.58, 34.27, 34.06, 33.92, 33.85, 33.76, 33.56, 31.66, 31.52, 31.45, 31.28, 31.20, 30.78, 30.31, 30.08, 29.97, 29.92, 29.44, 29.40, 29.37, 29.32, 29.25, 29.10, 29.06, 29.01, 28.90, 28.84, 28.82, 28.68, 28.40, 28.36, 28.27, 26.78, 25.67, 25.28, 24.71, 22.44, 22.07, 21.37, 18.94, 13.92, 13.85, 13.51, 7.45.

**MALDI-TOF-MS (M+Na):** 2175.65

**1.5.2.4. Compound (5f)**

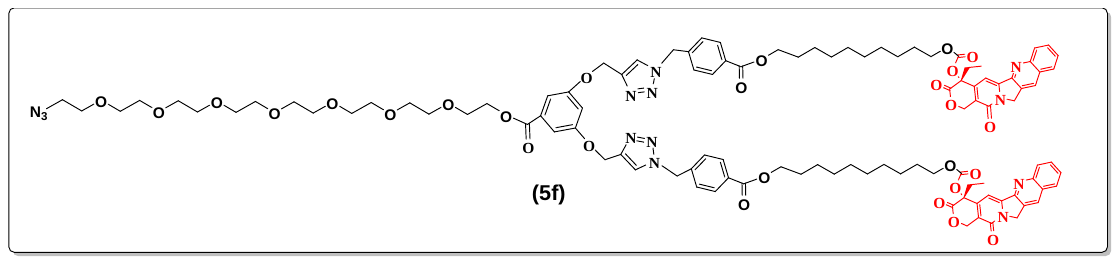

Mol. formula: C_107_H_123_N_13_O_27_

Mol. Weight: 2023.22

Physical appearance: Yellowish waxy solid

Yield: 72.41 %

The **compound (5f)** is synthesized using the general **procedure F**, starting with compound (5e) (0.250 g, 0.116 mmol), sodium azide (NaN_3_) (0.023 g, 0.348 mmol), and dimethylformamide (DMF) (10 mL). The resulting crude material was purified by normal phase chromatography (NPC) using a methanol (MeOH) and dichloromethane (DCM) mixture to obtain the pure product (0.170 g, 0.084 mmol), with an R_f_ value of 0.5 in 10% MeOH/DCM.

**^1^H NMR (400 MHz, CDCl_3_) δ:** 8.32 (s, 2H), 8.14 (d, *J* = 8.5 Hz, 2H), 7.95 (d, *J* = 7.7 Hz, 4H), 7.86 (d, *J* = 8.1 Hz, 2H), 7.81 – 7.45 (m, 8H), 7.28 (d, *J* = 5.7 Hz, 4H), 7.23 – 7.14 (m, 3H), 5.61 (d, *J* = 17.1 Hz, 2H), 5.55 (s, 4H), 5.32 (d, *J* = 17.1 Hz, 2H), 5.18 – 5.05 (m, 7H), 4.40 – 4.32 (m, 2H), 4.20 (t, *J* = 6.7 Hz, 4H), 4.14 – 3.94 (m, 6H), 3.73 (t, *J* = 4.9 Hz, 3H), 3.57 (dq, *J* = 14.1, 5.3, 3.3 Hz, 23H), 3.49 (s, 2H), 3.30 (t, *J* = 4.9 Hz, 2H), 2.28 – 2.05 (m, 5H), 1.61 (dq, *J* = 22.5, 7.1 Hz, 10H), 1.40 – 1.23 (m, 25H), 0.94 (t, *J* = 7.4 Hz, 6H).

**^13^C NMR (100 MHz, CDCl_3_) δ:** 175.89, 171.78, 167.38, 165.76, 159.06, 159.02, 158.40, 157.14, 153.70, 152.76, 152.08, 148.57, 146.99, 146.23, 145.79, 145.34, 144.69, 143.91, 141.69, 139.32, 139.07, 138.80, 135.87, 135.03, 132.82, 132.00, 131.26, 130.72, 130.59, 130.15, 129.73, 129.35, 128.70, 128.42, 128.17, 128.07, 127.95, 127.80, 127.74, 124.62, 124.37, 124.25, 123.85, 123.65, 123.37, 123.22, 120.06, 118.94, 118.72, 115.67, 113.99, 108.83, 108.71, 106.92, 106.63, 95.97, 77.91, 77.52, 77.36, 76.10, 70.55, 70.52, 70.50, 70.46, 70.44, 70.39, 70.34, 69.87, 69.21, 69.07, 68.96, 68.51, 66.88, 65.41, 65.18, 64.79, 64.48, 64.31, 64.25, 62.06, 62.02, 55.98, 53.55, 50.53, 49.92, 36.37, 34.85, 34.74, 34.69, 34.39, 34.31, 34.18, 34.05, 33.85, 33.68, 32.48, 31.78, 31.70, 31.55, 31.39, 31.32, 30.90, 30.20, 30.08, 30.03, 29.55, 29.52, 29.48, 29.37, 29.22, 29.18, 29.14, 29.07, 29.02, 28.97, 28.95, 28.80, 28.48, 28.38, 26.91, 25.79, 25.40, 24.80, 22.56, 21.51, 19.05, 14.02, 13.62, 7.55.

**MALDI-TOF-MS (M+Na):** 2046.45

**1.5.2.5. Compound (6e)**

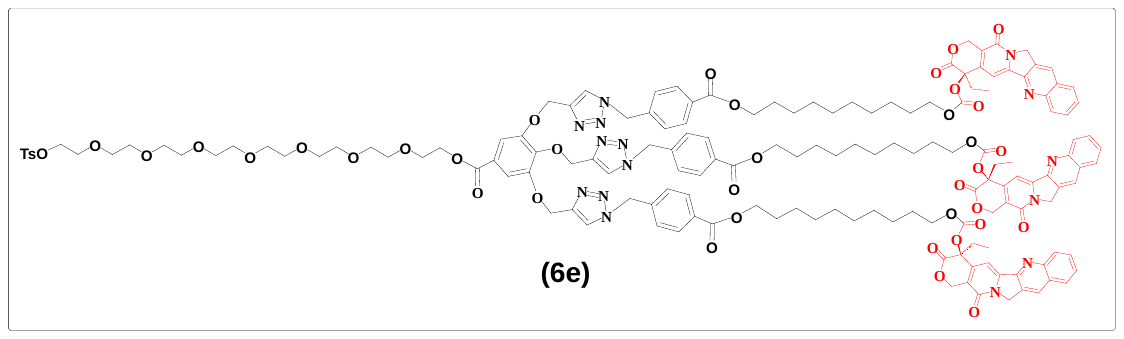

Mol. formula: C_156_H_173_N_15_O_39_S

Mol. Weight: 2914.22

Physical appearance: Yellowish liquid

Yield: 81.7 %

The **compound (6e)** is synthesized following general **procedure E**, starting with compound (6d) (0.100 g, 0.126 mmol) and compound (3d) (0.313 g, 0.442 mmol) and solution of 1 M sodium ascorbate and 1 M copper (II) sulphate (100 µL of each) is added to a solvent mixture of tetrahydrofuran (THF), dichloromethane (DCM), and water (2:1:1). The resulting crude material was purified by normal phase chromatography (NPC) using a methanol/dichloromethane (MeOH/DCM) mixture to yield the pure product (0.300 g, 0.1029 mmol) with an R_f_ value of 0.4 in a 10% MeOH/DCM mixture.

**^1^H NMR (400 MHz, CDCl_3_) δ:** 8.36 (s, 3H), 8.17 (d, *J* = 8.5 Hz, 3H), 8.03 – 7.57 (m, 21H), 7.29 (d, *J* = 7.1 Hz, 12H), 5.71 – 5.46 (m, 9H), 5.35 (d, *J* = 17.2 Hz, 3H), 5.20 (p, *J* = 16.9, 14.7 Hz, 10H), 4.89 (d, *J* = 10.1 Hz, 2H), 4.44 – 4.38 (m, 2H), 4.22 (t, *J* = 6.7 Hz, 7H), 4.14 – 4.01 (m, 9H), 3.78 (t, *J* = 4.9 Hz, 2H), 3.67 – 3.55 (m, 23H), 3.53 (s, 4H), 2.40 (s, 3H), 2.26 – 2.09 (m, 7H), 1.64 (dt, *J* = 18.7, 7.2 Hz, 13H), 1.41 – 1.23 (m, 37H), 0.97 (t, *J* = 7.4 Hz, 9H).

**^13^C NMR (100 MHz, CDCl_3_) δ:** 167.50, 165.92, 165.90, 165.67, 157.28, 153.82, 152.32, 148.84, 146.42, 145.82, 144.81, 139.23, 132.95, 132.62, 131.26, 130.87, 130.74, 130.68, 130.25, 130.11, 129.84, 129.60, 128.49, 128.25, 128.18, 128.06, 127.97, 127.95, 125.70, 120.24, 114.06, 109.47, 95.96, 77.63, 77.36, 70.70, 70.62, 70.59, 70.57, 70.52, 70.48, 69.30, 69.21, 69.14, 68.65, 67.05, 65.30, 65.27, 64.37, 49.99, 33.81, 31.91, 30.32, 29.68, 29.64, 29.60, 29.49, 29.42, 29.34, 29.32, 29.28, 29.14, 29.11, 29.08, 28.93, 28.61, 28.50, 25.91, 25.53, 22.68, 21.64, 14.13, 7.66.

**MALDI-TOF-MS (M+Na):** 2938.21

**1.5.2.6. Compound (6f)**

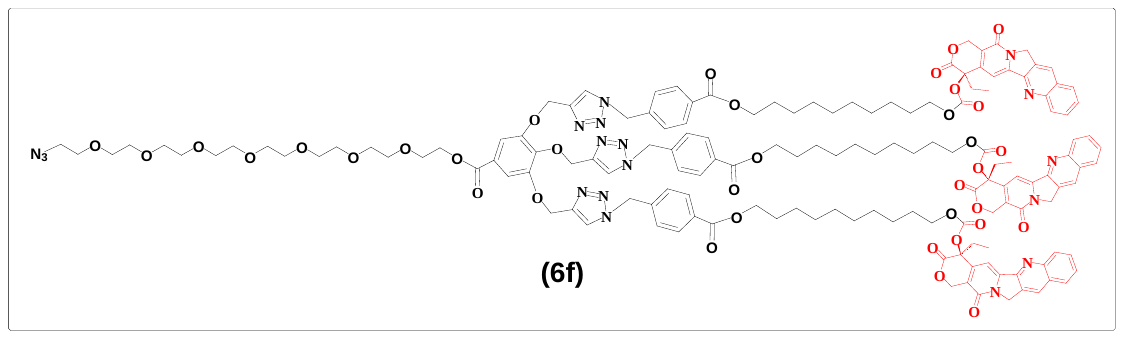

Mol. formula: C_149_H_166_N_18_O_36_

Mol. Weight: 2785.05

Physical appearance: Yellowish solid

Yield: 62.1 %

The **compound (6f)** was synthesized using general **procedure F**, starting with compound (6e) (0.300 g, 0.103 mmol), sodium azide (NaN_3_) (0.021 g, 0.308 mmol), and DMF (10 mL). The resulting crude material was purified by normal phase chromatography (NPC) using a methanol/dichloromethane (MeOH/DCM) solvent system to obtain the pure product (0.180 g, 0.064 mmol) with an R_f_ value of 0.5 in 10% MeOH/DCM.

**^1^H NMR (400 MHz, CDCl_3_) δ:** 8.33 (s, 3H), 8.16 (d, *J* = 8.5 Hz, 3H), 7.98 – 7.84 (m, 10H), 7.75 (dt, *J* = 16.5, 7.7 Hz, 4H), 7.60 (t, *J* = 7.5 Hz, 3H), 7.35 – 7.27 (m, 9H), 7.24 (d, *J* = 2.5 Hz, 2H), 6.96 (dd, *J* = 8.2, 2.6 Hz, 2H), 5.64 (d, *J* = 17.2 Hz, 3H), 5.52 (d, *J* = 13.4 Hz, 6H), 5.34 (d, *J* = 17.2 Hz, 3H), 5.24 – 5.13 (m, 10H), 4.96 – 4.86 (m, 2H), 4.40 (t, *J* = 4.5 Hz, 2H), 4.21 (t, *J* = 6.7 Hz, 6H), 4.05 (tt, *J* = 10.7, 4.6 Hz, 8H), 3.77 (t, *J* = 4.9 Hz, 2H), 3.65 – 3.54 (m, 22H), 3.51 (s, 2H), 2.30 – 2.06 (m, 7H), 1.62 (dp, *J* = 21.7, 6.9 Hz, 14H), 1.31 – 1.23 (m, 37H), 0.95 (t, *J* = 7.5 Hz, 9H).

**^13^C NMR (100 MHz, CDCl_3_) δ:** 176.24, 167.45, 165.90, 165.86, 165.61, 157.25, 153.78, 152.79, 152.76, 152.15, 152.07, 151.79, 148.67, 146.30, 145.92, 145.40, 144.74, 144.47, 143.78, 141.85, 141.47, 139.72, 139.40, 139.14, 135.91, 135.12, 132.88, 131.28, 131.09, 130.73, 130.66, 130.59, 130.19, 130.08, 129.78, 129.44, 128.46, 128.20, 128.13, 128.02, 127.87, 127.84, 127.74, 127.69, 125.62, 124.69, 124.48, 124.30, 123.75, 123.43, 123.28, 120.14, 118.79, 115.73, 114.04, 109.37, 96.11, 77.57, 77.36, 70.61, 70.56, 70.52, 70.50, 70.44, 69.93, 69.26, 69.14, 69.07, 68.57, 66.94, 66.13, 65.24, 65.21, 64.57, 64.32, 62.99, 53.59, 53.38, 50.58, 50.00, 36.44, 34.91, 34.75, 34.37, 34.24, 34.12, 33.94, 33.75, 32.54, 31.85, 31.77, 31.61, 31.45, 31.38, 30.96, 30.39, 30.25, 30.14, 30.09, 29.62, 29.59, 29.55, 29.44, 29.29, 29.26, 29.22, 29.14, 29.08, 29.05, 29.02, 28.87, 28.55, 28.44, 26.97, 26.65, 25.85, 25.47, 24.86, 22.62, 22.27, 21.56, 20.96, 14.08, 7.60.

**MALDI-TOF-MS (M+Na):** 2808.26

**2. Cellular Studies**

**2.1. Protocol for Mammalian Cell Culture: MCF7 cells**

**2.1.1. Cell Seeding:**

MCF7 cells were cultivated until reaching 40-50% confluency. The media was discarded from the flask, and the cells were washed with 1X PBS. To diminish cell adhesion, 0.05% Trypsin was applied to the flask for 30-40 seconds at 37°C. The cells were centrifuged at 300 rpm for 3-5 minutes at room temperature. The supernatant was removed, and 1 mL of DMEM was added and mixed thoroughly. 900 µL of this mixture was diluted in 20 mL of media, which was subsequently used to seed the cells for the experiment. 100 µL was reserved for growth to maintain the culture.

**2.1.2. Cell Splitting:**

When the cell confluency reaches 80%, the media is removed, and the plate is rinsed with 1 mL of 1X PBS. A 500 µL solution of trypsin-EDTA was added to the plate and incubated for 1 minute at 37 °C. Following this, 1 mL of media (DMEM) was added to the plate and washed using a pipette. The media was poured into a 15 mL Falcon tube and centrifuged for 3 minutes at 300 g at room temperature. The supernatant was discarded, and the pellet was resuspended in 500 µL of media. This resuspended pellet was transferred to a petri dish containing 9.5 mL of media and incubated at 37 °C.

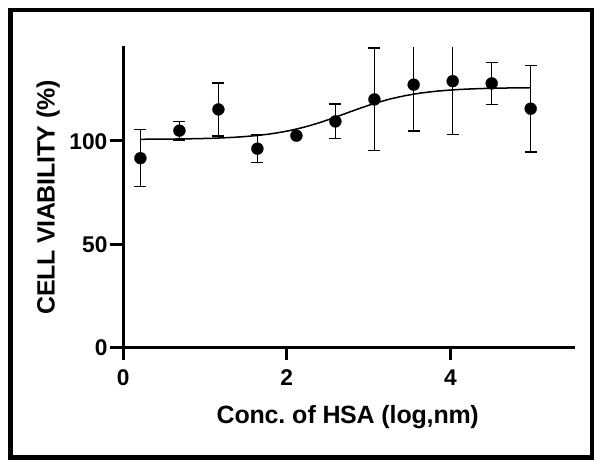

**Figure 1:** Cytotoxicity assessments of HSA in the MCF7 cell line were carried out after a 72-hour incubation at 37°C.

**3. NMR data**

**Compound (1a)**

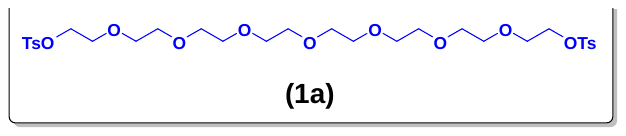

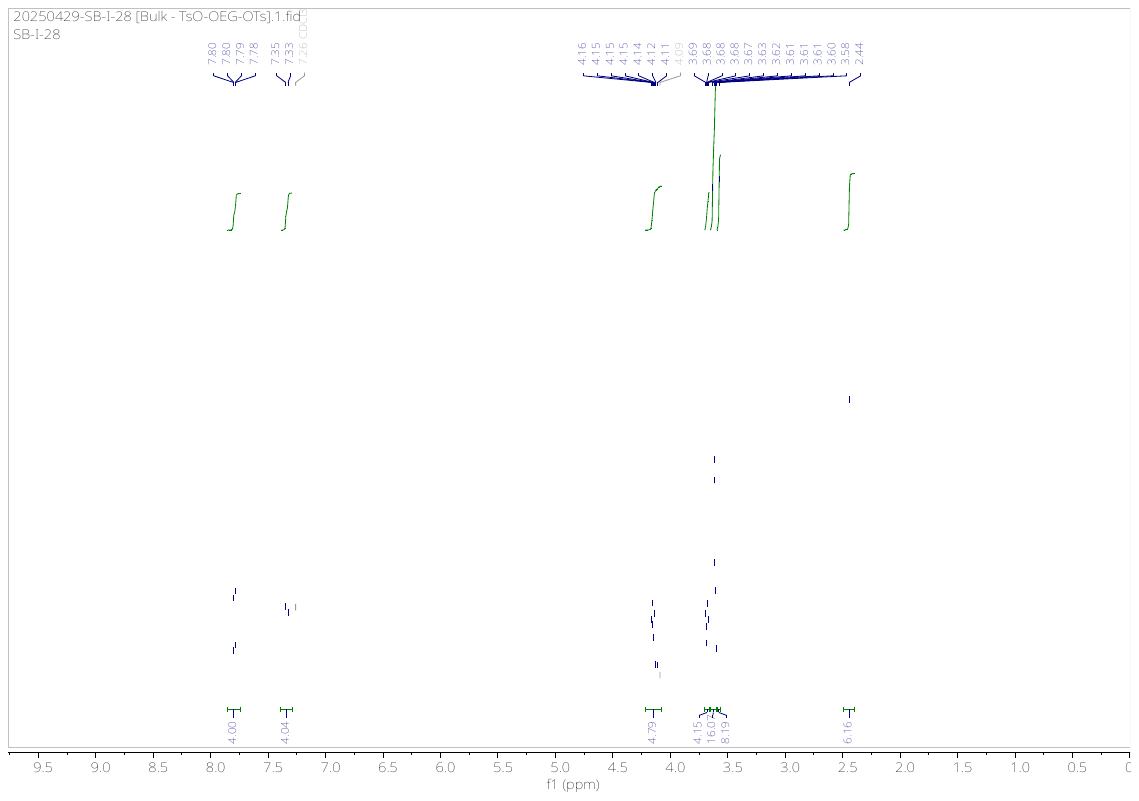

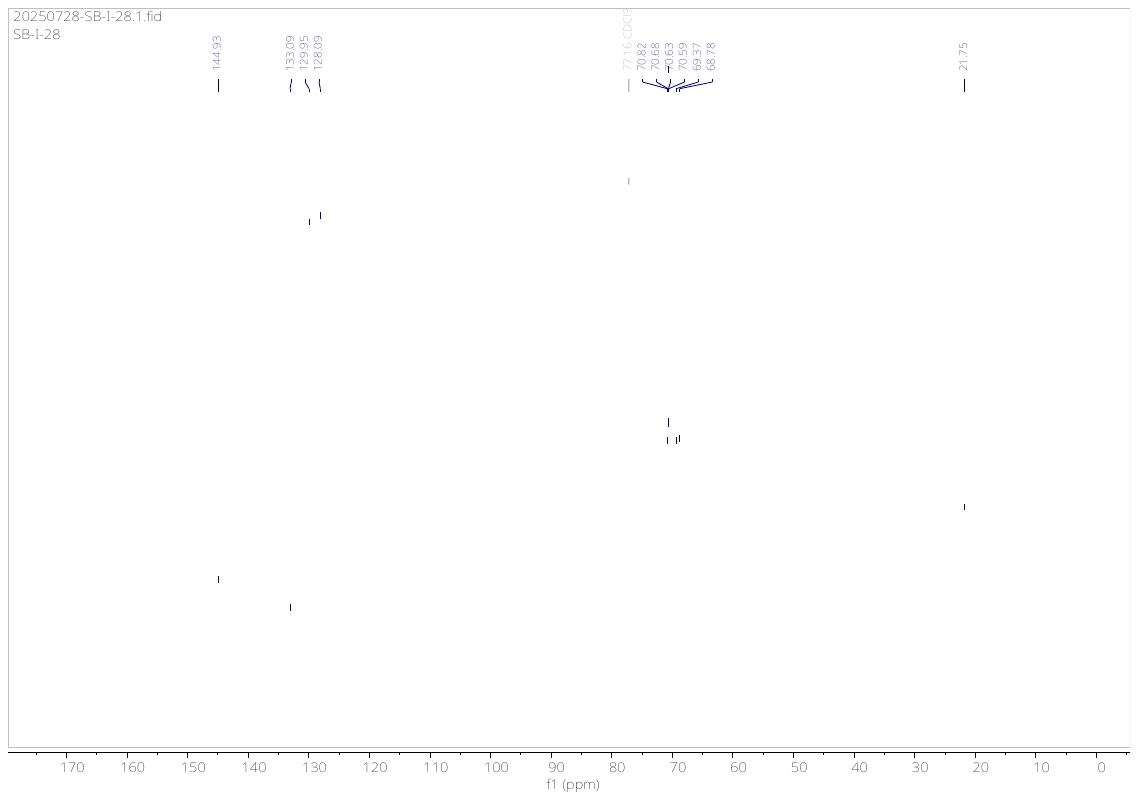

**Compound (1b)**

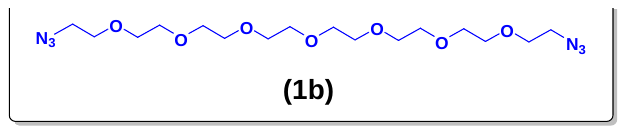

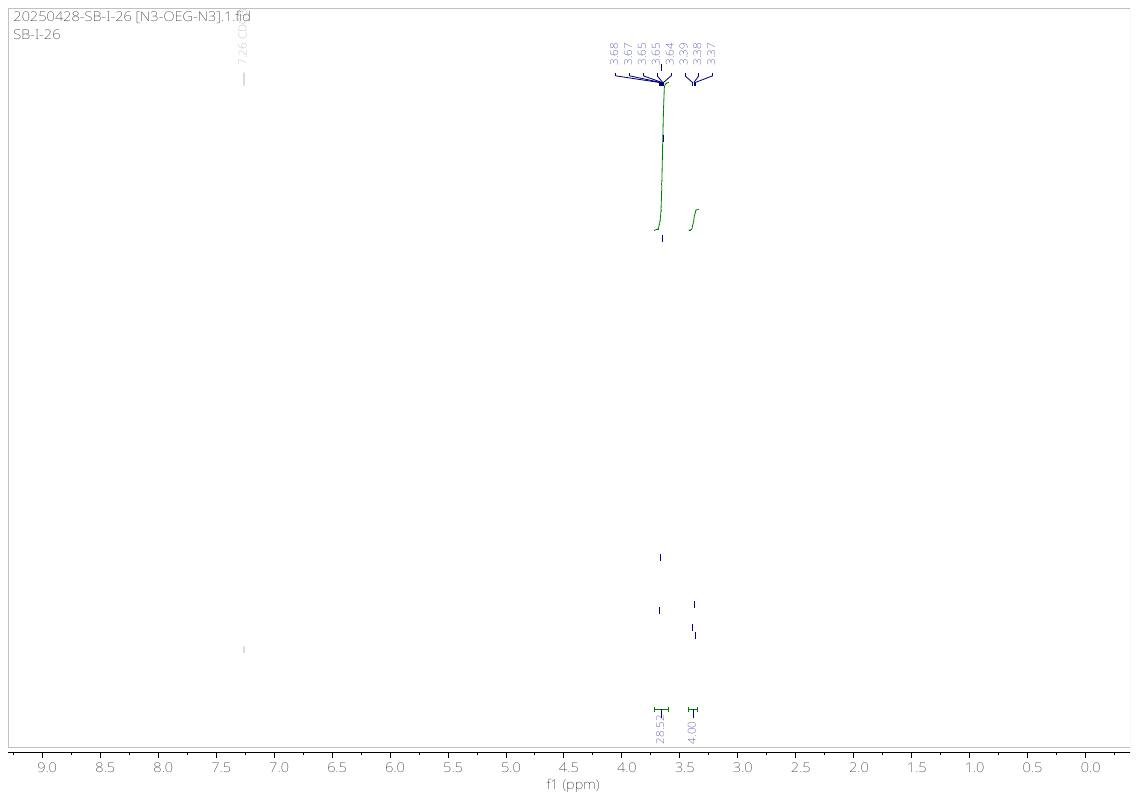

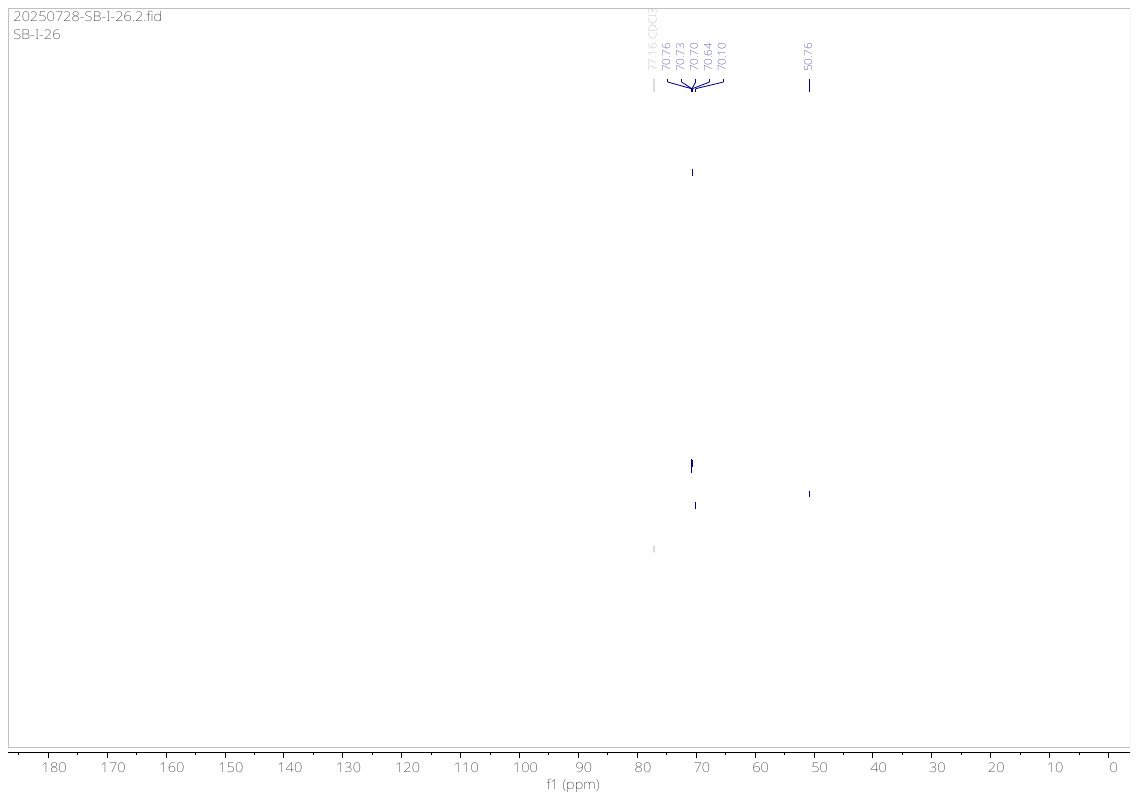

**Compound (1c)**

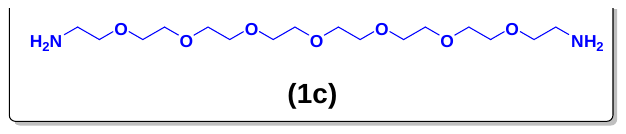

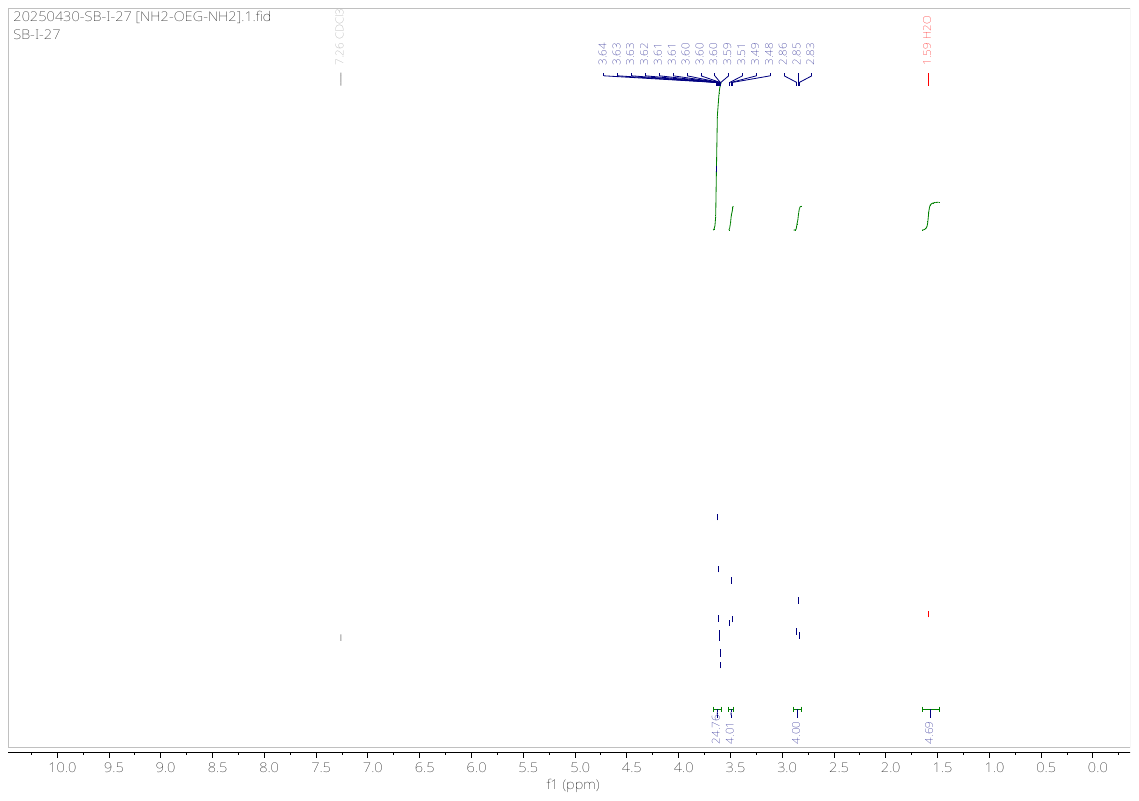

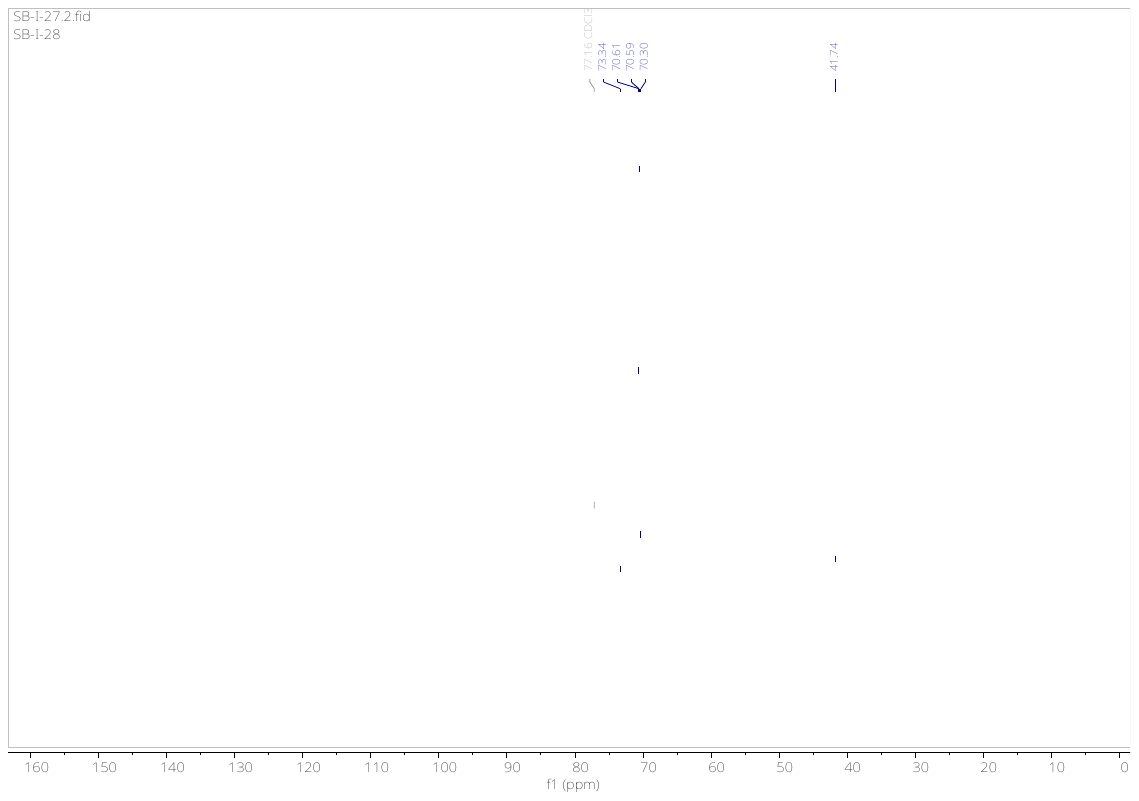

**Compound (1d)**

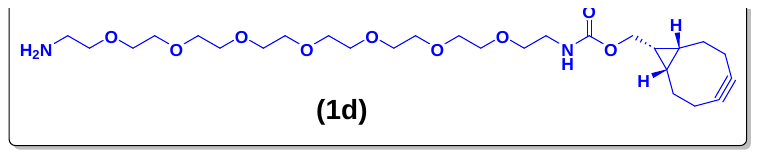

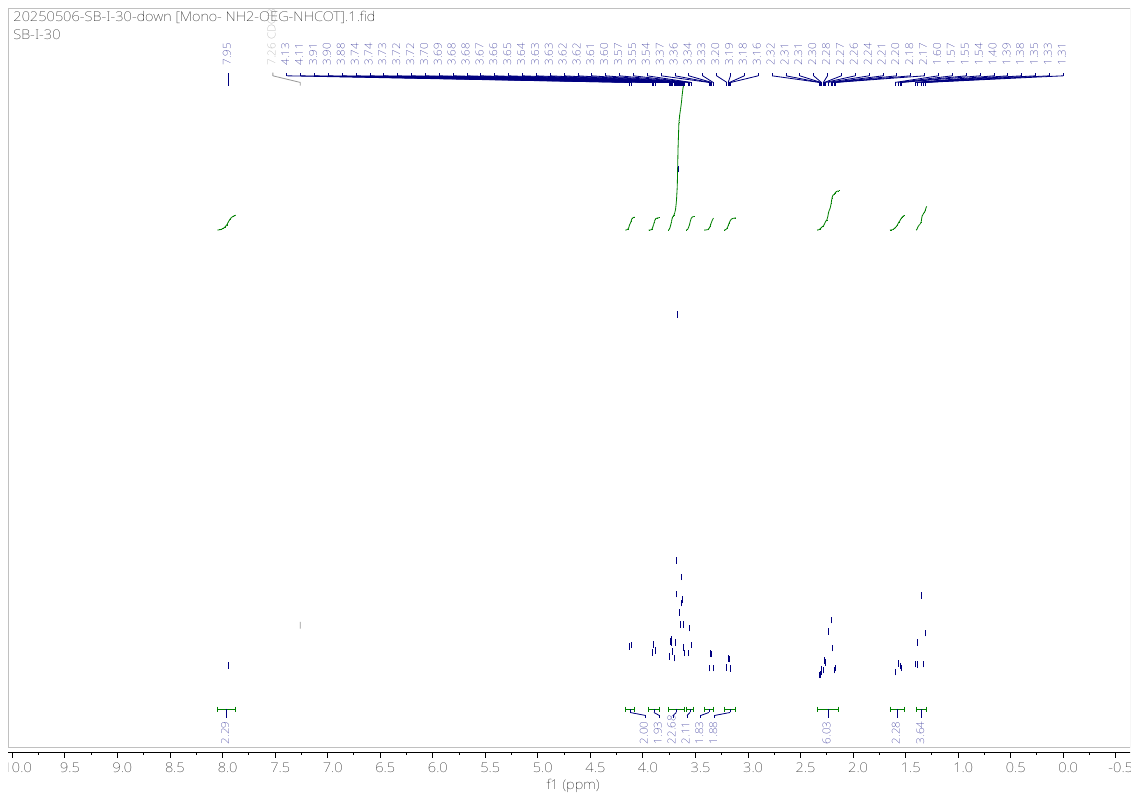

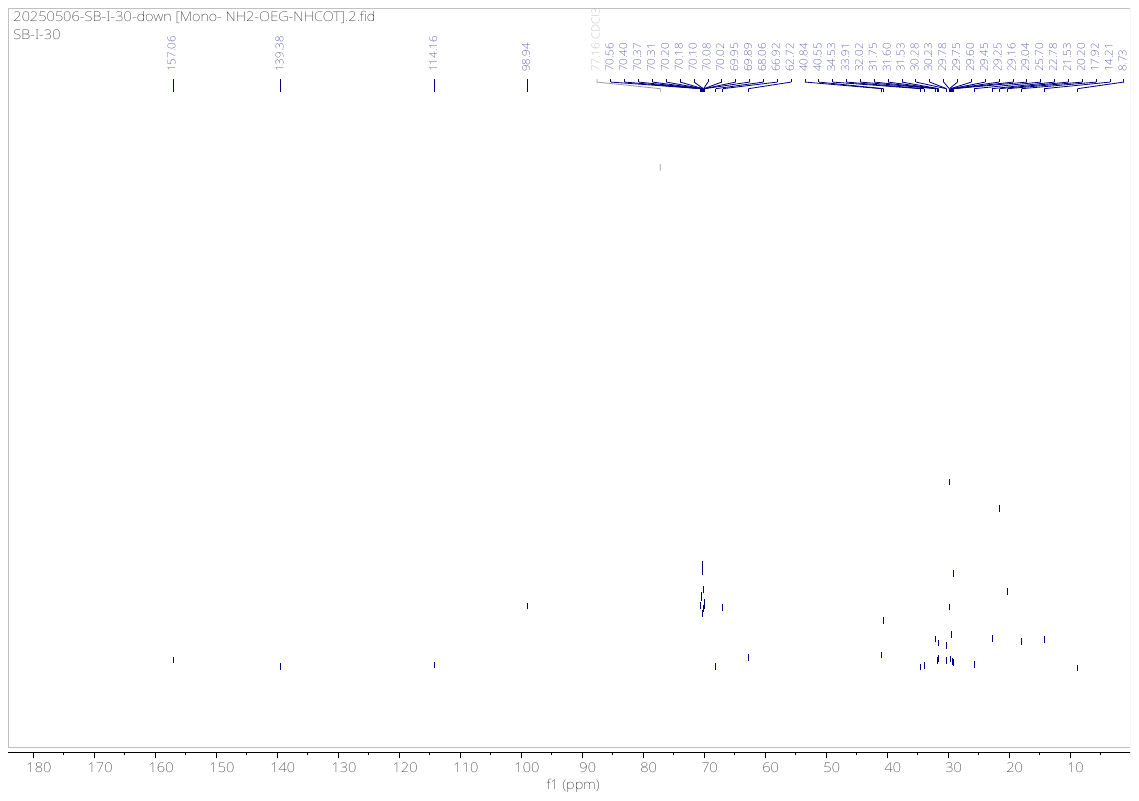

**Compound (1e)**

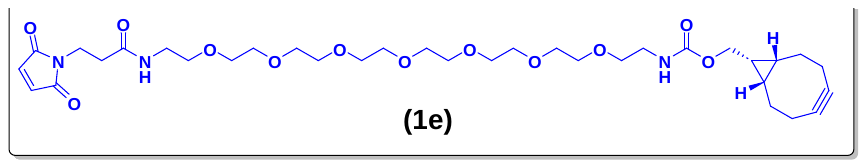

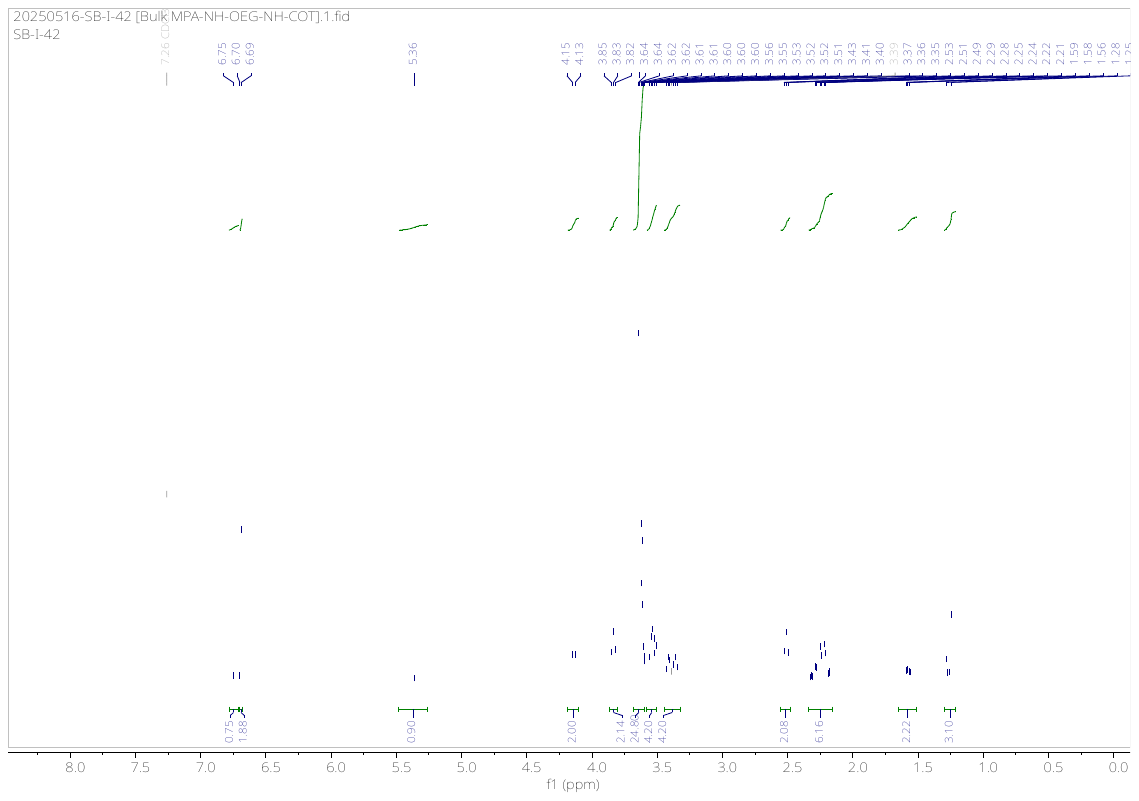

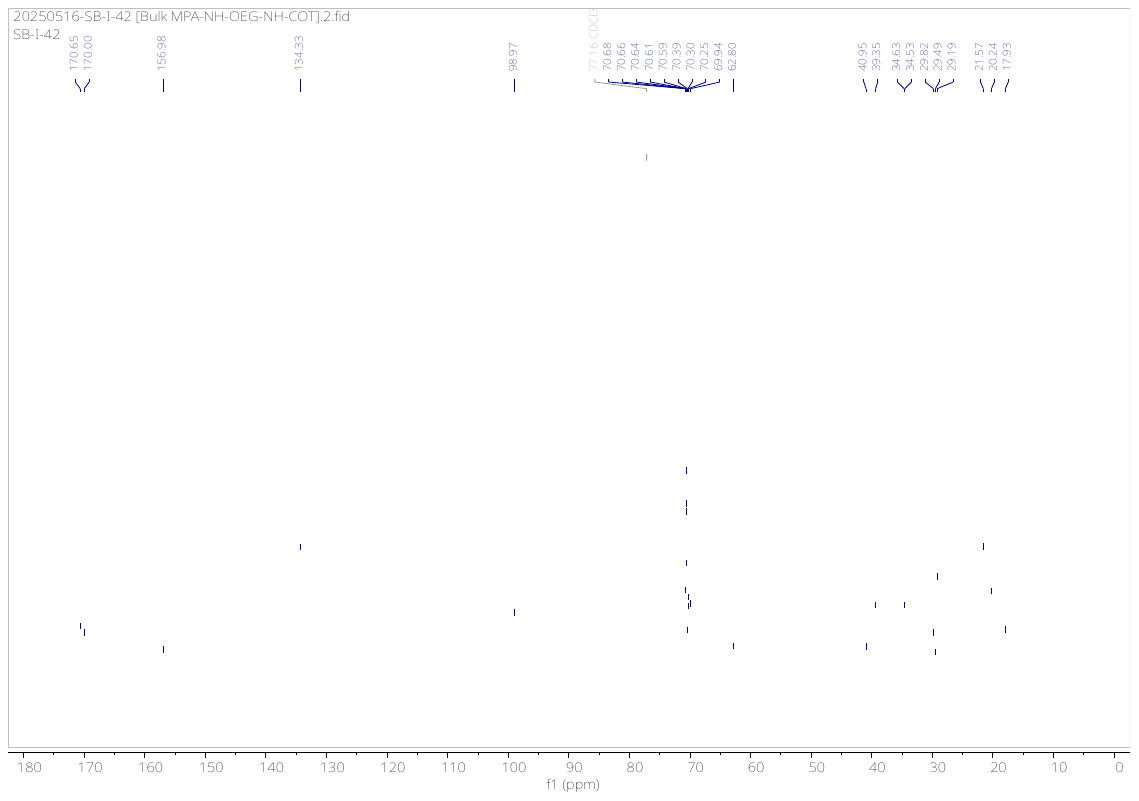

**Compound (2a)**

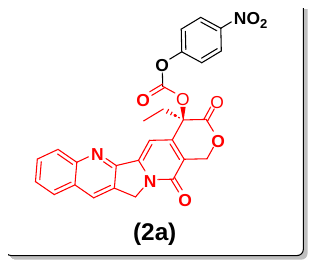

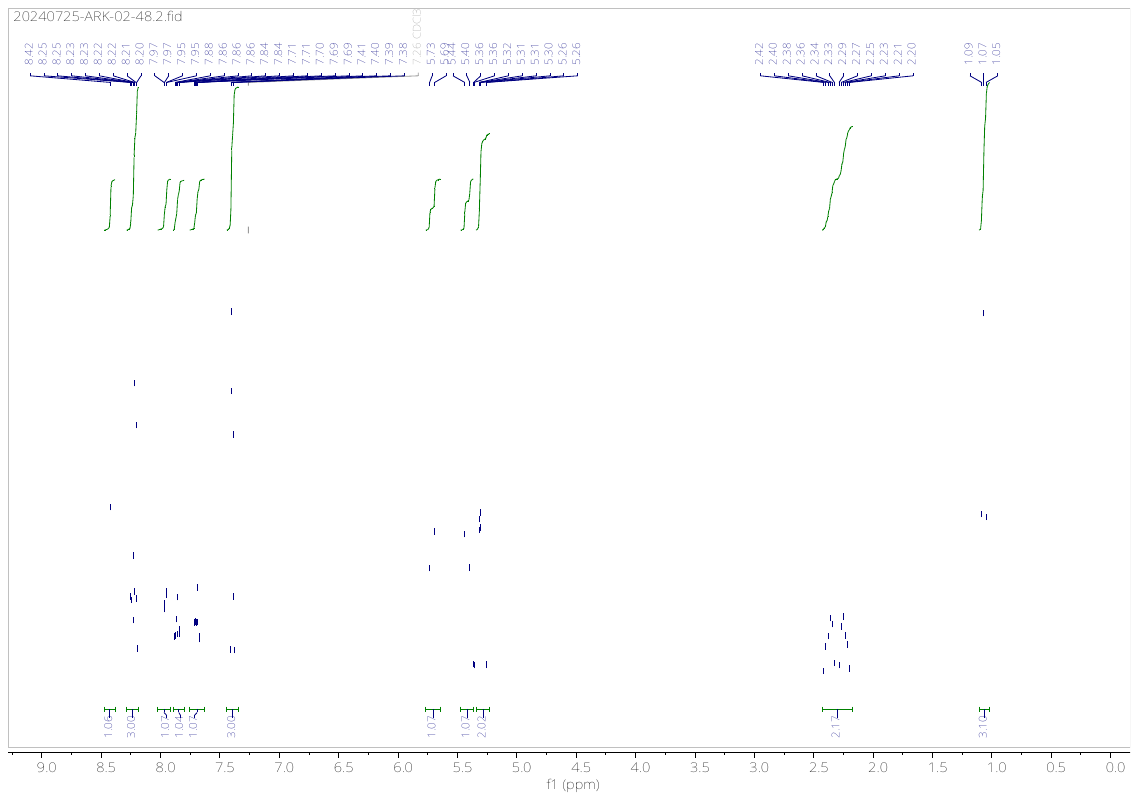

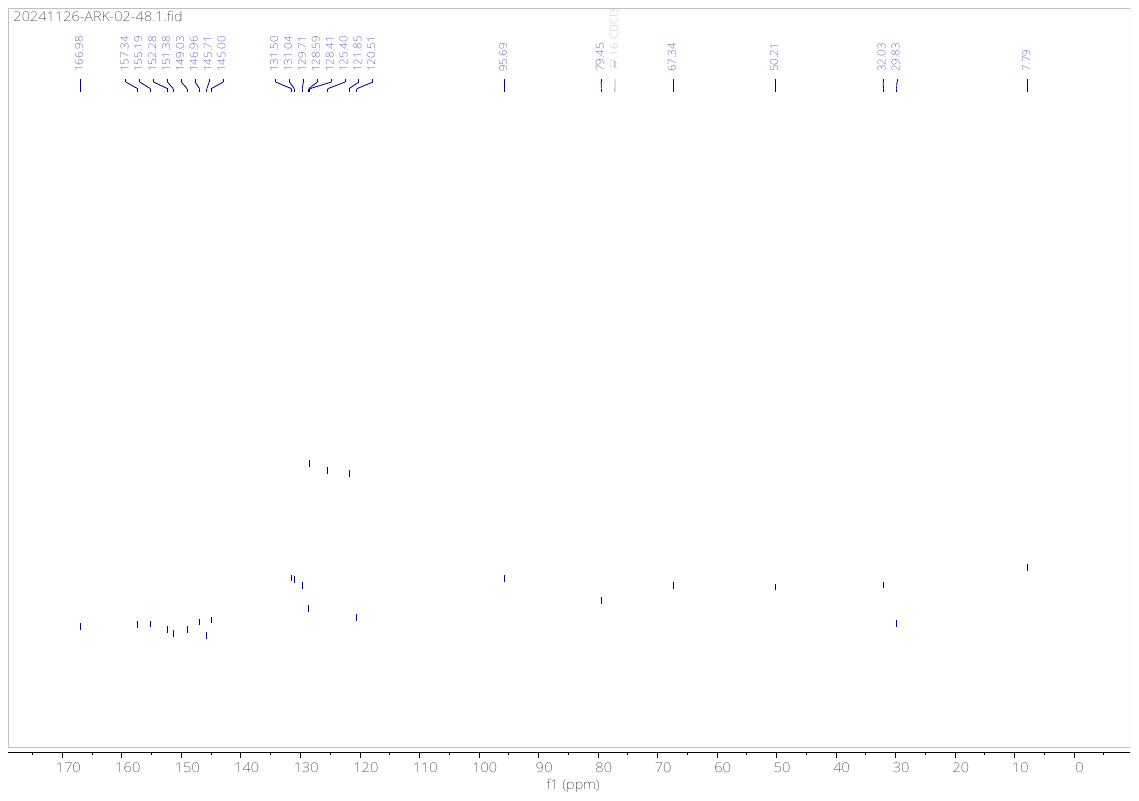

**Compound (3a)**

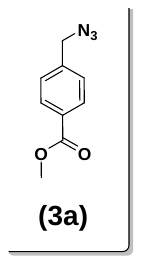

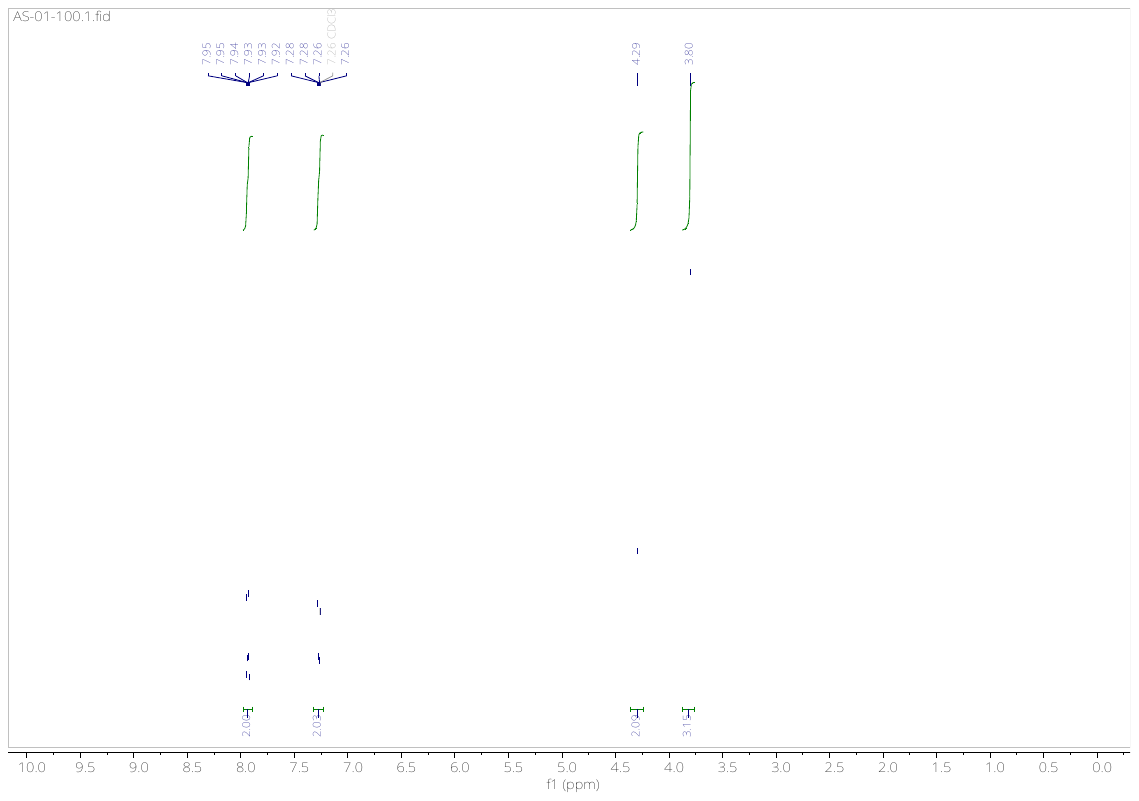

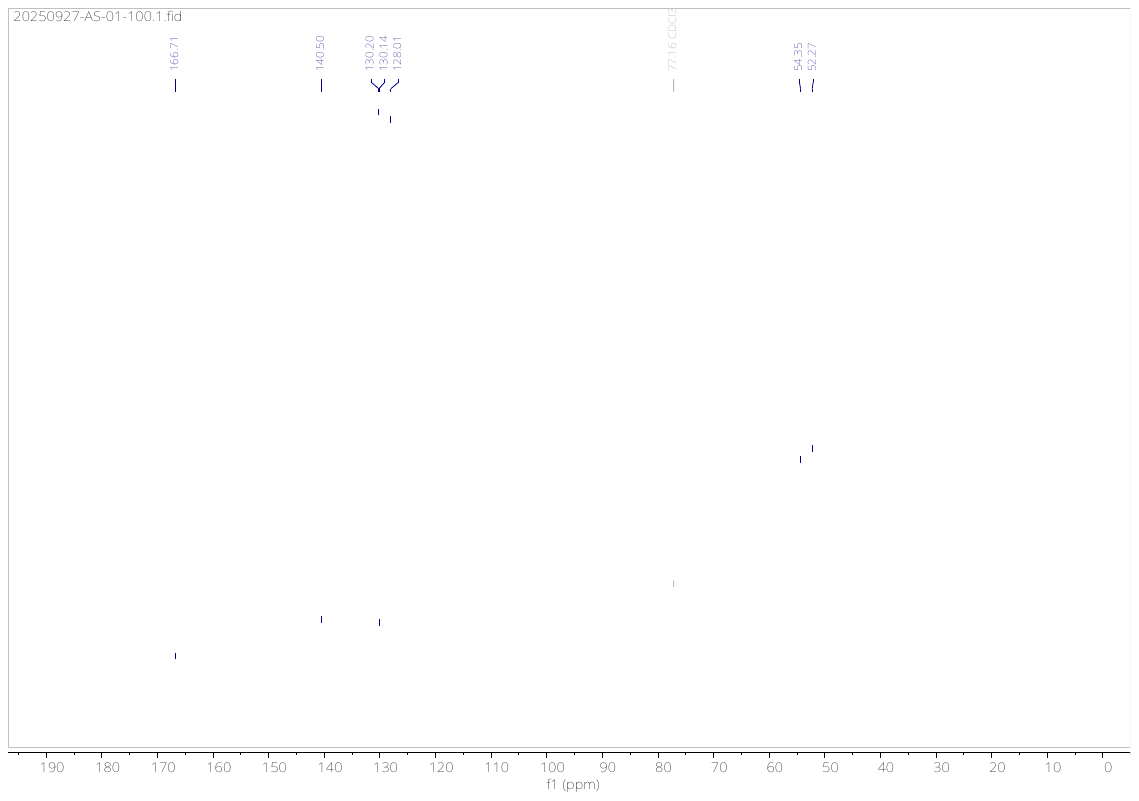

**Compound (3c)**

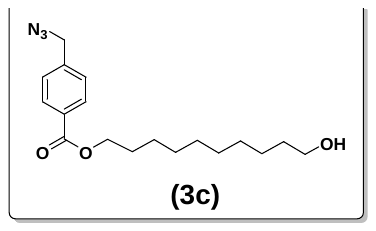

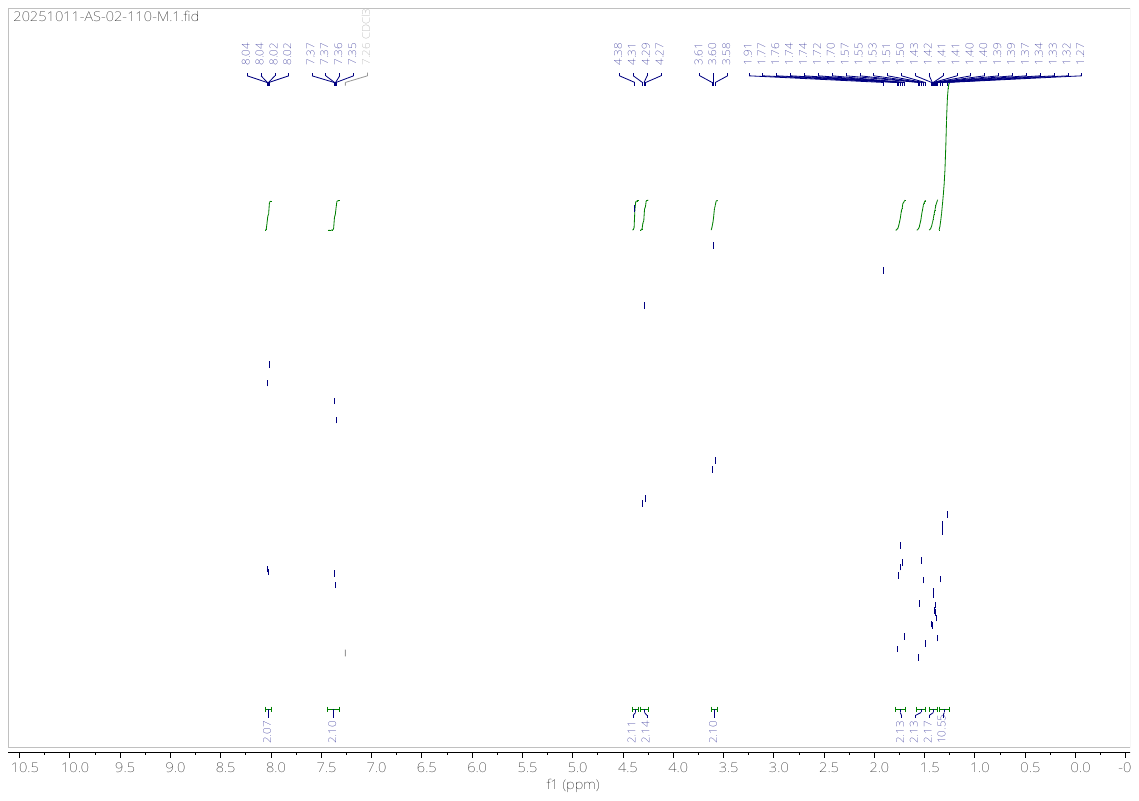

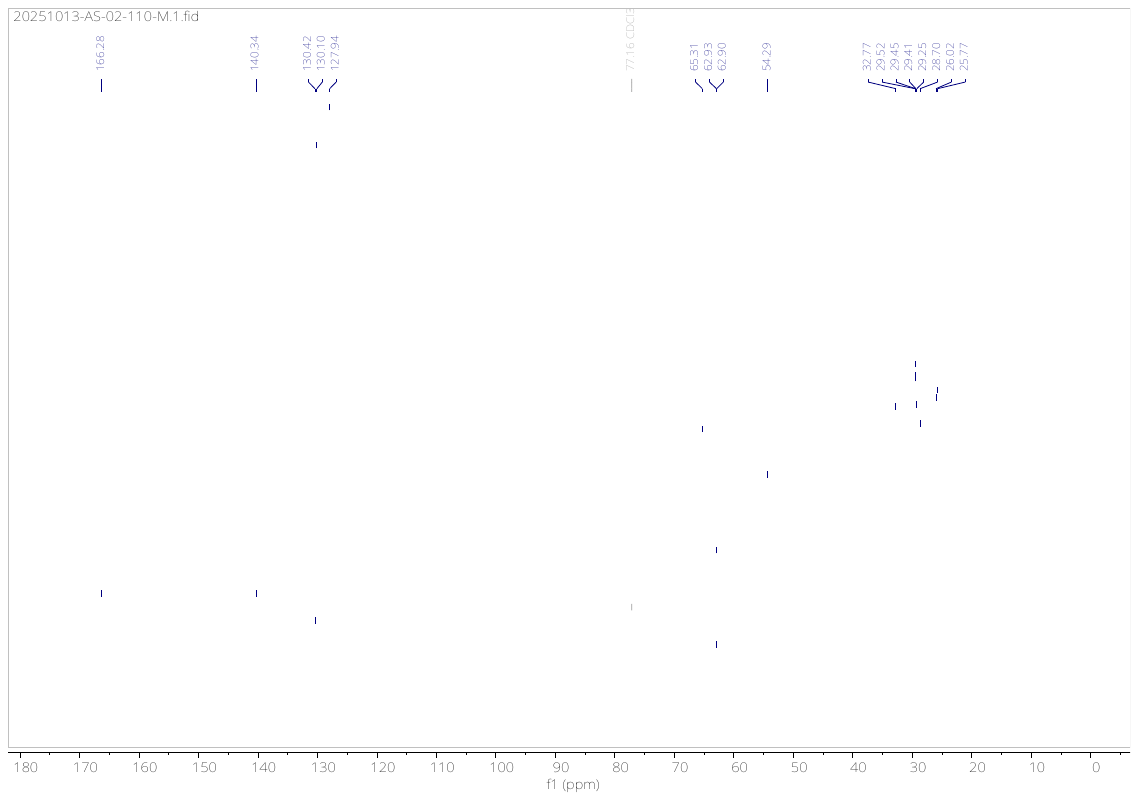

**Compound (3d)**

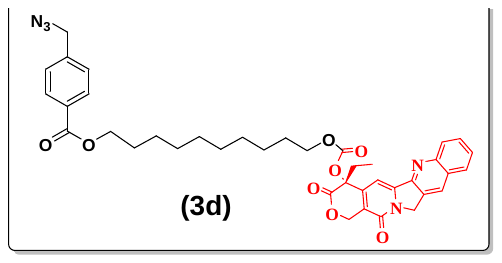

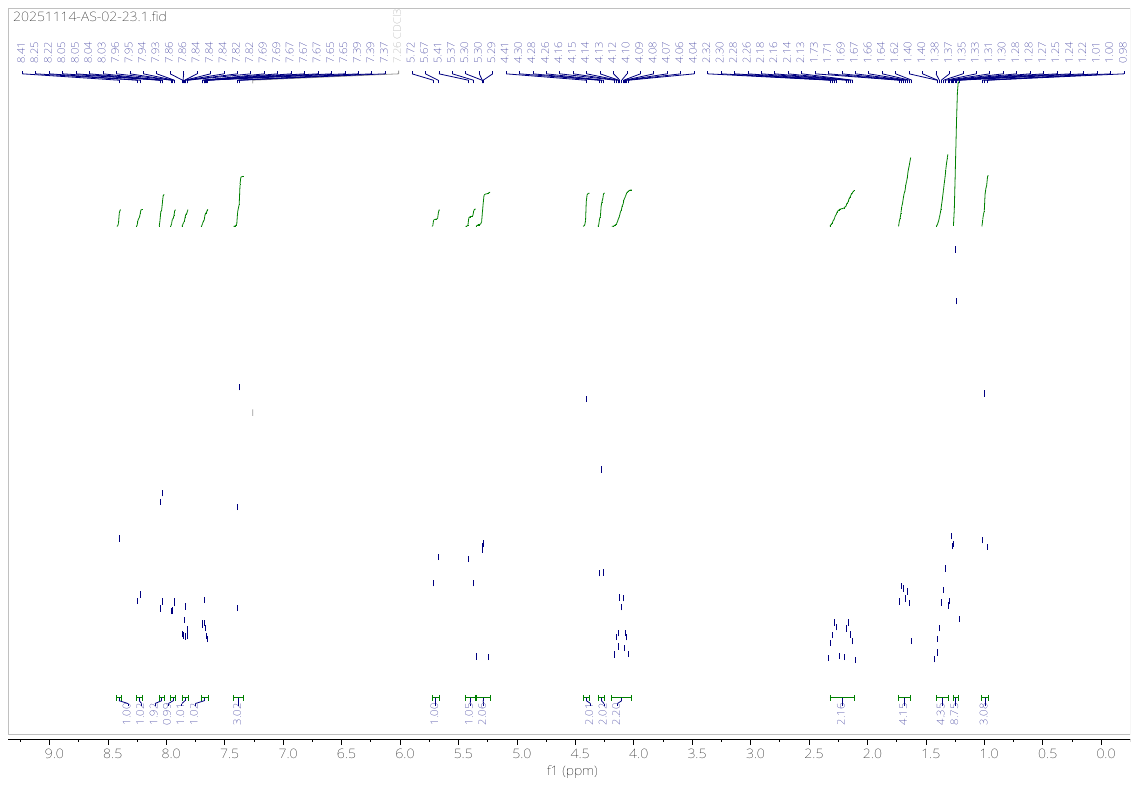

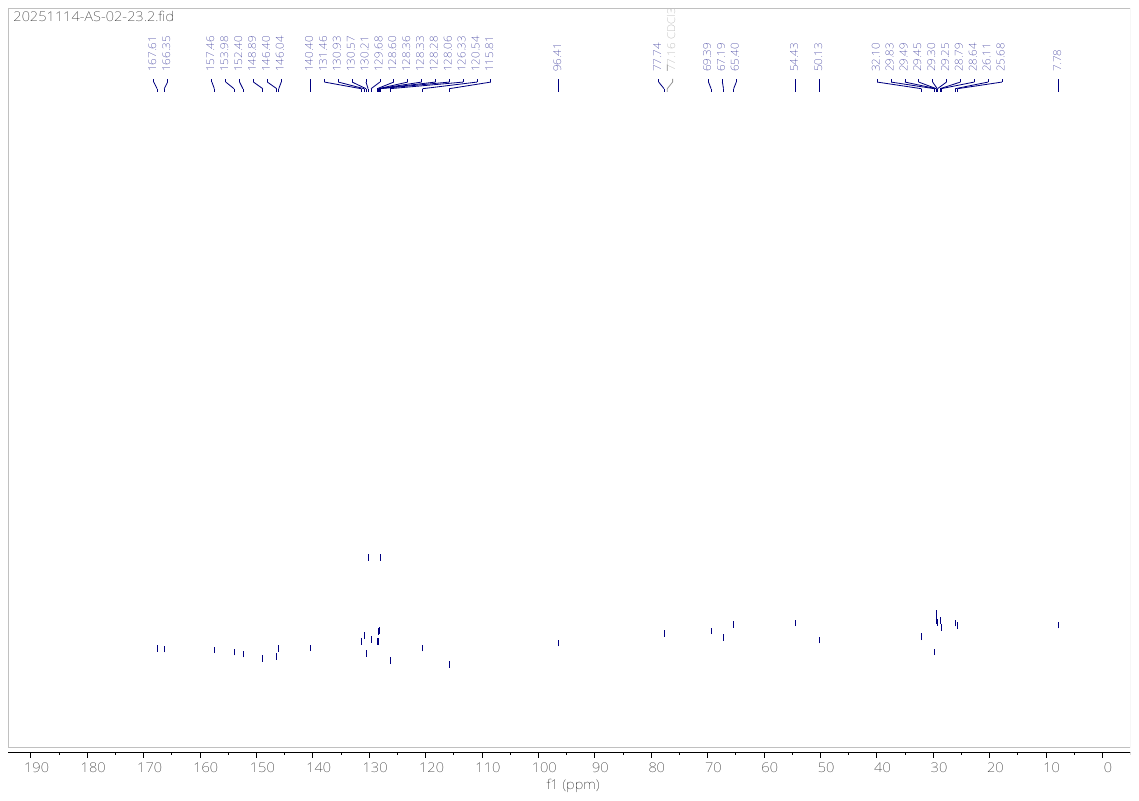

**Compound (4b)**

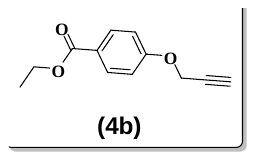

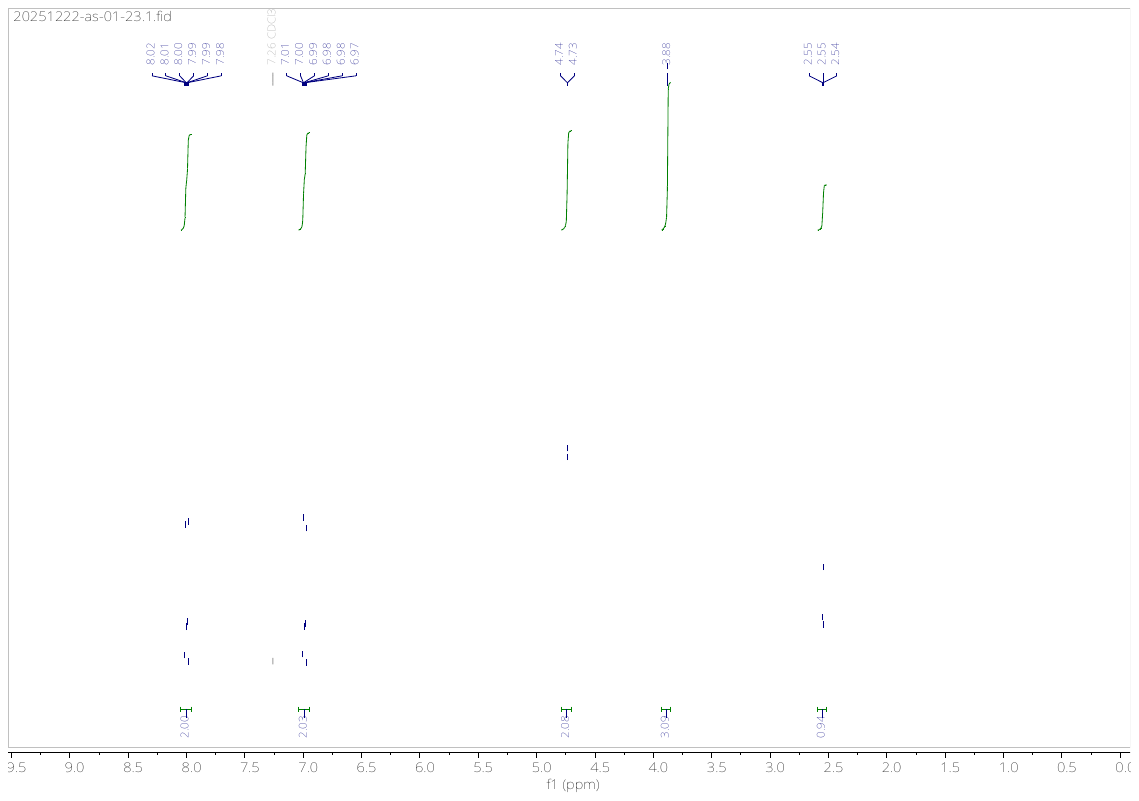

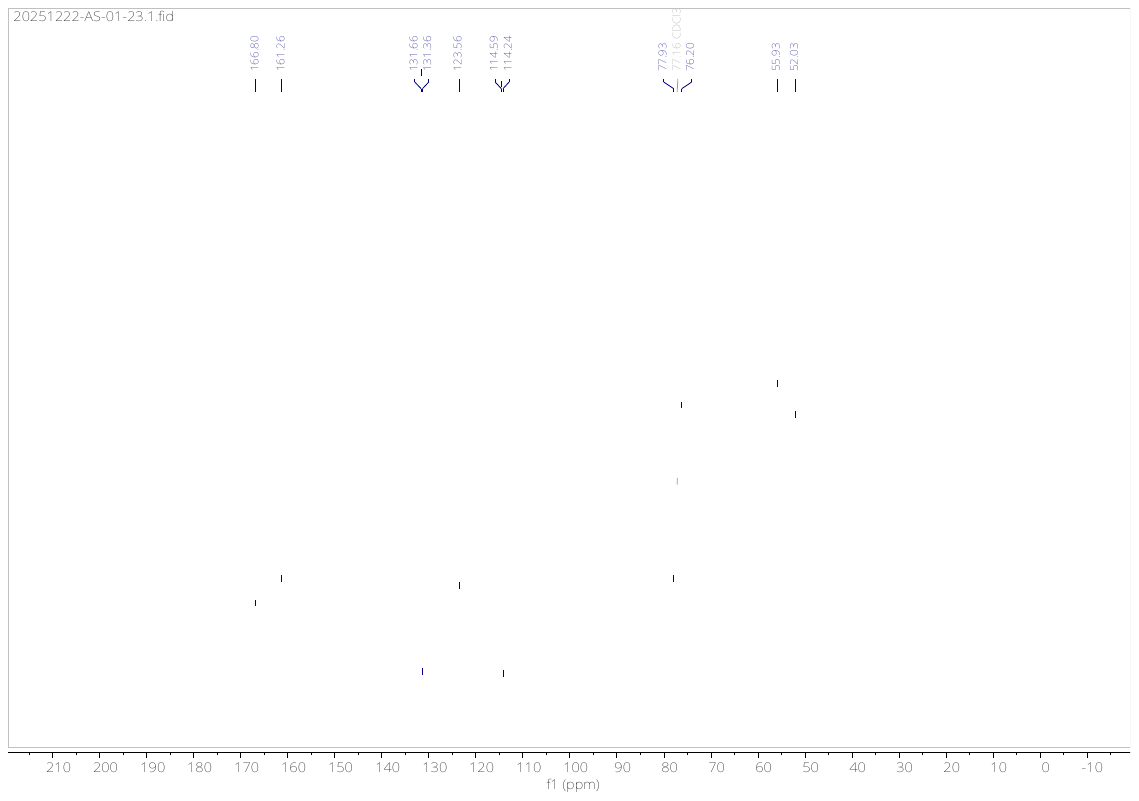

**Compound (4d)**

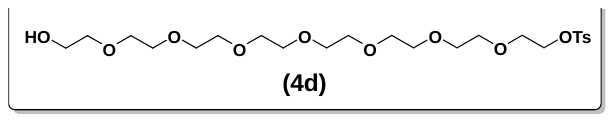

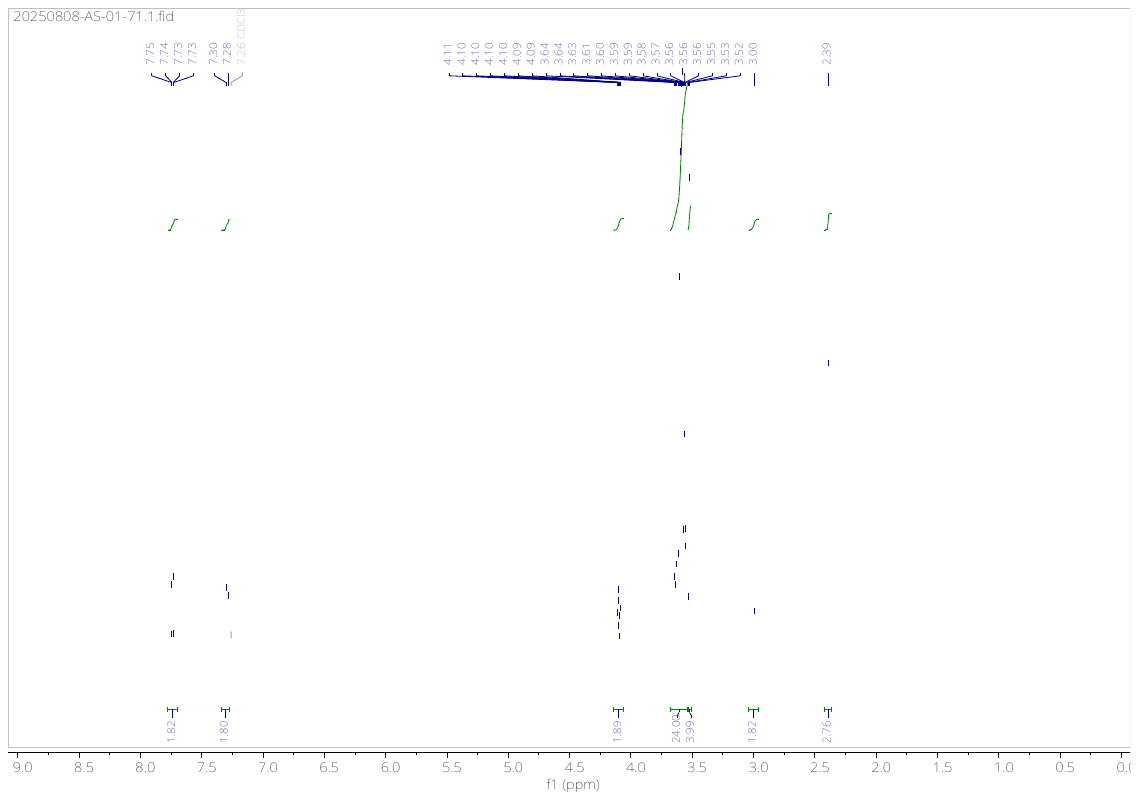

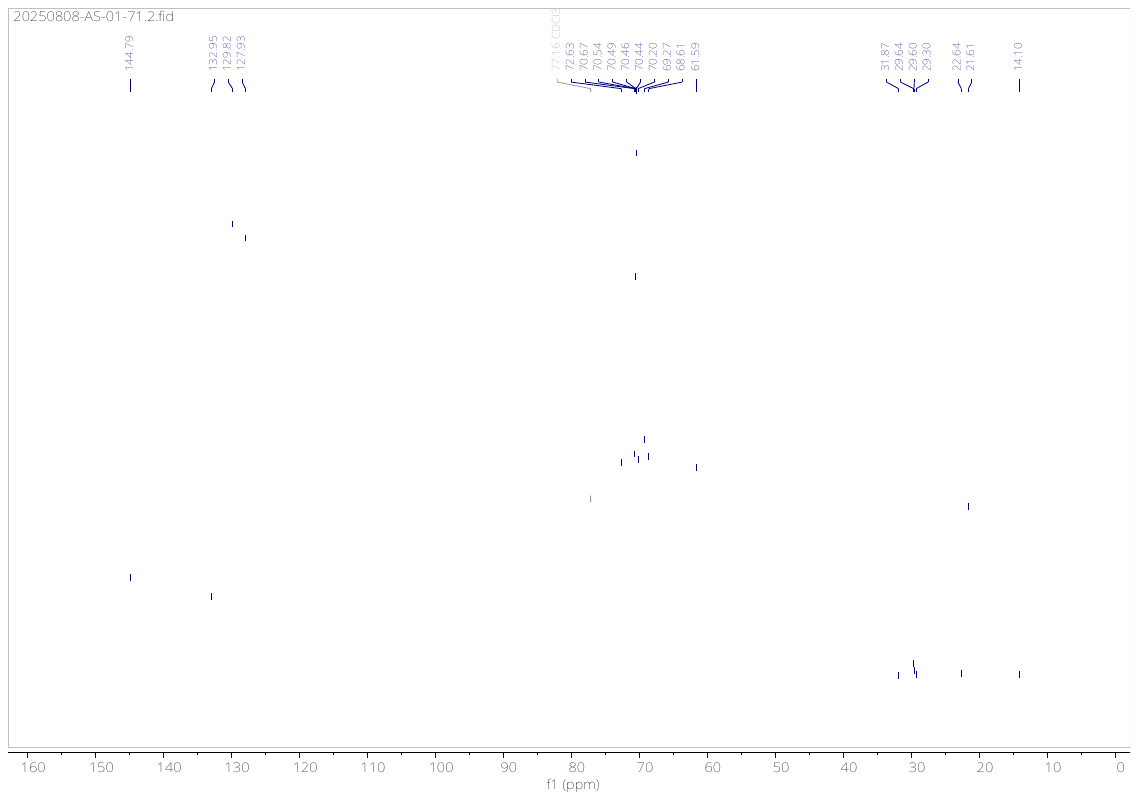

**Compound (4e)**

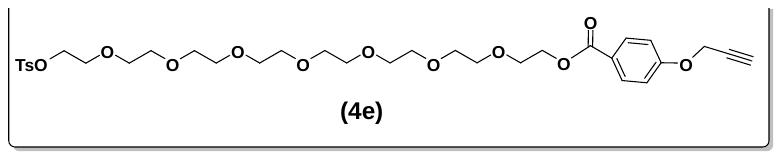

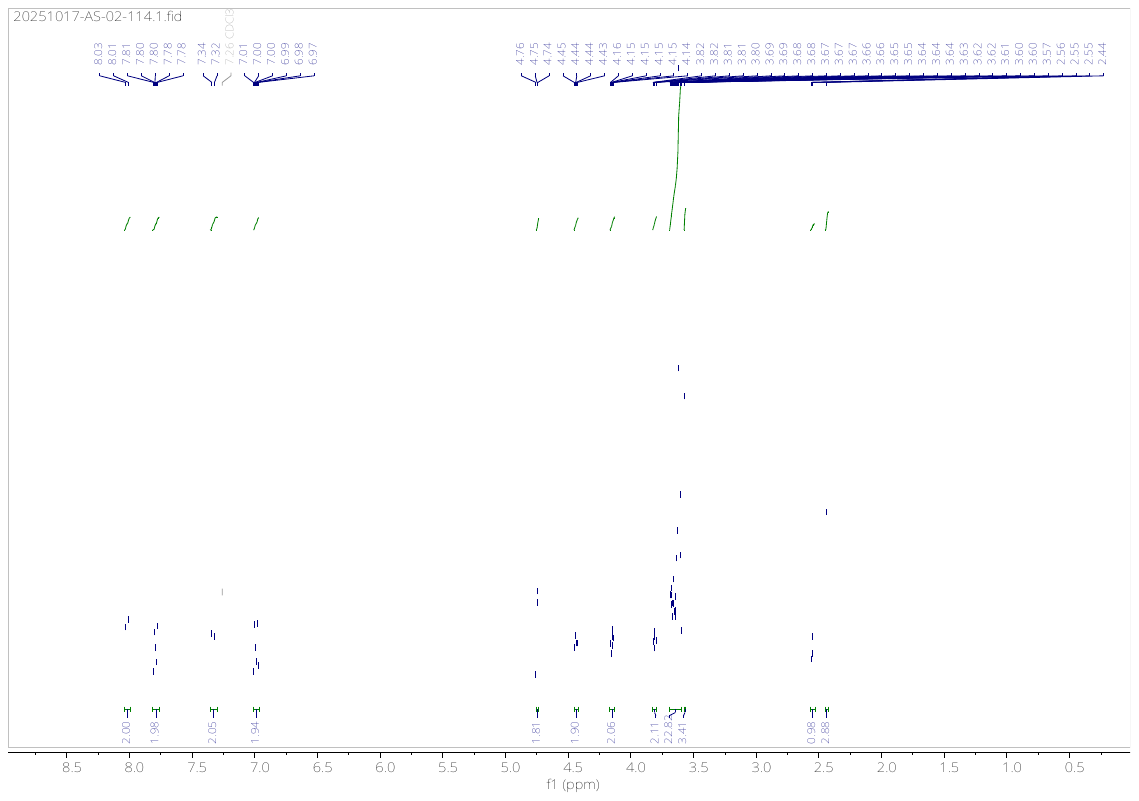

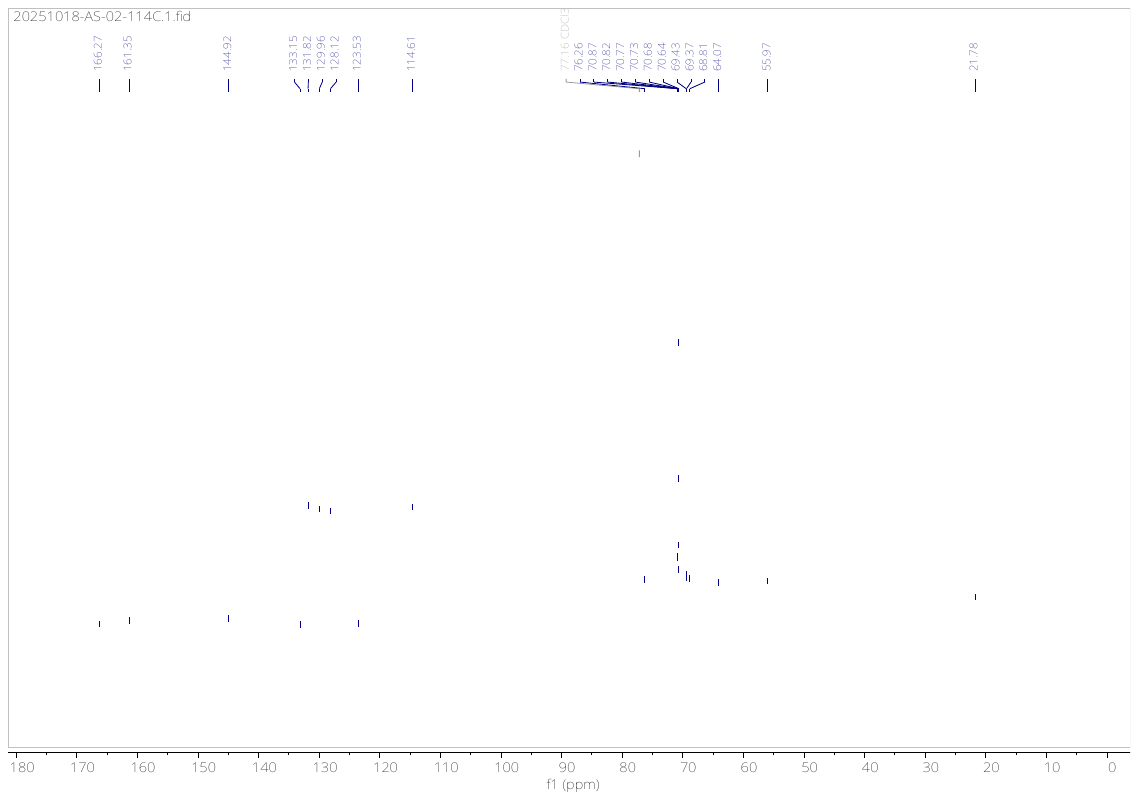

**Compound (4f)**

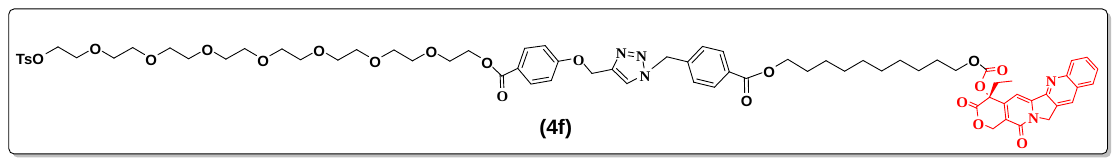

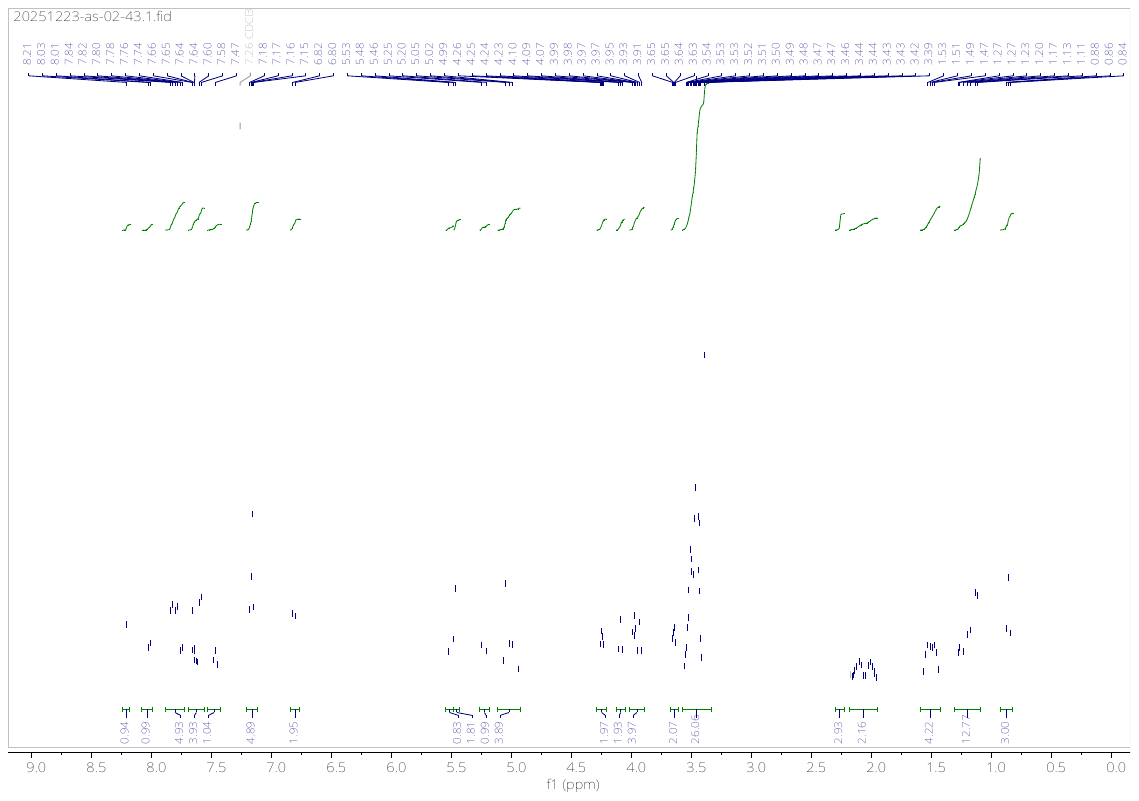

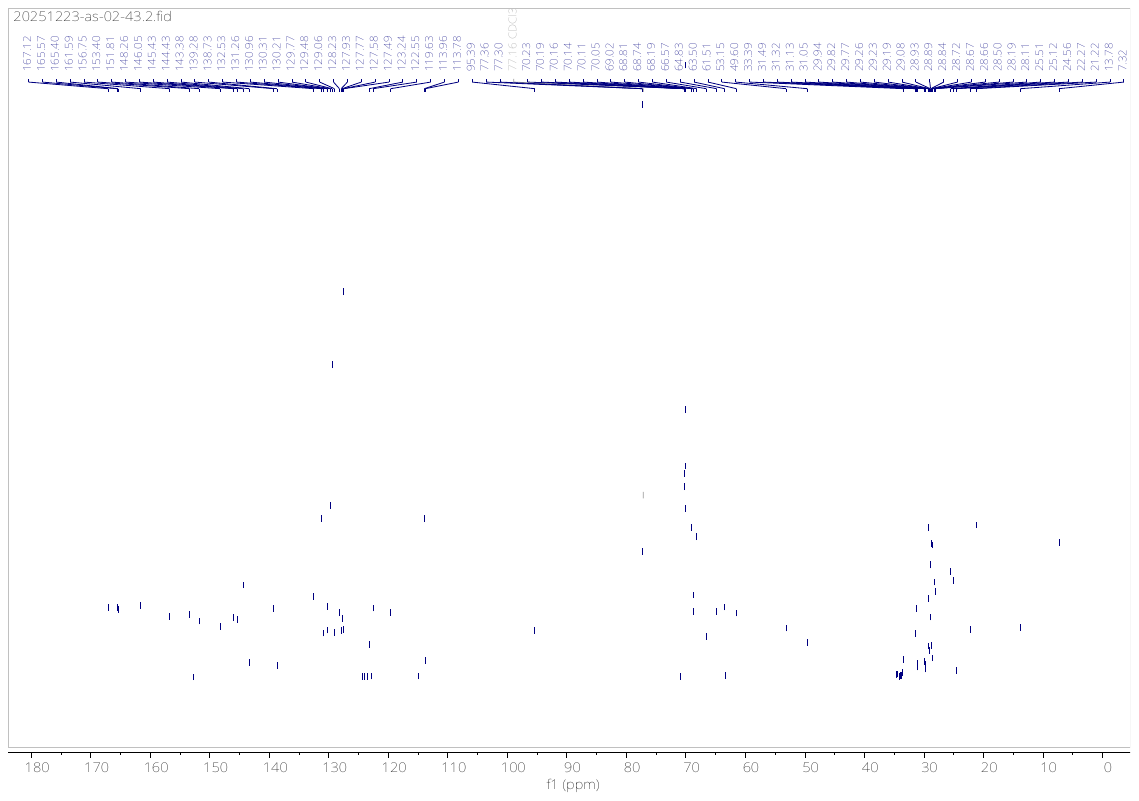

**Compound (4g)**

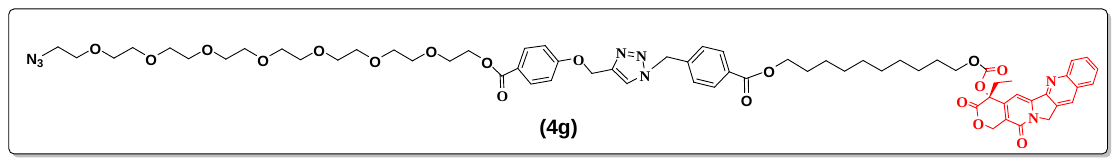

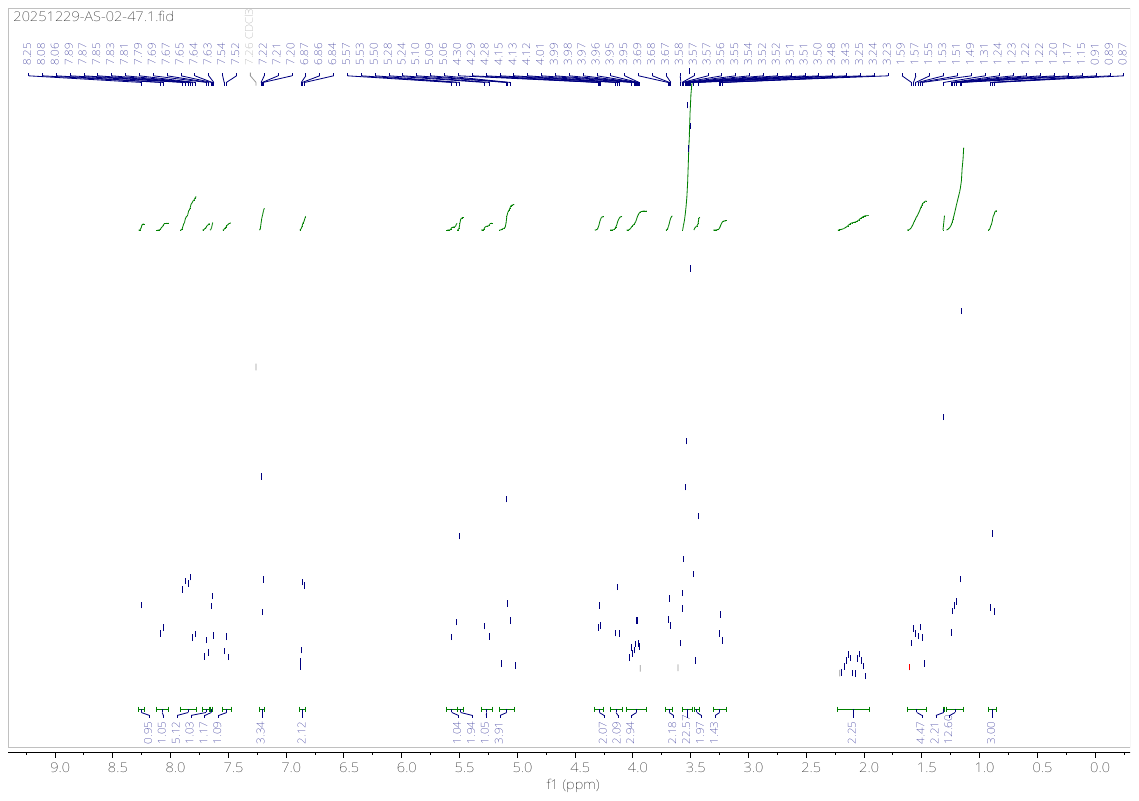

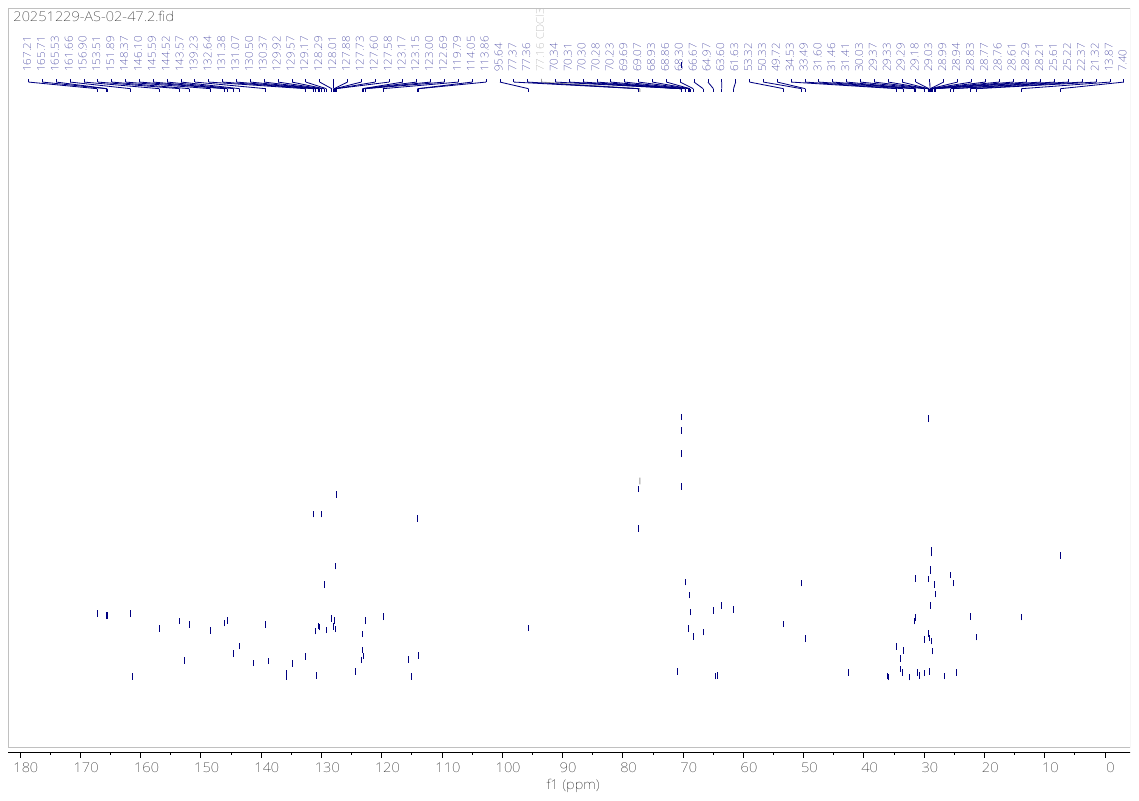

**Compound (5b)**

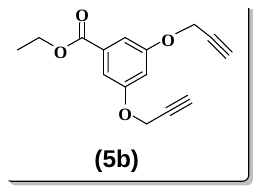

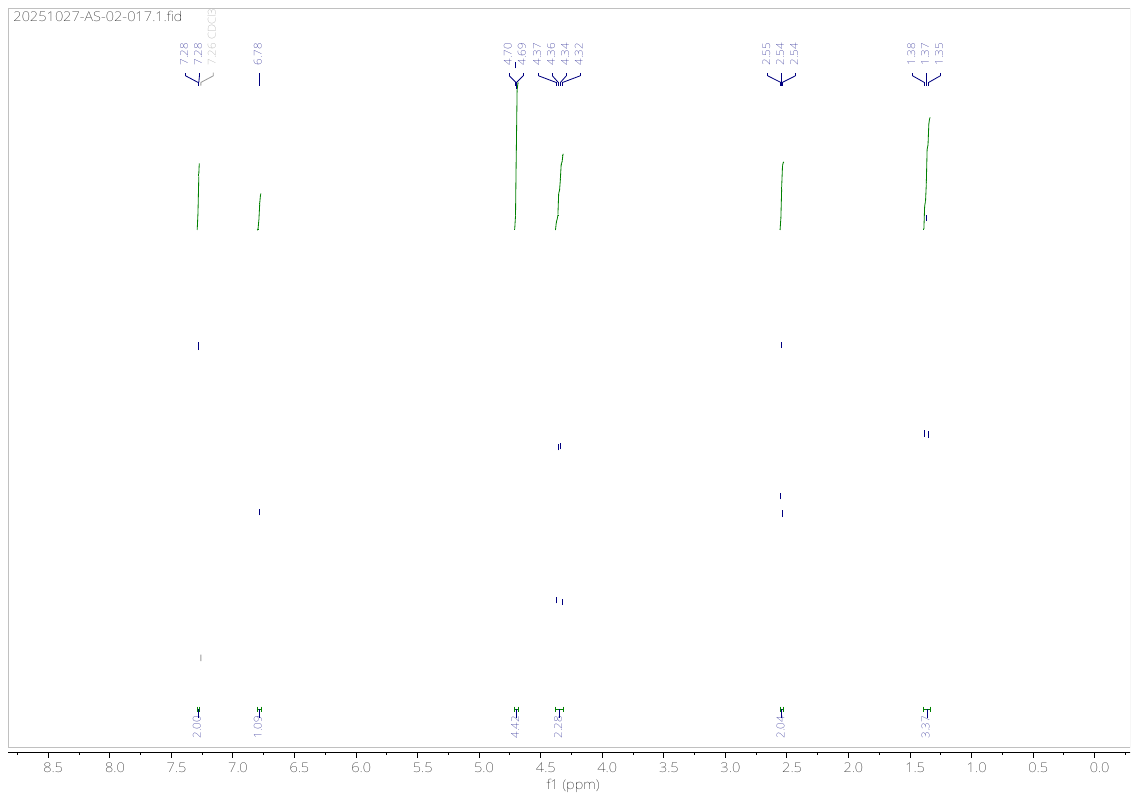

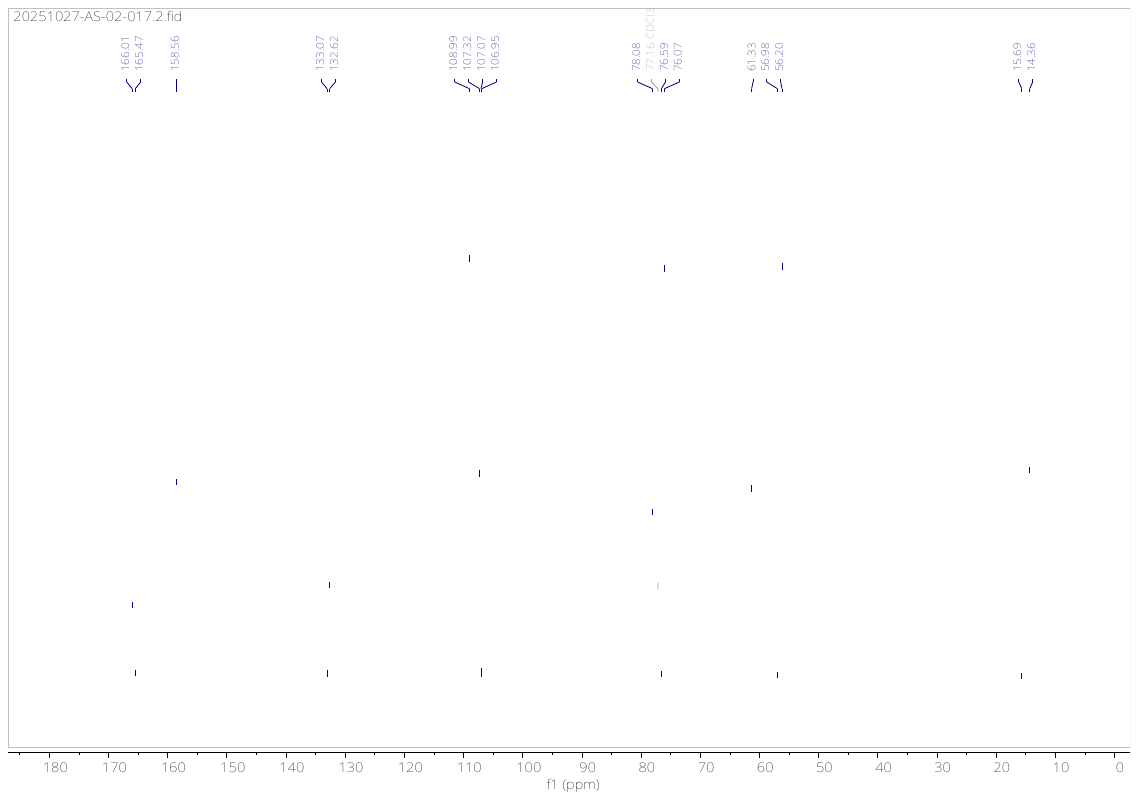

**Compound (5d)**

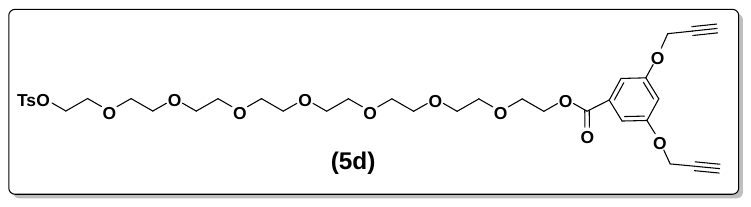

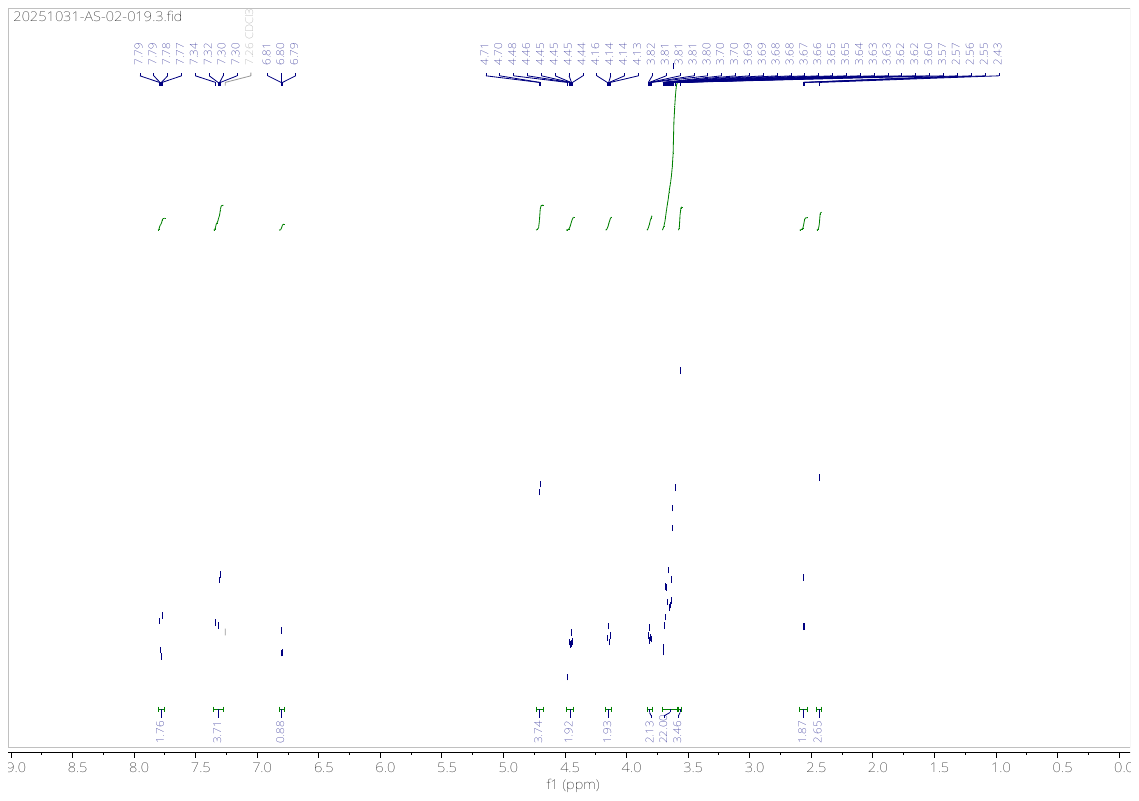

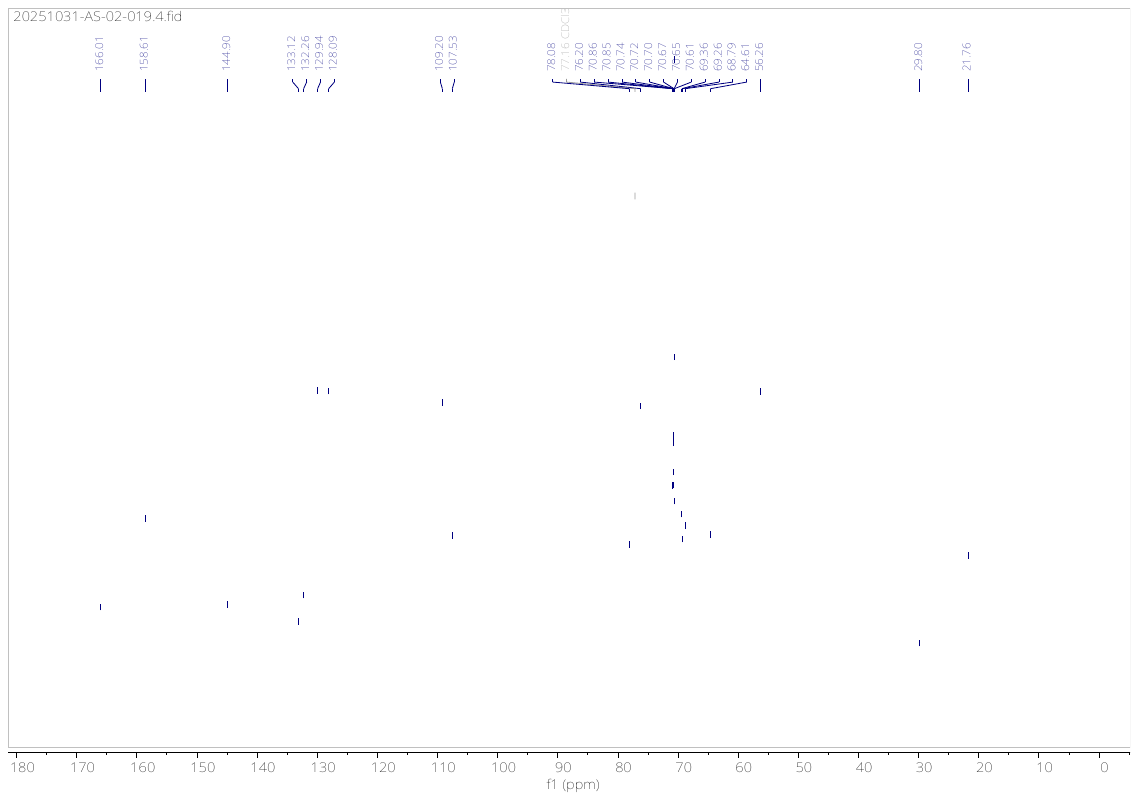

**Compound (5e)**

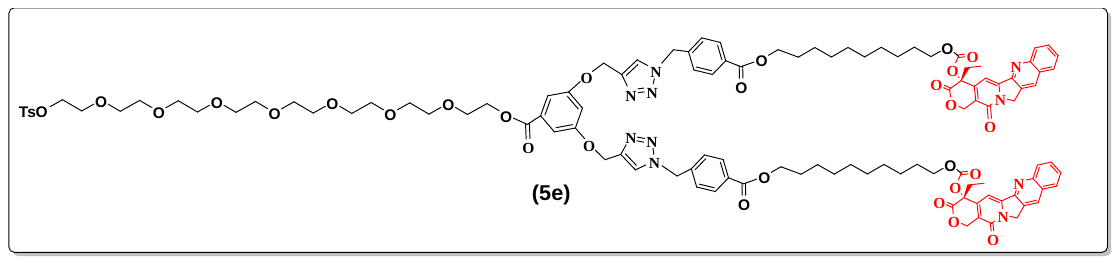

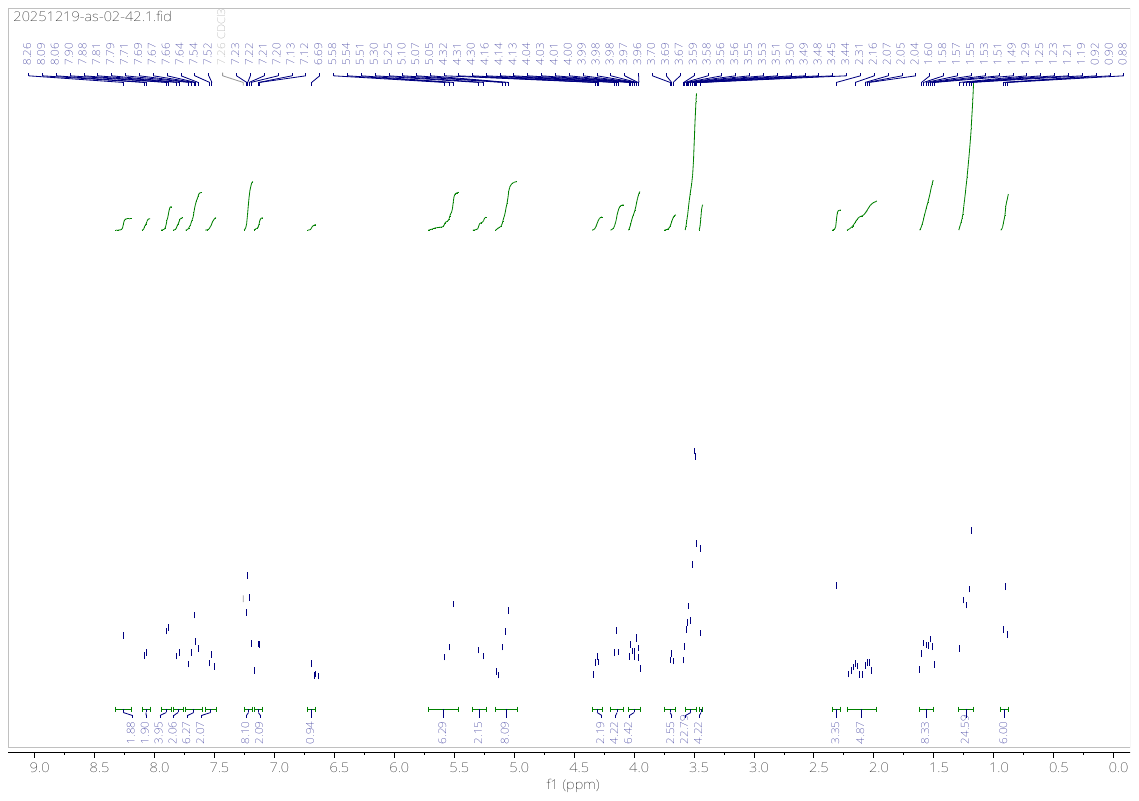

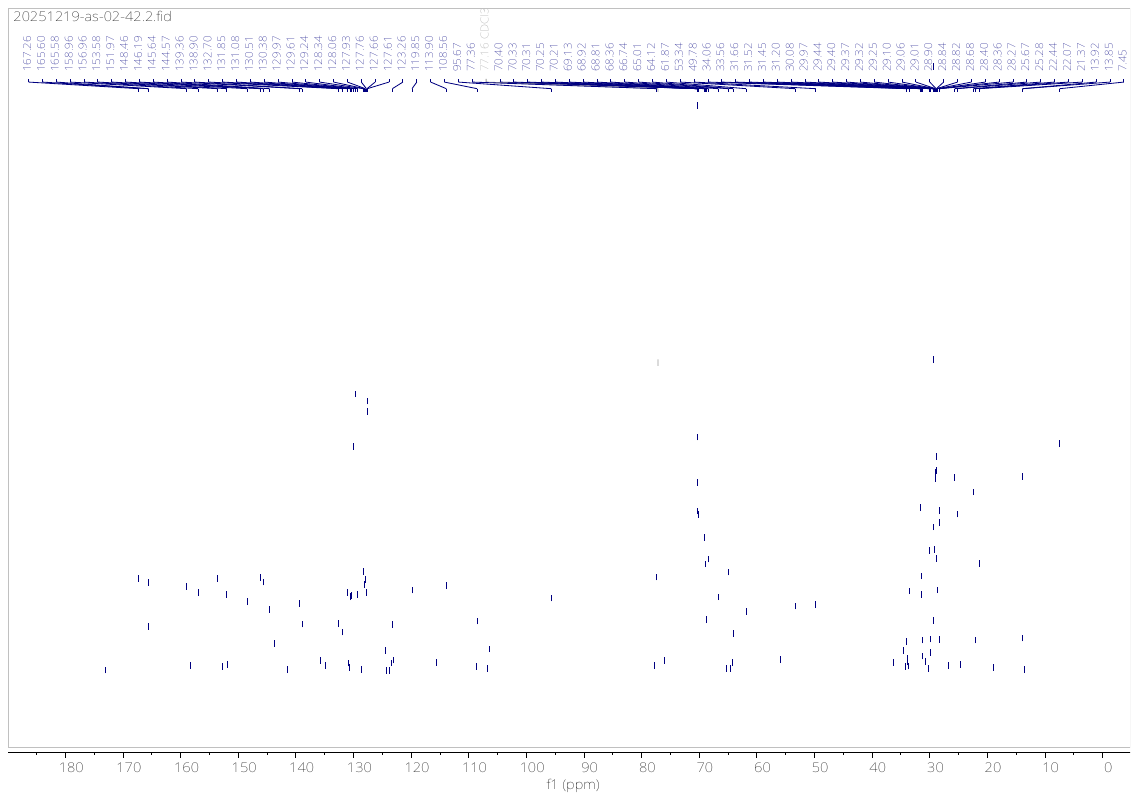

**Compound (5f)**

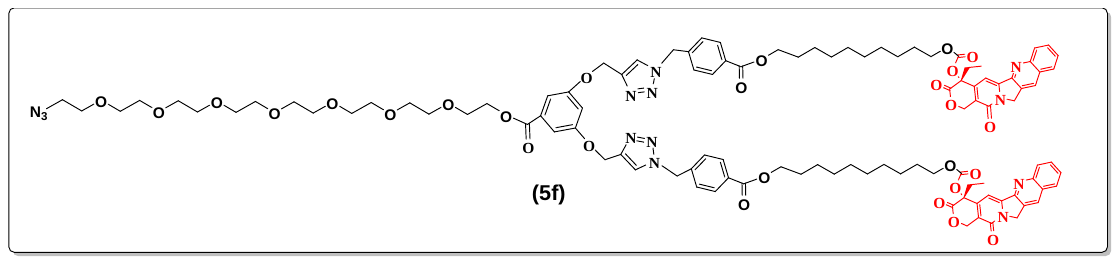

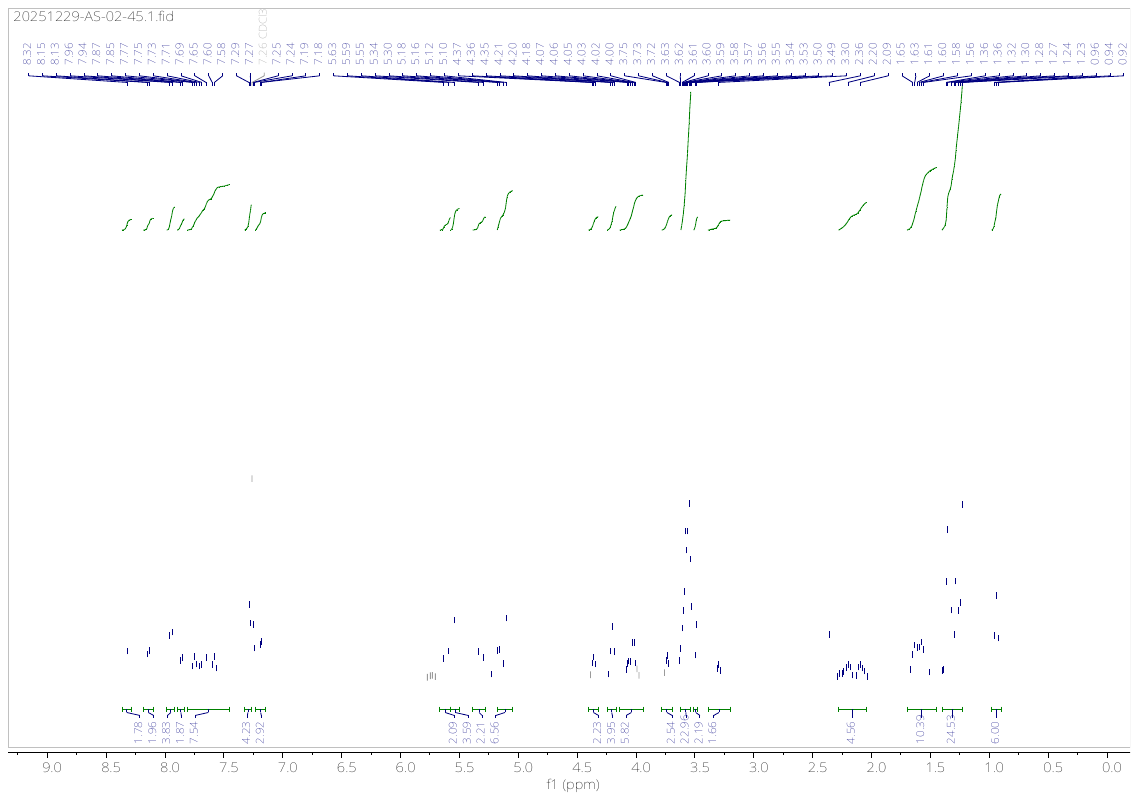

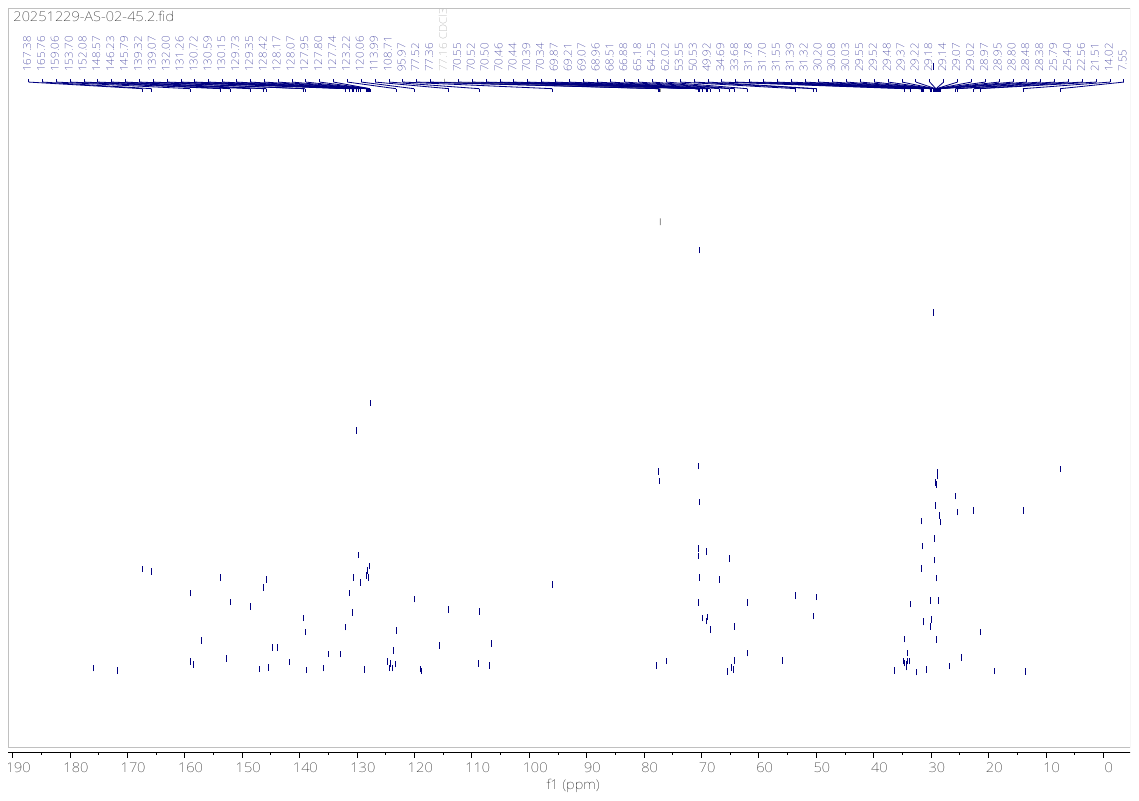

**Compound (6b)**

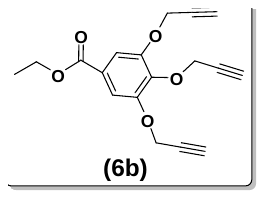

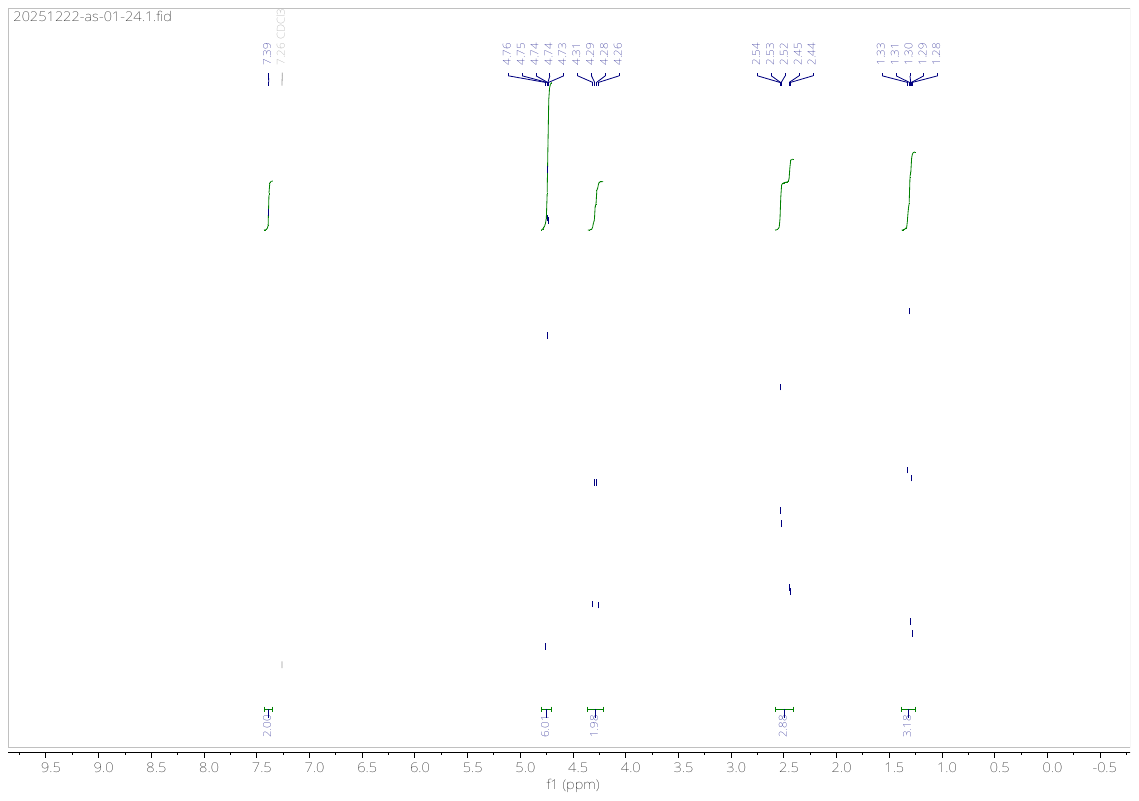

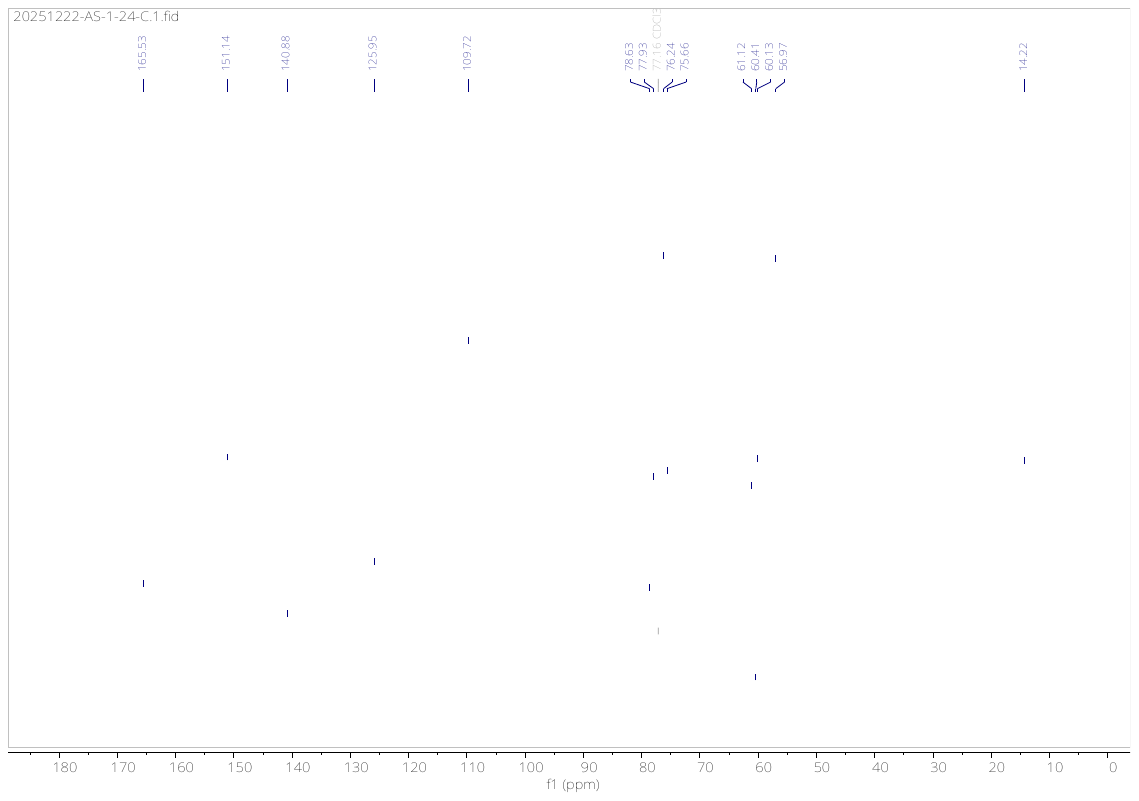

**Compound (6d)**

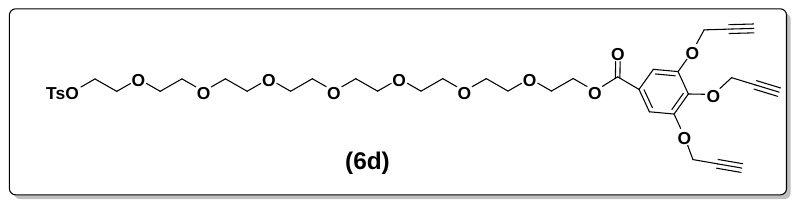

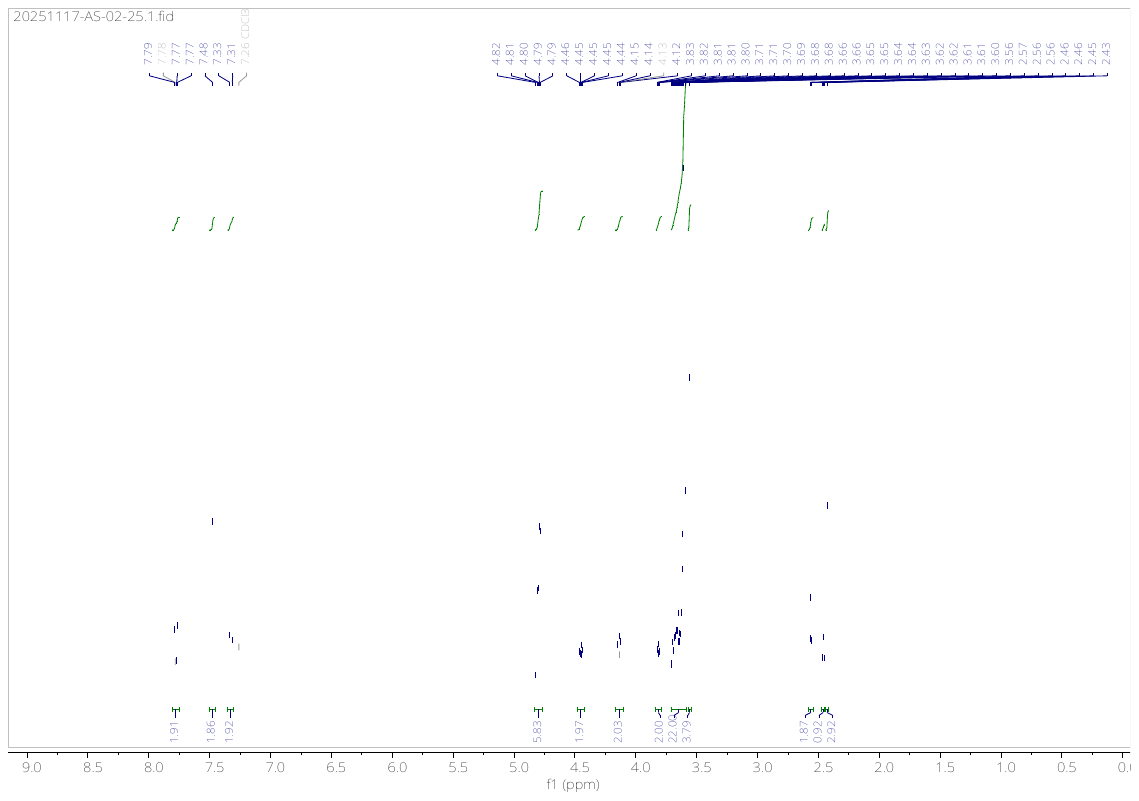

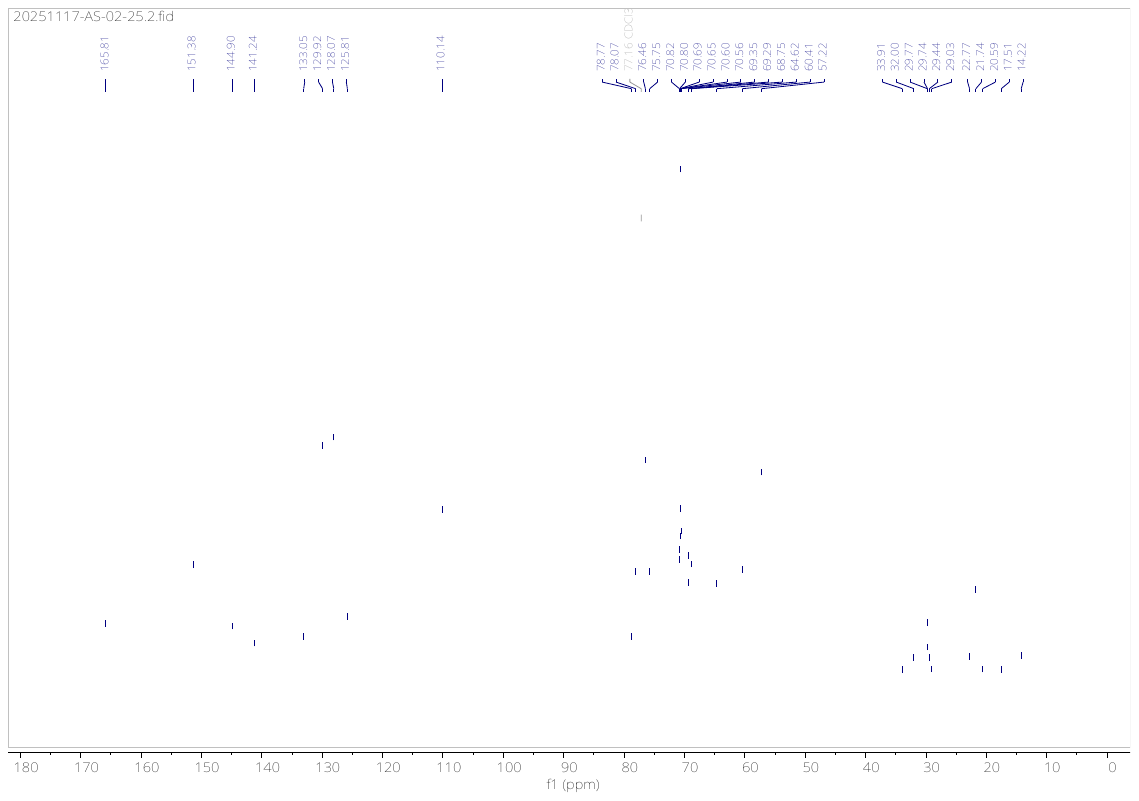

**Compound (6e)**

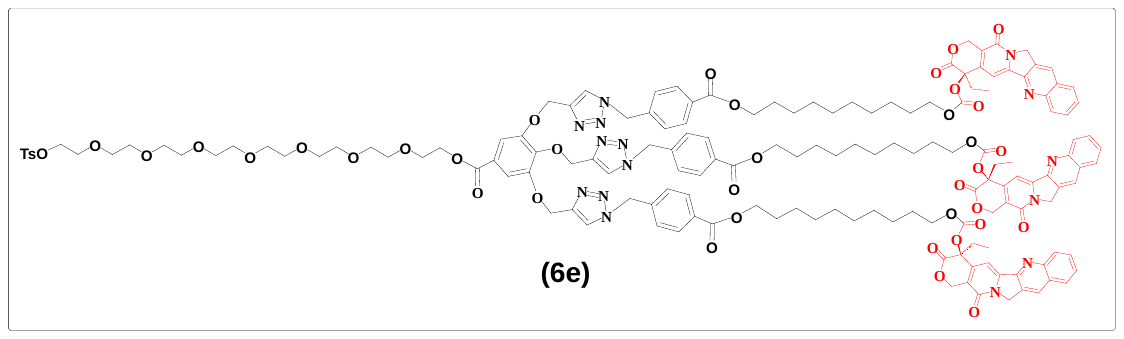

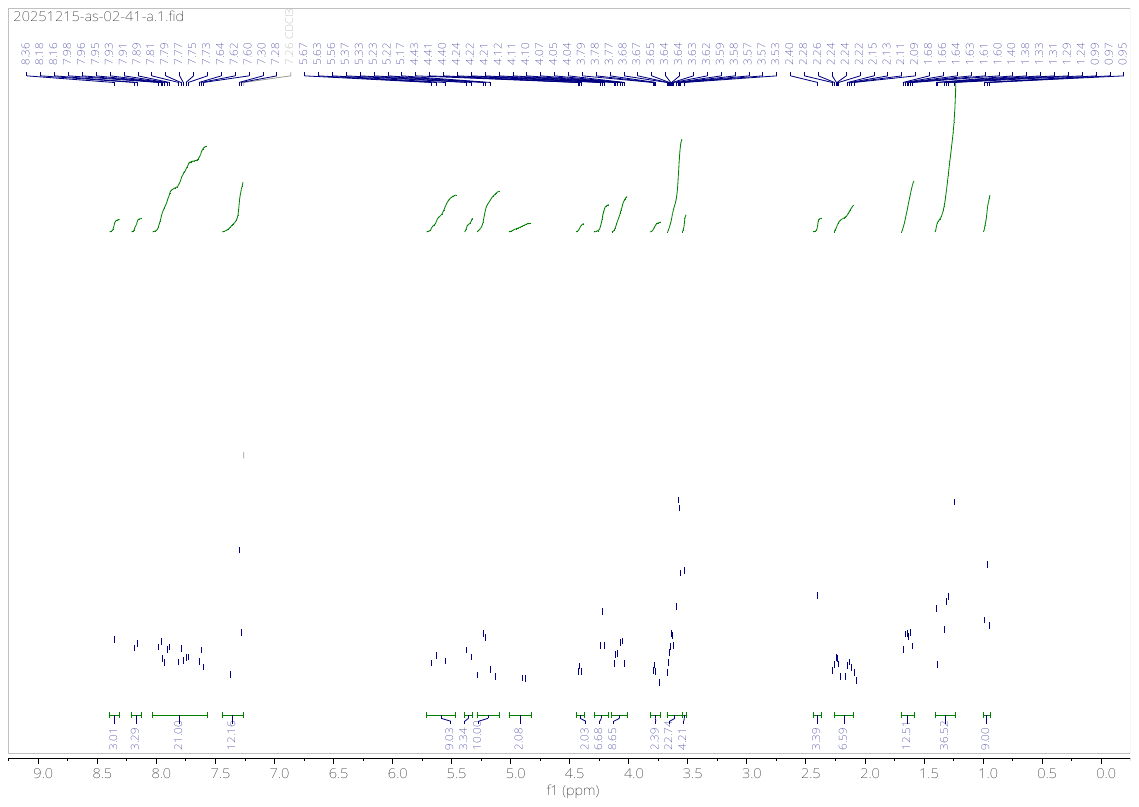

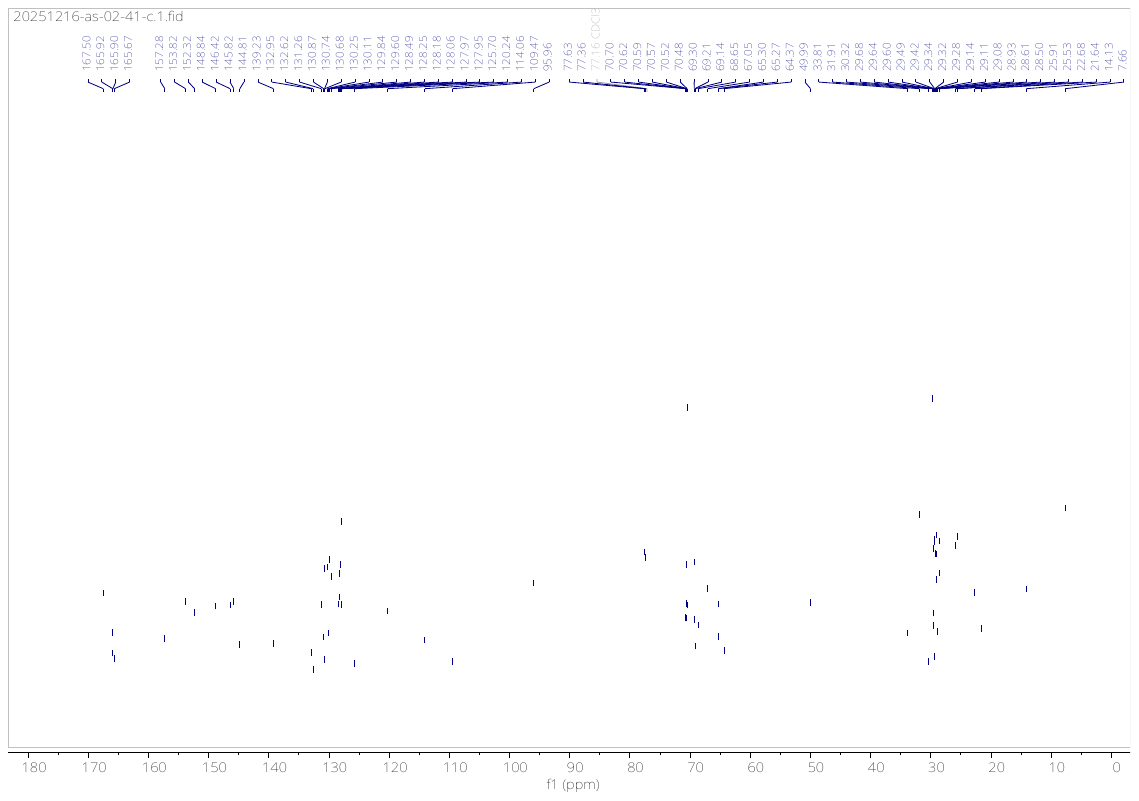

**Compound (6f)**

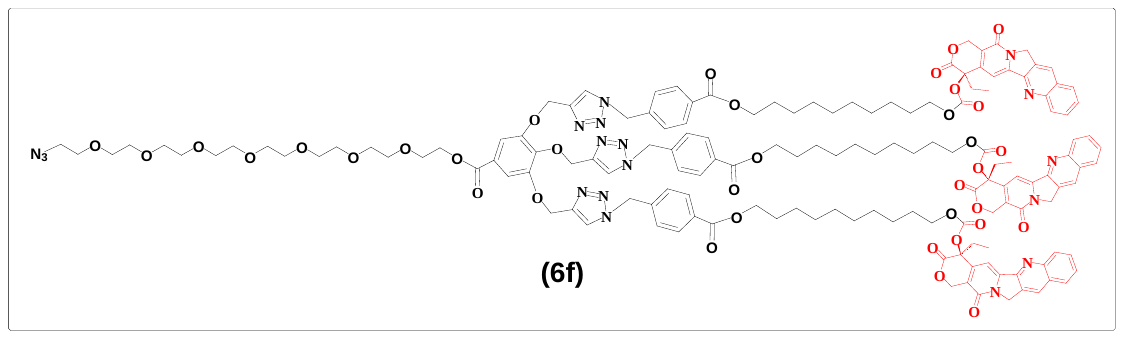

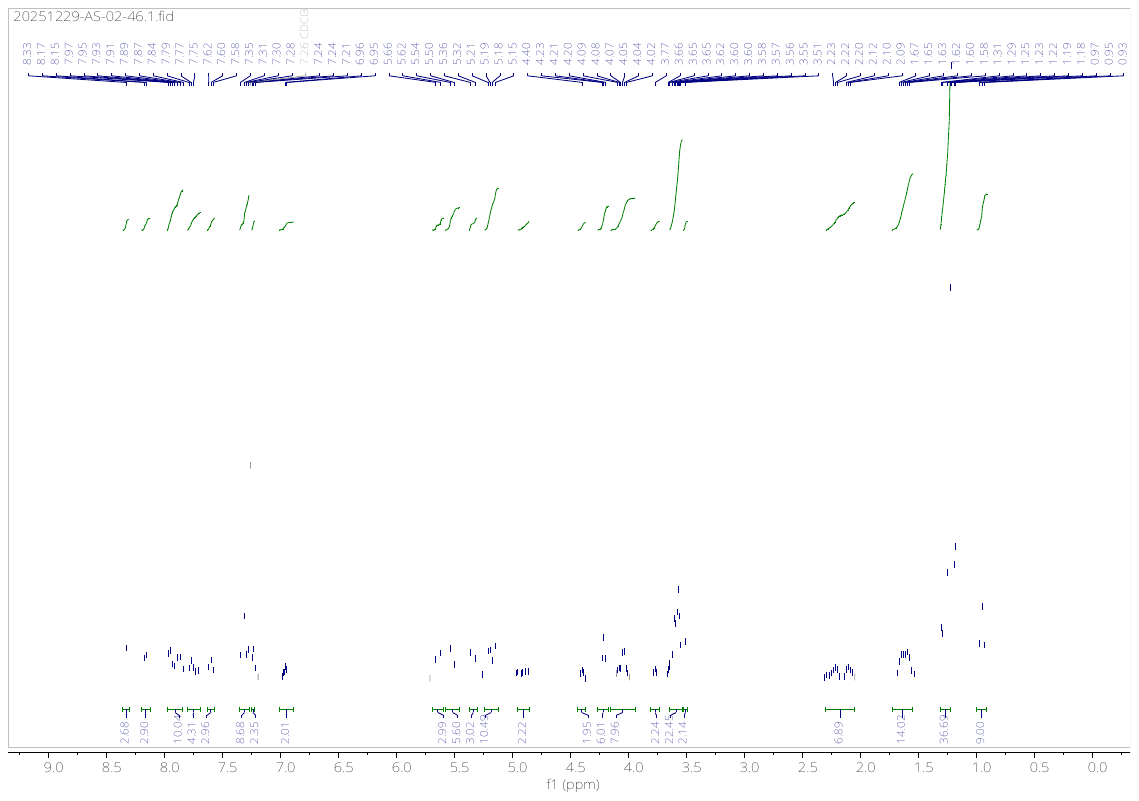
